# Self-assembly of mutant huntingtin exon-1 fragments into large complex fibrillar structures involves nucleated branching

**DOI:** 10.1101/195297

**Authors:** Anne S. Wagner, Antonio Z. Politi, Anne Ast, Kenny Bravo-Rodriguez, Katharina Baum, Alexander Buntru, Nadine U. Strempel, Lydia Brusendorf, Christian Hänig, Annett Boeddrich, Stephanie Plassmann, Konrad Klockmeier, Juan M. Ramirez-Anguita, Elsa Sanchez-Garcia, Jana Wolf, Erich E. Wanker

## Abstract

Huntingtin (HTT) fragments with extended polyglutamine (polyQ) tracts self-assemble into amyloid-like fibrillar aggregates. Elucidating the fibril formation mechanism is critical for understanding Huntington’s disease pathology and for developing novel therapeutic strategies. Here, we performed systematic experimental and theoretical studies to examine the self-assembly of an aggregation-prone N-terminal HTT exon-1 fragment with 49 glutamines (Ex1Q49). Using high resolution imaging techniques such as electron microscopy and atomic force microscopy, we show that Ex1Q49 fragments in cell-free assays spontaneously convert into large, highly complex bundles of amyloid fibrils with multiple ends and fibril branching points. Furthermore, we present experimental evidence that two nucleation mechanisms control spontaneous Ex1Q49 fibrillogenesis: (1) a relatively slow primary fibril-independent nucleation process, which involves the spontaneous formation of aggregation-competent fibrillary structures, and (2) a fast secondary fibril-dependent nucleation process, which involves nucleated branching and promotes the rapid assembly of highly complex fibril bundles with multiple ends. The proposed aggregation mechanism is supported by studies with the small molecule O4, which perturbs early events in the aggregation cascade and delays Ex1Q49 fibril assembly, comprehensive mathematical and computational modelling studies, and seeding experiments with small, preformed fibrillar Ex1Q49 aggregates that promote the assembly of amyloid fibrils. Together, our results suggest that nucleated branching *in vitro* plays a critical role in the formation of complex fibrillar HTT exon-1 aggregates with multiple ends.

## Introduction

Proteins with diverse sequences, structures and functions can spontaneously self-assemble into morphologically similar insoluble protein aggregates, often referred to as amyloid [1–3]. Amyloid aggregates are highly stable fibrillar structures with a characteristic cross-β-sheet conformation [4]. They are associated with a wide range of neurodegenerative and localized non-neuropathic diseases [5–7]. However, they also play important roles in normal cellular processes like the biogenesis of melanosomes or the regulation of translation in yeast [8, 9].

The process of fibril self-assembly can be divided into two phases: a lag phase where little or no fibrils are formed and a fibril growth phase where a large increase of fibril mass is observed [10]. Theoretical models typically describe fibrillogenesis as depending on primary nucleation, non-nucleation processes and possibly secondary processes [10–14]. The formation of an initial “nucleus” is regarded as the rate-limiting step in amyloid formation. The nucleus is currently thought to be a kinetically unstable oligomer of low abundance spontaneously forming from interacting monomers during the lag phase [12]. Once assembled, it is assumed to grow into large, highly stable β-sheet-rich structures with a fibrillary morphology by monomer addition. The characteristic fibril growth profile observed for many amyloidogenic polypeptides reflects the greater ease of addition of monomers onto existing β-sheet-rich aggregates compared to the *de novo* formation of new amyloidogenic oligomers (nuclei) from monomers through primary nucleation [14]. Several recent studies have demonstrated the involvement of secondary processes such as fragmentation [15, 16], secondary nucleation, i.e. nucleation at the surface of an existing fibril [17–19] or fibril branching [20] in the aggregation of amyloidogenic polypeptides, and have shown that theoretical models with a single nucleation mechanism cannot fully explain the aggregation process under different conditions (e.g. [15, 17–19, 21–23]).

Evidence has been provided that N-terminal huntingtin exon-1 (HTTex1) fragments with pathogenic polyglutamine (polyQ) tracts spontaneously self-assemble into insoluble aggregates with fibrillar morphology [24–30]. This indicates that formation of amyloid-like HTTex1 aggregates is mechanistically similar to α-synuclein or amyloid-β aggregation observed in various model systems [26]. Spontaneous HTTex1 fibrillogenesis critically depends on the length of the polyQ tract as well as on protein concentration and time [25]. It can be stimulated by preformed fibrils, hinting that it is dominated by a nucleation-dependent aggregation mechanism [25]. Based on studies with simple polyQ peptides [31] it was initially assumed that a single, fibril-independent primary nucleation step is critical for HTTex1 polymerization *in vitro* and *in vivo*. However, studies with full-length and C-terminally truncated HTTex1 fragments provided experimental evidence that the aggregation mechanism is indeed much more complex, involving the formation of non-fibrillar and fibrillar oligomers followed by the assembly of large fibrillary structures [27, 32–38]. Previous investigations indicate that HTTex1 fragments are produced by abnormal splicing in HD patients and transgenic animals [39], supporting their potential relevance in disease. A better understanding of the assembly cascade of HTTex1 fragments with pathogenic polyQ tracts therefore might provide important insights into pathogenesis.

In the present study, we have systematically analysed the aggregation mechanism of a disease-relevant HTTex1 fragment with 49 glutamines (Ex1Q49), using biochemical, biophysical and computational methods. We show that self-assembly of Ex1Q49 fragments released by proteolytic cleavage from soluble GST fusions results in the formation of large, highly complex bundles of Ex1Q49 fibrils with multiple ends. Furthermore, we present evidence that Ex1Q49 aggregation is controlled by a double nucleation mechanism. Initially, a primary fibril-independent nucleation process leading to the formation of aggregation-competent structures dominates Ex1Q49 aggregation. It is followed by a rapid secondary fibril-dependent nucleation process, which involves in particular the growth of new branches from the surface of existing fibrils which remain attached to the parent fibril; we denote this process nucleated fibril branching. Perturbation studies with the small molecule O4 that directly targets Ex1Q49 polypeptides and mathematical modelling of time-resolved Ex1Q49 aggregation data support our experimental results that the observed fibrillar Ex1Q49 bundles grow by a nucleated branching mechanism. Furthermore, seeding experiments with small preformed Ex1Q49 fibrils substantiate the view that nucleated branching, instead of lateral aggregation of fibrils, is the reason for the observed formation of large, complex fibril bundles. The implications of these mechanistic investigations for the pathogenesis of HD are discussed.

## Results

### Ex1Q49 spontaneously self-assembles into large, SDS-stable protein aggregates in cell-free assays

To study the kinetics of polyQ-mediated HTT exon-1 (HTTex1) aggregation and the nature of the nucleating aggregate species, a cell-free aggregation assay was established which allows the systematic investigation of aggregation reactions in a time- and concentration-dependent manner under non-denaturing conditions [40]. A purified, soluble glutathione S-transferase (GST) HTT exon-1 fusion protein with 49 glutamines (GST-Ex1Q49) was produced and purified from *E. coli* protein extracts as the starting material for the investigation of spontaneous Ex1Q49 aggregation *in vitro* (Fig. 1a). It migrates at ∼60 kDa in denaturating SDS gels (Fig. S1a). Analysis by gel filtration under non-denaturating conditions, however, revealed that the fusion protein is a stable, soluble oligomer with a size of ∼500 kDa (Fig. S1b).

**Figure 1.**
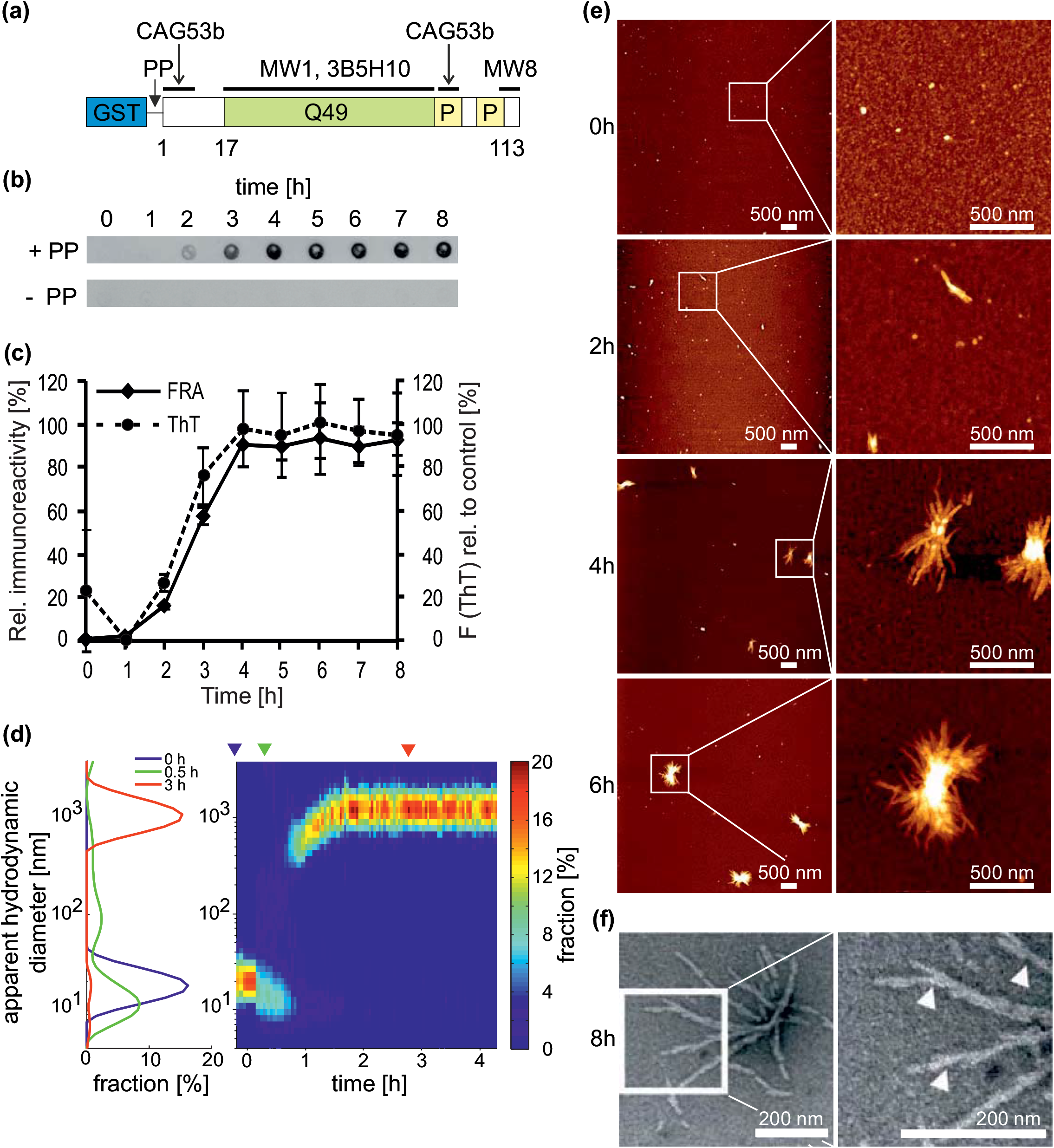
Spontaneous formation of large SDS-resistant Ex1Q49 aggregates *in vitro*. (a) Schematic representation of GST-Ex1Q49 fusion protein and epitope binding sites of monoclonal and polyclonal antibodies used. (b) Time-resolved analysis of SDS-resistant Ex1Q49 aggregates by FRA. Ex1Q49 aggregation was initiated by proteolytic cleavage of GST-Ex1Q49 fusion protein (2 µM) with PP. Aggregates retained on filter membranes were detected using the polyclonal anti-HTT antibody CAG53b. (c) Quantification of spontaneously formed Ex1Q49 aggregates by FRAs and ThT assays (solid line, mean ± SD, *n* = 4; dashed line, mean ± SD, *n* = 3). 2 µM GST-Ex1Q49 fusion protein was incubated with PP at 20°C. The highest signals were set to 100%. (d) Analysis of Ex1Q49 aggregation (20.9 µM) by dynamic light scattering. Left panel: particle size distribution prior (0 h) and after PP addition (at 0.5 and 3 h). Right panel: color-coded particle diameter distribution over time. Shown are the results for one representative experiment out of four. (e) Analysis of GST-Ex1Q49 (2 μM) aggregation reactions by AFM at different time points after addition of PP. (f) Analysis of Ex1Q49 fibril bundles by TEM revealed potential fibril branching points, which are indicated by triangles. Ex1Q49 aggregates were examined 8 h after addition of PP to GST-Ex1Q49 fusion protein (2 μM). Samples were incubated at 20°C.

Next, the purified GST-Ex1Q49 fusion protein (2 μM) was treated with the sequence-specific PreScission protease (PP) in order to release aggregation-prone Ex1Q49 fragments (Fig. 1a and S1c). Samples were incubated at 20°C in microtiter plates and the formation of stable Ex1Q49 protein aggregates was monitored over time by a denaturating membrane filter retardation assay (FRA) using the anti-HTT antibody CAG53b. This antibody specifically recognizes amino acids in the conserved N17 domain as well as the proline-rich region in HTTex1 (Fig. 1a and S1d). Previous studies revealed that denaturating FRAs can be applied to specifically quantify the abundance of SDS-stable, fibrillar HTTex1 aggregates with lengths >0.2 μm in complex protein solutions, while monomers and small oligomeric structures are not detected [25]. We found that PP treatment disintegrates soluble GST-Ex1Q49 oligomers and results in the rapid formation of SDS- and heat-stable Ex1Q49 protein aggregates, which are detectable after a short lag phase of ∼2 h (Fig. 1b and c). This confirms previous observations that HTTex1 fragments with pathogenic polyQ tracts rapidly self-assemble into highly stable protein aggregates, once they are released from GST fusion proteins by proteolytic cleavage [25, 41]. As expected, no SDS-stable Ex1Q49 aggregates were detected by FRAs in the absence of PP (Fig. 1b). The cleavage of the GST-Ex1Q49 fusion protein with PP was confirmed by SDS-PAGE and immunoblotting (Fig. S1e).

As previous studies indicate that insoluble polyQ protein aggregates are β-sheet-rich structures [42], we examined the formation of Ex1Q49 aggregates using a Thioflavin T (ThT) dye-binding assay commonly used to quantify cross-β-sheet amyloid [43]. We found that aggregates detected by FRAs and ThT-reactive aggregates form at similar rates (Fig. 1c), indicating that the observed aggregates by FRAs are β-sheet-rich structures.

We also addressed the question of whether the kinetics of spontaneous Ex1Q49 aggregation in cell-free assays is influenced by agitation. Previous investigations have demonstrated that shaking of samples with amyloidogenic polypeptides stimulates amyloid growth due to fibril breakage and growth at the ends of newly formed fibrils [15, 16], suggesting that also the aggregation of Ex1Q49 might be influenced by mechanical forces. Aggregation reactions with GST-Ex1Q49 fusion protein and PP were systematically analysed with (300 and 600 rpm) and without agitation using FRAs. We found that shaking of samples does not promote spontaneous Ex1Q49 fibril assembly *in vitro* (Fig. S2a). In particular, we did not observe an impairment of Ex1Q49 aggregation without agitation. The kinetics of the spontaneous process remained also unaltered when increasing concentrations of additional proteins such as bovine serum albumin (BSA) were added to the reactions (Fig. S2b), indicating that this process is very robust and cannot be easily influenced.

Finally, we examined the kinetics of spontaneous Ex1Q49 aggregation by dynamic light scattering (DLS), which provides size distribution profiles of aggregating protein molecules [44, 45]. For DLS analysis, a higher concentration of GST-Ex1Q49 fusion protein (20.9 μM) was needed than for FRA analysis (2 μM, Fig. 1b and c). From the autocorrelation curves, the size distribution was computed (Fig. 1d). Proteolytic cleavage of GST-Ex1Q49 oligomers with a diameter of 19.27±1.28 nm (*n* = 4) resulted in the release of smaller particles (GST and Ex1Q49 molecules) with a size distribution that peaked at 10.89±0.92 nm (*n* = 4). This is larger than the previously reported radii for GST in solution (4.2 nm, hydrodynamic radius) and Ex1Q49 (∼2.5 nm, radius of gyration), respectively [46, 47]. GST has been shown to dimerize in solution [47]. The observed diameter could therefore be explained from a mixed population of GST and Ex1Q49 oligomers. After a lag phase of ∼1 h, large particles (diameter >700 nm) rapidly started to appear, indicating the spontaneous formation of Ex1Q49 aggregates. After 2 h the size distribution reached a steady state and peaked at 1,204±93 nm (*n* = 4). Thus, our DLS studies confirm the results obtained by FRAs (Fig. 1c) and demonstrate that large Ex1Q49 aggregates are rapidly formed from monomers or very small oligomers that are released from larger GST-Ex1Q49 oligomers by proteolytic cleavage (Fig. 1d).

### Ex1Q49 spontaneously forms bundles of highly connected amyloid fibrils with multiple ends

Our studies indicate that Ex1Q49 fragments rapidly self-assemble into large, highly stable protein aggregates after a short lag phase (Fig. 1b-d). To investigate the morphology of these structures, we analysed the time-dependent formation of Ex1Q49 aggregates by atomic force microscopy (AFM). We observed that soluble, uncleaved GST-Ex1Q49 oligomers are very small spherical particles (Fig. 1e), confirming previous reports [41, 48]. However, in PP treated samples bundles of Ex1Q49 fibrils with multiple ends were detectable. These structures increased in their height, width and length over time (Figs 1e, S3a and b). This suggests that β-sheet-rich, amyloidogenic Ex1Q49 aggregates grow *in vitro* by fibril branching, a secondary process that was previously described for the peptide hormone glucagon [20].

To substantiate these observations, aggregation reactions with GST-Ex1Q49 and PP were systematically analysed by transmission electron microscopy (TEM). These studies confirmed that Ex1Q49 molecules spontaneously self-assemble into complex fibrillar bundles with multiple ends (Fig. 1f). Strikingly, the analysis by EM also revealed multiple branching points in Ex1Q49 fibrils (see white triangles in Fig 1f and red arrows in S3c), suggesting that the observed fibril ends in bundles could be formed by nucleated fibril branching rather than a lateral association of initially formed fibrils.

### Epitope-specific antibodies reveal conformational changes in soluble aggregation-prone Ex1Q49 molecules

To study fibril assembly of Ex1Q49 molecules over time under non-denaturating conditions, dot blot assays (DBAs) with epitope-specific anti-HTT antibodies were performed [49]. We initially examined the time-dependent aggregation of Ex1Q49 using the anti-polyQ antibodies MW1 and 3B5H10 (Fig. 1a), both of which recognize short polyQ epitopes in soluble HTT proteins [50–52]. We found that both antibodies readily recognize uncleaved GST-Ex1Q49 fusion protein (Fig. 2a and b), indicating that linear polyQ epitopes are exposed on the surface of GST-Ex1Q49 oligomers (Fig. S1b). Proteolytic cleavage of GST-Ex1Q49 with PP, however, caused a time-dependent decrease of antibody binding to the polyQ tract in Ex1Q49 fragments (Fig. 2a and b), indicating that the polyQ epitopes are less accessible for antibody binding in released Ex1Q49 fragments. We observed a more rapid loss of immunoreactivity with the MW1 than with the 3B5H10 antibody, indicating that the MW1-reactive polyQ conformation is accessible for antibody binding for a shorter time. We found that the rate of GST-Ex1Q49 fusion protein cleavage detected by SDS-PAGE and immunoblotting was similar to the rate of MW1-immunoreactivity loss observed by DBAs (Fig. S4a), indicating that proteolytic cleavage of the GST fusion protein is responsible for the time-dependent decrease in MW1 immunoreactivity. Thus, our studies with DBAs suggest that Ex1Q49 fragments, when released from GST fusions by proteolytic cleavage, undergo a rapid conformational rearrangement, which prevents MW1 antibody binding. This conformational change is likely to occur in soluble Ex1Q49 molecules and precedes spontaneous amyloid polymerization.

**Figure 2.**
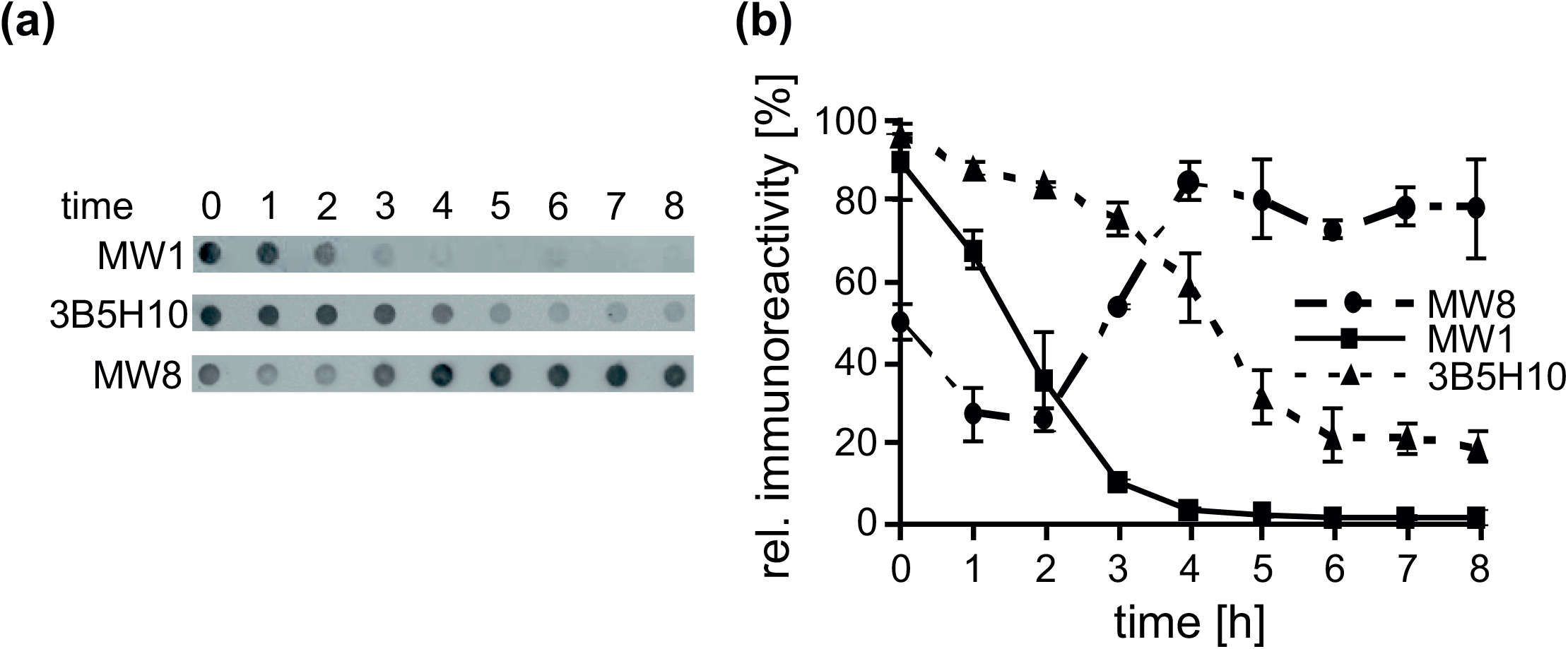
Investigation of Ex1Q49 aggregation under non-denaturing conditions using epitope-specific antibodies. (a) Analysis of spontaneous Ex1Q49 (2 μM) aggregation by non-denaturing DBAs. Samples were spotted onto a nitrocellulose membrane; retained Ex1Q49 protein was detected using the epitope-specific antibodies MW1, 3B5H10 or MW8. (b) Quantification of antibody immunoreactivity. The highest signals were set to 100%.

In comparison to the time-dependent decrease in MW1 immunoreactivity, the decrease of 3B5H10 immunoreactivity was shifted to later time points (Fig. 2a and b). This suggests that this antibody binds to monomers or small, soluble Ex1Q49 oligomers but not to large β-sheet-rich amyloid fibrils that appear later in aggregation reactions (Fig. 1c-f). To address this question in more detail, preformed fibrillar Ex1Q49 aggregates were also systematically analysed with DBAs in independent experiments (Fig. S4b). We found that the 3B5H10 antibody did not recognize these structures, confirming our hypothesis that it preferentially binds to soluble, non-fibrillar Ex1Q49 structures.

Using DBAs, we finally examined the binding of the epitope-specific antibody MW8 to aggregating Ex1Q49 molecules. This antibody recognizes a C-terminal proline-rich region in the Ex1Q49 fragment (Fig. 1a). We found that MW8 weakly binds to uncleaved GST-Ex1Q49 fusion protein in DBAs. Treatment of the fusion protein with PP further decreased MW8 immunoreactivity (Fig. 2a and b), suggesting that the epitope is less accessible for antibody binding in released, aggregation-competent Ex1Q49 molecules. However, it is also possible that the time-dependent decrease in immunoreactivity is due to an antibody avidity effect, when the spherical GST-Ex1Q49 oligomers are disintegrated upon PP treatment. After a lag phase of ∼2 h, however, a steep increase of MW8 immunoreactivity was observed (Fig. 2b), indicating that the antibody efficiently binds to spontaneously formed fibrillar Ex1Q49 aggregates (Fig. 1e). A similar result was observed when Ex1Q49 aggregation reactions were analysed in a time-dependent manner with denaturating FRAs that specifically detect large SDS-stable aggregates (Fig. S4c), confirming that the MW8 antibody preferentially recognizes β-sheet-rich, fibrillar structures in complex aggregation reactions.

### The small molecule O4 delays spontaneous Ex1Q49 polymerization in cell-free assays

To study the mechanism of Ex1Q49 aggregation and to perturb specific microscopic steps in the protein assembly process, we searched for small molecules that can influence amyloid aggregation. We selected four previously reported amyloid-binding compounds, namely curcumin (Curc), methylene blue (MB), PGL-34 and O4 (Table S1) and systematically tested their activity in the established Ex1Q49 aggregation assay. We found that only the compound O4 affected Ex1Q49 aggregation in a concentration-dependent manner (Figs 3a, 3b and S5a-c), indicating that it directly targets aggregation-prone Ex1Q49 molecules. SDS-PAGE and immunoblotting confirmed that the effect of O4 on Ex1Q49 aggregation is not due to inhibition of PP cleavage (Fig. S5d).

**Figure 3.**
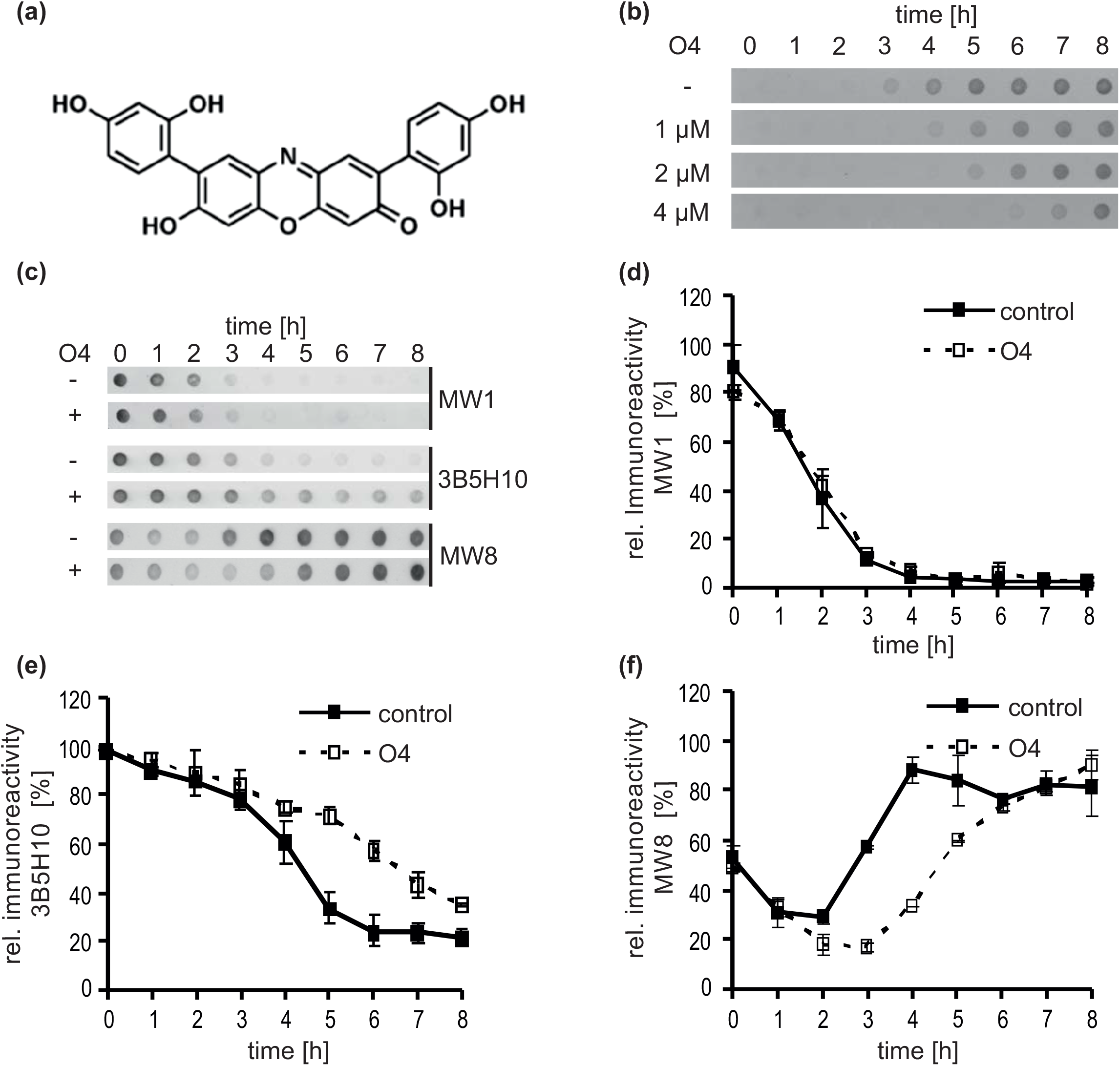
The small molecule O4 extends the lag phase in the Ex1Q49 aggregation cascade. (a) Chemical structure of O4. (b) Analysis of spontaneous Ex1Q49 aggregation in the presence and absence of O4 by FRA. GST-Ex1Q49 fusion protein (2 µM) was incubated with PP and different concentrations of O4 at 20°C. SDS- and heat-stable aggregates retained on filter membranes were immunodetected with CAG53b antibody. (c) Analysis of Ex1Q49 aggregation reactions with DBAs using the monoclonal antibodies MW1, 3B5H10 and MW8. GST-Ex1Q49 (2 µM) fusion protein was incubated with PP and O4 (2 µM) at 20°C. (d-f) Quantification of dot blot results shown in (c).

More detailed investigations revealed that O4 significantly extends the lag phase of Ex1Q49 aggregation in a concentration-dependent manner (Fig. S5e), but does not significantly affect the maximal fibril growth rate. A similar effect of the compound was observed with DLS (Fig. S5f), indicating that O4 perturbs early rather than late events in the Ex1Q49 aggregation cascade. Furthermore, studies by AFM and TEM revealed that in the presence of O4 typical fibril bundles with multiple ends are detectable (Fig. S5g), indicating that the compound does not alter the putative branching process or impair Ex1Q49 fibril growth.

We also examined the effect of O4 on spontaneous Ex1Q49 aggregation using DBAs (Fig. 3c-f). We observed that the time-dependent decrease of 3B5H10 immunoreactivity is significantly delayed upon compound treatment, while no compound effect was observed on the decrease of MW1 immunoreactivity. This suggests that O4 preferentially targets soluble, 3B5H10-reactive Ex1Q49 molecules that are released from GST fusions and delays their incorporation into fibrillar aggregates. Strikingly, a delay in the increase of MW8 immunoreactivity was also observed (Fig. 3f), suggesting a causal relationship between disappearance of 3B5H10- and appearance of MW8-reactive Ex1Q49 molecules in spontaneous aggregation reactions. Together, these results support the data obtained by FRAs and DLS (Figs 3b and S5f).

### Molecular dynamic simulations indicate that O4 directly targets HTTex1 molecules with expanded polyQ tracts and alters their conformation

We employed computational approaches for an assessment of how O4 might interact with aggregation-prone HTTex1 molecules, using two *in silico* generated structures: i) the most representative conformation of a β-sheet-rich HTT exon-1 protein with 47 glutamines (Ex1Q47) [53] and ii) a structure of Ex1Q46 with lower β-sheet content from a different study [54]. These proteins are similar in size to the Ex1Q49 fragment that was used in our experimental studies.

We first performed systematic docking calculations for Ex1Q47 with AutodockVina using both rigid and flexible approaches and different grid sizes [55]. We then selected seven possible binding sites, based on their geometrical differences and their calculated docking affinities, which cover a range from −7.6 to −10.4 kcal/mol (Table S2). To better discriminate between the docking conformations and to take explicitly into account solvent and dynamics effects, we performed Quantum Mechanics/Molecular Mechanics Molecular Dynamics (QM/MM MD) simulations (see Materials and Methods for details) with these seven binding sites. The analysis of the QM energies from the QM/MM MDs indicates that O4 binding to positions A7 and D1 is more favourable, followed by A5 and A8 (Table S2). Therefore, these four binding sites were used for further computational studies. Contacts between O4 and the polyQ region were found for all binding positions, including those less favoured, which is not surprising given the size of this region. Interestingly, the interaction motifs of O4 with Ex1Q47 on the preferred binding sites A7 and D1 suggested that the ligand associates not only with the β-sheet-rich polyQ tract but also with other β-sheet regions at the N-terminal 17 amino acids in Ex1Q47 as well as the C-terminus (Fig. 4a and Table S2). The N-terminal residues in HTTex1 were previously shown to promote protein aggregation in cell-free assays [28, 56], substantiating our hypothesis that direct binding of O4 to soluble, aggregation-prone Ex1Q49 molecules might slow down the formation of fibrillar aggregates (Fig. 3c-f).

**Figure 4.**
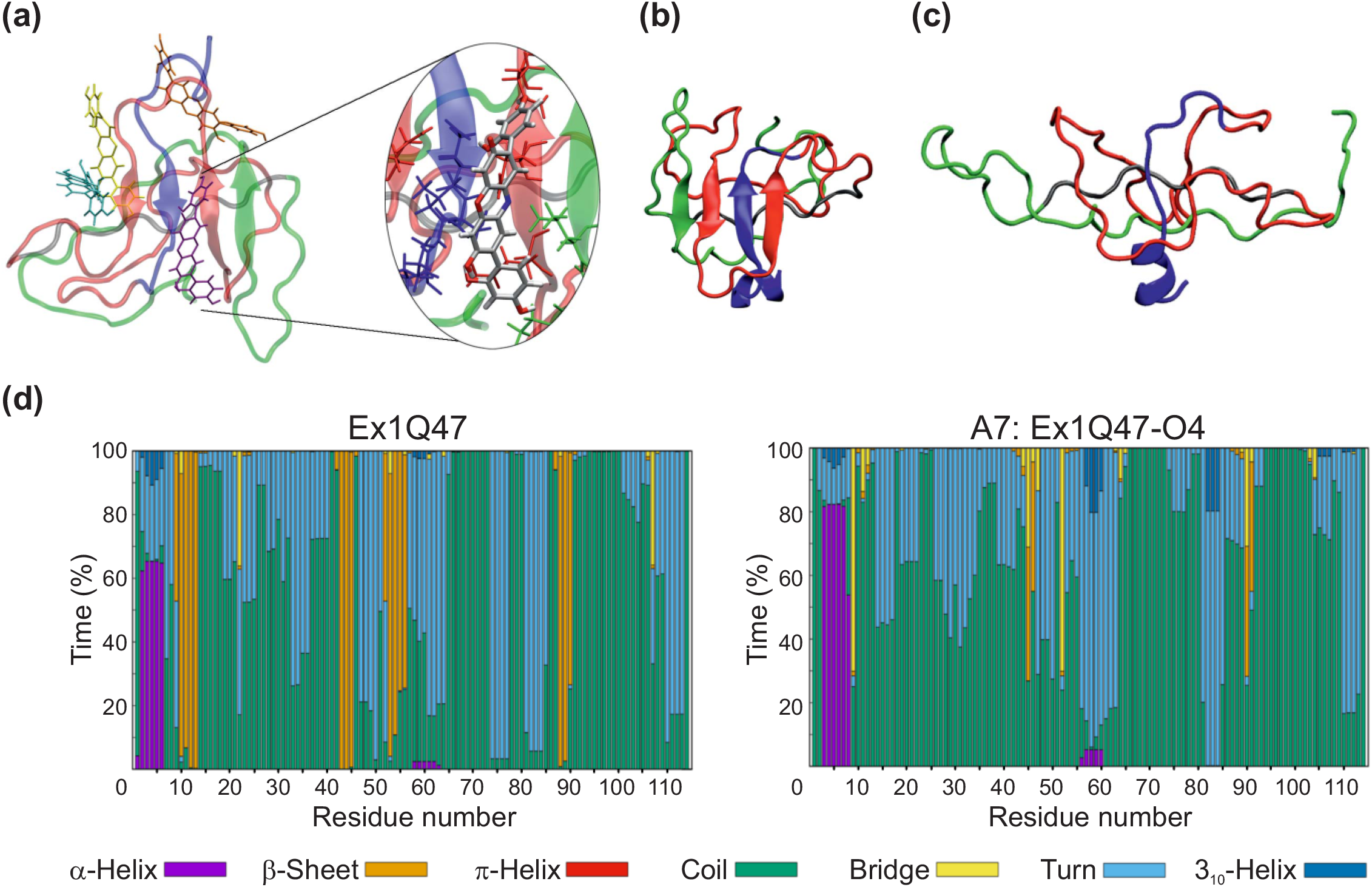
The compound O4 perturbs the structure of an N-terminal, β-sheet-rich Ex1Q47 fragment. (a) Computational docking and QM/MM MD studies predict that O4 binds to β-sheet-rich Ex1Q47 molecules. O4 is colored according to the binding motif: A7 purple, D1 orange, A5 yellow and A8 cyan (see also Fig. S6). Different colors are used for the Ex1Q47 regions: N-terminus in blue, polyQ in red, polyP in gray and the mixed region including the C-terminus is shown in green. One of the most favoured binding motifs, A7, is displayed in more detail. The residues within a 3 Å distance between O4 and Ex1Q47 are shown. (b,c) O4 alters the β-sheet structure of Ex1Q47 monomers. Representative most populated structures of Ex1Q47 (MD simulations); (b) reference system without O4 and (c) Ex1Q47 in the Ex1Q47-O4 system (binding site A7, O4 omitted). (d) Secondary structure propensity per residue for the MD simulations of Ex1Q47 without (left) and with (right, binding site A7) O4. The decrease in β-sheet content in Ex1Q47 due to O4 binding is particularly evident in residues 9 – 13 and 52 – 56.

Next, to examine whether O4 binding influences the β-sheet content in Ex1Q47, we performed additional molecular dynamics (MD) simulations, focussing in this case, on four putative compound bindings sites (e.g., A7, D1, A5 and A8) in the Ex1Q47 protein (Table S2). These simulations indicate that O4 binding promotes a time-dependent disassembly of the initially available β-sheet structure in Ex1Q47 (Figs 4b-d, S6 and Movies S1 and S2). For instance, the β-sheet content in Ex1Q47 was found to be 12% in simulations without O4 (Table S3). With the only exception of A8, where it remains constant, this value decreases to 9, 7 and 2% for A5, D1 and A7 respectively (Table S3 and Fig. S8). In the absence of O4, the initial four β-strands remain unchanged during the entire simulation (Fig 4d). In comparison, an almost complete loss of the four β-strands was observed when the binding site A7 was analysed in the presence of O4 (Figs 4d and S6). Similarly, the β-strand covering residues 9 – 13 and 52 – 56 in Ex1Q47 were lost with the compound in the context of the binding site D1 (Fig. S8). In the case of A5, the β-strands are not completely lost in the presence of O4 but residues 9 – 13, 52 – 56 and 87 – 91 explore coil and turn conformations for more than half of the simulations time (Fig. S8).

To investigate whether the same compound effects can also be observed with pathogenic HTTex1 conformations having a lower β-sheet content, we performed two additional independent simulations with an Ex1Q46 structure, either complexed with or without O4. The total combined simulated time in this study reached 1.6 microseconds. The MD simulations performed with Ex1Q47 suggested that, after O4 destroys the β-sheets, it starts exploring the surface of Ex1Q47 without any preferred binding site (Fig. S10). Therefore, and due to the relatively low β-sheet content of the Ex1Q46 structure, we used an initial geometry in which O4 was placed within 4 angstroms of the surface-exposed side of the β-strands of the Ex1Q46 conformer. As with Ex1Q47, the simulations with Ex1Q46 suggested that O4 binding promotes a time-dependent disassembly of the initial β-sheet structure (Figs S7 and S9). For Ex1Q46, the β-sheet propensity decreases by 3% in the presence of O4 in both simulations (Table S3).

The inter-residue contact maps also evidence a decrease in the β-sheet content in both Ex1Q47 and Ex1Q46. This can be seen in the patterns for the interactions between residues 9-12 and 42-45 of Ex1Q47 (Fig. S12) and residues 24 – 33 of Ex1Q46 (Fig. S13). Beyond the β-sheet region, O4 also impacts the structures of both proteins by promoting more open conformations with less inter residue contacts. For example, the interactions between residues 70 – 80 and 85 – 100 are weaker in A5, A7, A8 and D1 with respect to Ex1Q47 without ligand (Fig. S12). Since the initial structures of Ex1Q46 are highly disordered, the contact maps for this protein are less revealing than for Ex1Q47. Nevertheless, for the MD simulations with Ex1Q46 the impact of O4 is evident by the loss of the interactions between residues 32 – 40 and 52 – 60 (Fig. S13).

Taken together, the simulations suggest that O4 promotes a binding-dependent destruction of the β-strands in Ex1Q47 and Ex1Q46. Interestingly, O4 does this without having a conserved binding site in the HTTex1 protein. Thus, after the β-strands are destroyed, the binding of O4 to HTTex1 becomes weak and O4 starts exploring the protein surface (Fig. S11). This potentially might impair the formation of newly formed β-sheet-rich structures in Ex1Q49 molecules, which are released in our experimental studies by proteolytic cleavage of GST-Ex1Q49 fusion proteins (Fig. 1a-c).

### A theoretical nucleated branching model correctly reproduces the effect of O4 on spontaneous Ex1Q49 aggregation

The experimental results suggest that fibril branching could be responsible for the rapid formation of Ex1Q49 fibril bundles in cell-free assays (Fig. 1f). To better understand the molecular events underlying spontaneous Ex1Q49 fibrillogenesis, we next developed a theoretical aggregation model that, besides primary nucleation and templated polymerization as an elongation process [11, 12, 15], also includes nucleated fibril branching, a special form of secondary nucleation (see Material and Methods and Supplemental Methods, Kinetic modelling of Ex1Q49 aggregation). Previously, various theoretical models have been reported that include primary nucleation, templated fibril polymerization, and secondary processes [15, 16, 18]. However, none of these models explicitly accounts for the fate of newly nucleated branches that remain attached to mother fibrils as observed for instance for glucagon fibrils [20]. Since we did not observe an effect of agitation in our experimental studies (Fig. S2a), fragmentation was not included in our model.

The theoretical Ex1Q49 aggregation model with nucleated branching describes four processes: *i*) the release of the aggregation-prone Ex1Q49 molecules by proteolytic cleavage of GST-Ex1Q49 through the action of PP; *ii*) the initial formation of a growth nucleus via primary nucleation (Fig. 5a green arrows; size parameter *n_c_* and rate constants *k_n_*_1_ and *k_-n_*_1_,); iii) fibril elongation growth by addition of Ex1Q49 monomers via templated polymerization (Fig. 5a, red arrows, rate constant *k*_1_); and iv) secondary nucleation at the surface of Ex1Q49 fibrils, resulting in the formation of multiple interconnected growing fibril branches (Fig. 5a blue arrows, order *n*_b_ and rate constant *k_b_*). The last process is denoted as nucleated branching. Compared to previous models with secondary nucleation [21], our model assumes that each protein unit only nucleates a limited number of branches and more importantly considers the fate of the newly nucleated fibrils that remain attached to the mother fibril. The latter modification is necessary to describe the large complex bundles observed *in vitro* and the effect of perturbations on their size. Since spontaneously formed fibrillar Ex1Q49 aggregates are very stable structures (Fig. 1b and c), our theoretical model assumes irreversible reactions for templated polymerization and nucleated branching (see Materials and Methods and Supplemental Methods Kinetic modelling of Ex1Q49 aggregation). To compare the model to the experimental FRA data, we developed a normalization procedure and characterized the initial PP cleavage reaction (see Material and Methods and Fig. S14a-c). Since the FRA assay only detects Ex1Q49 fibrils when they are SDS-stable and have a length of >0.2 μm [40], we used a size threshold for aggregate detection (*C_min_*, see Supplemental Methods, Eqs. S32-S33) estimated together with the kinetic model parameters.

**Figure 5.**
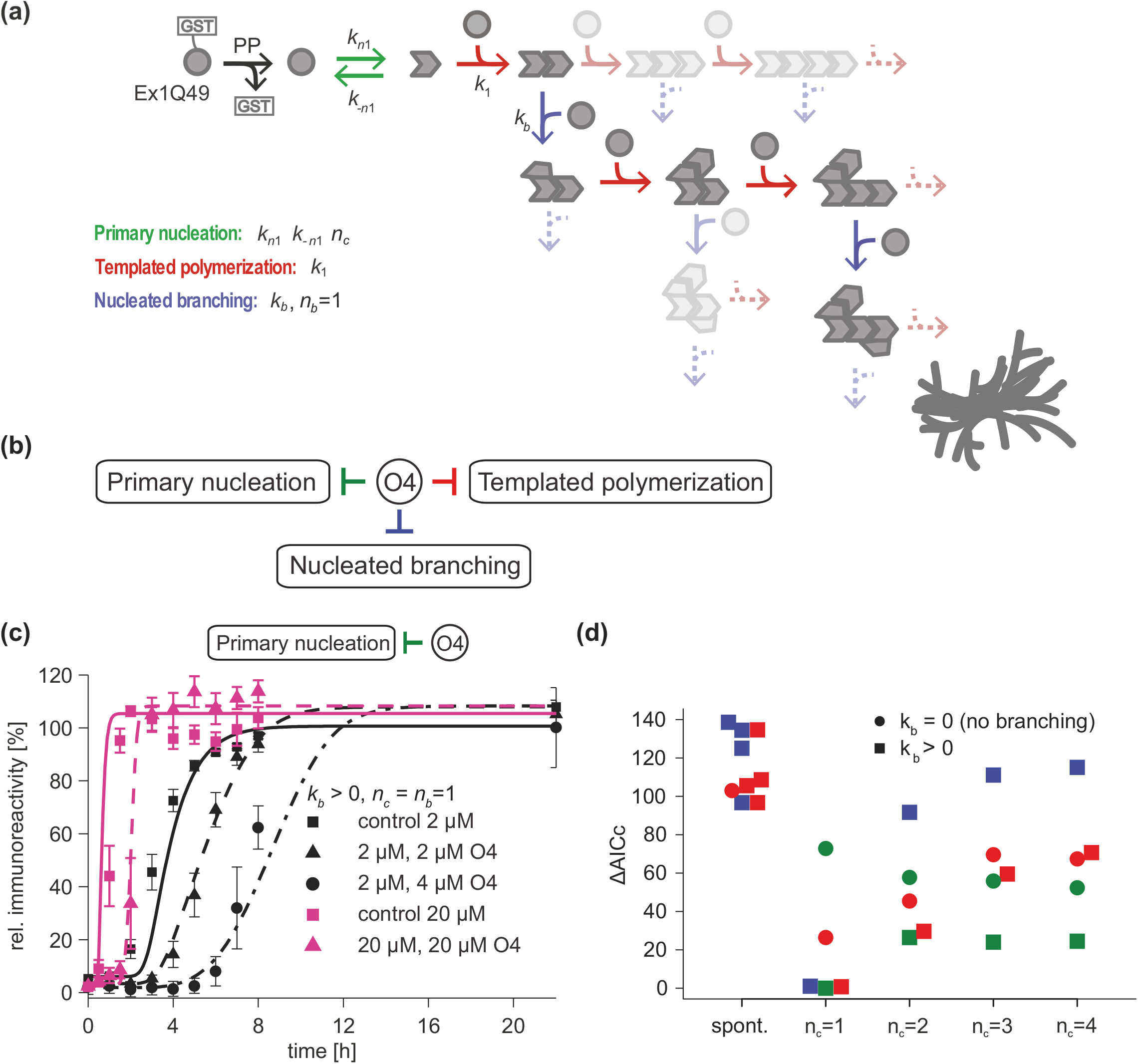
The theoretical model with nucleated branching correctly reproduces the effect of O4 on spontaneous Ex1Q49 aggregation. (a) Schematic for a nucleated branching model. After PP cleavage of GST, Ex1Q49 can spontaneously nucleate and switch to an active aggregation-prone conformation with rate constant *k_n1_*. Fibrils can then elongate by addition of monomers in templated polymerization, rate constant *k_1_*. In addition, a nucleated branching reaction can occur with rate constant *k_b_*. This creates a new growing fibril. If the branching rate constant *k_b_* = 0, the model reduces to a linear templated polymerization model. (b) In the model O4 can either decrease the primary nucleation rate constant *k_n1_* (green), the nucleated branching rate constant *k_b_* (blue), or the templated polymerization rate constant *k_1_* (red) in a concentration-dependent way. (c) Simulated FRA data from the model with nucleated branching (*n_c_* = 1 and *n_b_* = 1) without (solid lines) and with O4 (dashed lines). The experimental data (symbols, mean ± std) show aggregation for 2 different initial GST-Ex1Q49 concentrations and 3 different O4 concentrations as indicated using the CAG53b antibody. Experimental data and model were normalized to maximal value of the control with 2 µM initial GST-Ex1Q49. Parameter ranges and values are listed in Tables S4 and S5. (d) Differences in the corrected Akaike information criterion (*AICc*) with respect to the best fitting model for three different O4 mediated inhibition scenarios (shown in colored symbols) and the model without nucleated branching (*k_b_* = 0, circle) or with nucleated branching (*k_b_* > 0, squares) as well as different values of the nucleus size *n_c_* and the branching order *n_b_*. Models were fitted to the experimental data from (c) (see Supplemental Data). A large value indicates a less good fit compared to the best model.

Our experimental studies indicate that O4 treatment delays the lag phase of spontaneous Ex1Q49 aggregation in a concentration-dependent manner, while it does not significantly influence the more rapid amyloid growth phase (Fig. 3b). Previous studies provide evidence that various microscopic processes like primary and secondary nucleation and even fibril growth can occur during the lag phase of spontaneous aggregation reactions [57], suggesting that O4 might interfere with different mechanisms active in this phase of aggregate formation. We next asked whether theoretical models with or without nucleated branching (Fig. 5a) can describe the observed compound effects in our experiments (Fig. 3b), when assuming that the compound inhibits either primary nucleation, templated polymerization or nucleated branching (Fig. 5b). We fitted the theoretical models that either include or lack nucleated branching to Ex1Q49 aggregation data that were obtained in the presence and absence of O4 at different initial concentrations of GST-Ex1Q49 (see Fig. S14; Material and Methods; Supplemental Methods, Fitting of HTTex1 kinetic models to data and Tables S4 and S5). Spontaneous aggregation of Ex1Q49 was measured in a time-dependent manner by denaturating FRAs using the CAG53b antibody (Figs 1b and 5c, symbols). For all simulations, we assumed that the nucleus size is similar for both primary nucleation and nucleated branching, *n_c_* = *n_b_*. Models where *n_c_* ≠ *n_b_* gave less or equally good fits. By systematically varying the nucleus size *n_c_*, we found that the experimental data is best reproduced by the model with nucleated branching and a nucleus of size *n_c_* = 1, independent of the assumed inhibiting mode of O4 (Figs 5c, lines and S15a-b, lines). The distinction was made according to the corrected Akaike information criterion (Fig. 5d, green, red and blue squares for n_c_=1). In contrast, the models without nucleated branching (*k_b_* = 0) failed to reproduce the data (Figs 5d, green and red circles and S15c). The reason is that when sufficient fibrils are formed at the end of the lag phase the positive feedback mediated by nucleated branching overcomes the effect of O4. This process is independent of the specific mechanism by which O4 inhibits aggregation. Together, these results indicate that the nucleated branching process in theoretical models (Fig. 5a) is required to describe the experimental data. Furthermore, they suggest that the perturbation of different microscopic processes (primary nucleation, nucleated branching or templated polymerization) could be responsible for the observed extension of the lag phase when O4 is added to spontaneous reactions (Fig. 3b).

Previous studies indicate that time-resolved aggregation profiles obtained at various protein concentrations are sufficient to distinguish between aggregation mechanisms when the experimental data are globally fitted to different theoretical aggregation models [15, 19]. One assumption in these studies was that the aggregate detection assay (e.g. a ThT assay, [15]) can monitor a very broad range of aggregate species including very small structures such as dimers, trimers and small oligomers as well as large fibrils consisting of thousands of molecules. However, there is no clear evidence that standard aggregate detection methods like the ThT assay are sensitive enough to monitor the very early aggregate species. The denaturating FRAs applied in our study exclusively monitor the growth of relatively large, SDS-stable Ex1Q49 aggregates [40]. Thus, as for the fitting of the models to the O4 data (Fig. 5c), an aggregate size detection threshold was applied when the theoretical aggregation models were globally fitted to aggregation profiles obtained at various GST-Ex1Q49 concentrations (Fig. S16a, symbols). Under these assumptions, the generated concentration-dependent data sets are not sufficient for the prioritization of specific aggregation mechanisms because theoretical models with and without branching both can describe concentration-dependent aggregation data equally well (Figs S16a and b, filled symbols). Under the assumption that the FRA method detects large as well as very small Ex1Q49 protein assemblies, however, the theoretical model with nucleated branching (Fig. 5a) indeed globally fits concentration-dependent aggregation profiles significantly better than the theoretical model without this process (Figs S16b and c, open symbols). This means that a detailed qualification of the microscopic processes in the aggregation cascade is not possible when theoretical models are fitted to simple concentration-dependent aggregation data. Use of a perturbator molecule like O4 provides critical experimental data that enable meaningful distinctions of aggregation mechanisms with theoretical models.

Taken together, our experimental and theoretical studies with the small molecule O4 support the hypothesis that nucleated branching shapes the aggregation kinetics of Ex1Q49 and is likely to be critical for the formation of large bundles observed by AFM and TEM (Fig. 1e and f). All critical assumptions for the theoretical modelling studies are summarized in Fig. S17.

### The theoretical nucleated branching model predicts that O4 preferentially perturbs primary nucleation in the Ex1Q49 aggregation cascade

Our MD simulation studies suggest that O4 treatment perturbs the β-sheet structure in aggregation-prone HTTex1 molecules (Fig. 4). This might impair primary nucleation, a critical early step in the amyloid polymerization cascade [57] and lead to an extension of the lag phase in Ex1Q49 aggregation reactions (Fig. 3b). Our theoretical studies suggest that addition of O4 together with PP to Ex1Q49 aggregation reactions might extend the lag phase because the compound inhibits primary nucleation, templated polymerization or nucleated branching (Figs 5c-d and S15a,b). We hypothesized that inhibition of primary nucleation with O4 should have a different effect on the length of the lag phase than inhibition of nucleated branching or templated polymerization, especially when the compound is added at later time points to aggregation reactions. Indeed, the theoretical aggregation model with nucleated branching predicts different outcomes of such an experiment depending on which microscopic process is inhibited by O4 (Fig. 6a-c). For example, if we consider an inhibition of primary nucleation by O4, the model predicts that addition of O4 after 2 h does not significantly influence the lag phase. In contrast, in the cases where nucleated branching or templated polymerization are inhibited by O4, the model predicts a significant extension of the lag phase, even when the compound is added to reactions after 2h (Fig. 6b and c). This can be explained by the fact that primary nucleation is a very early event in the aggregation cascade, whereas nucleated branching and templated polymerization occur later in the process. Thus, when primary nucleation is perturbed, compound addition after 2 h has a weak effect on the length of the lag phase as the GST cleavage (requires ∼2h, Fig. S1e) and the primary nucleation process (requires ∼10 min, Table S4) are nearly complete after 2h. However, in scenarios where the compound perturbs nucleated branching or templated polymerization, it can still extend the lag phase in Ex1Q49 aggregation reactions.

**Figure 6.**
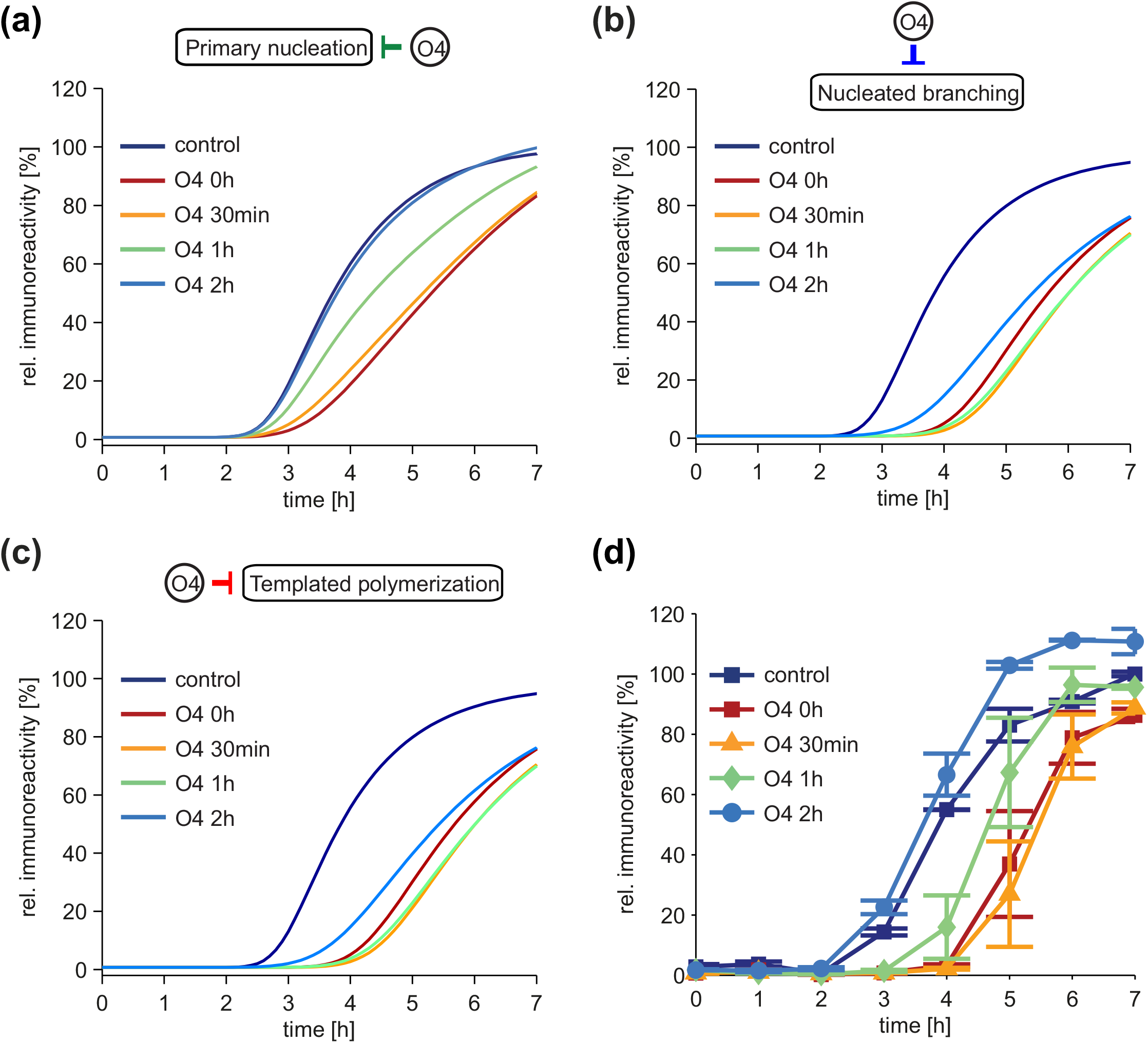
The theoretical model with nucleated branching indicates that O4 inhibits primary nucleation. (a) Simulations of the aggregation of Ex1Q49 (2 μM GST-Ex1Q49) of the best fitting model from Fig. 5c-d in which O4 is assumed to inhibit primary nucleation for addition of 2 μM O4 at different time points post PP addition as indicated. (b) Predicted kinetics for the addition of 2 μM O4 at the indicated time points to the aggregation of Ex1Q49 (2 μM GST-Ex1Q49) in the model with nucleated branching where O4 inhibits the nucleated branching process. (c) Predicted kinetics for the addition of 2 μM O4 at the indicated time points to the aggregation of Ex1Q49 (2 μM GST-Ex1Q49) in the model with nucleated branching where O4 inhibits the templated polymerization process. (d) 2 µM GST-Ex1Q49 was incubated in the absence (control) or presence of 2 µM O4 added together with PP (0 h), 30 min, 1 h, or 2 h post PP addition. Aliquots were analysed by FRAs and were immunodetected with CAG53b antibody. Data were normalized to the control value at 7 h.

Finally, we investigated experimentally whether the addition of O4 at later time points influences the lag phase in spontaneous Ex1Q49 aggregation reactions. We found that the addition of O4 after 0.5, 1 or 2 h to the fibril formation process was indeed less efficient in extending the lag phase (Fig. 6d) than the addition of the compound at t = 0 h together with PP. This supports our hypothesis that O4 targets primary nucleation events in the Ex1Q49 aggregation cascade.

### Predicting the impact of seeding on spontaneous Ex1Q49 aggregation

Preformed aggregated β-sheet-rich polypeptide structures of varying sizes can act as nuclei or seeds that potently stimulate the formation of fibrillar structures [58, 59]. When they are added to aggregation reactions at early time points, they efficiently shorten the lag phase prior to the fibril growth phase [60]. We first studied putative seeding effects of preformed Ex1Q49 fibrils (seeds) on spontaneous HTTex1 aggregate formation using our theoretical model with nucleated branching. The model predicts that addition of increasing amounts of seeds to aggregation reactions causes a decrease in the lag phase without affecting the rate of fibril formation (slope, Fig. 7a, showing the percentage of protein bound in complexes larger than the simulated seed size). Interestingly, we found that the addition of seeds to reactions also changes the predicted size distribution of Ex1Q49 aggregates. In Fig. 7b, we show the average size at steady state, normalized to the average aggregate size without seed addition (seed fraction = 0), as a function of the seed fraction (Fig. 7b, red line). Strikingly, for higher seed concentrations the size of the predicted Ex1Q49 fibrils was strongly decreased. Moreover, the model predicts that high concentrations of seeds also strongly decrease the average number of branches per aggregate (Fig. 7b, blue line). This is due to the fact that the only pathways for aggregate growth in the theoretical model are templated polymerization and nucleated branching and fibrils can only grow by monomer addition. The seeds in the reaction compete for a limited pool of monomers, which leads to the formation of small aggregates when seed concentration is high, while large aggregates should form when the concentration is low. For very low numbers of seeds, we observed a small increase in aggregate size and number of branches (Fig. 7a). The explanation for this increase is that the added seeds outcompete spontaneously formed nuclei for monomers and grow more rapidly into larger fibrils.

**Fig. 7:**
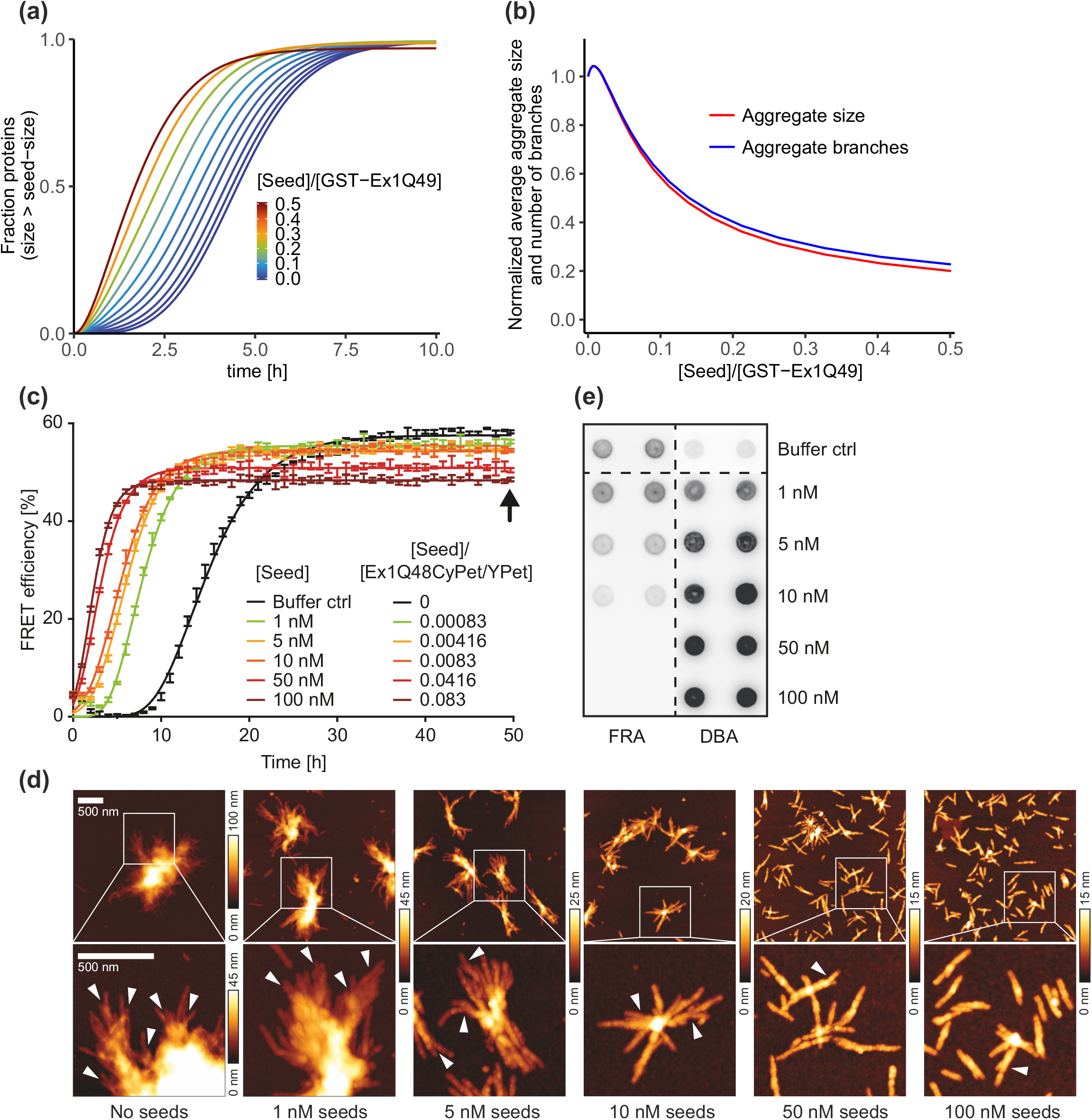
A nucleated branching process correctly predicts the outcomes of HTTex1 seeding experiments. (a) Simulation of the best fitting model from Fig. 5c-d with addition of different fractions of seeds [Seed]/[GST-Ex1Q49]. Shown is the percentage of protein bound in complexes above the seed size. In the simulations, we use dimers as seeds, this is the smallest possible seed size. (b) Average simulated amyloid size (red line) and number of branches per aggregate (blue line) at steady-state for different fractions of seeds computed from Eq. S26, Supplement. All quantities are normalized to the values without seeds. (c) Co-aggregation of the sensor proteins Ex1Q48-CyPet/-YPet was monitored in the presence of different concentrations of sonicated Ex1Q49 aggregates (seeds) by quantification of FRET for 50 h. Indicated seed concentrations are equivalent to initially applied monomer concentrations. The black arrow indicates the time point at which aggregation reactions were analysed by DBAs, FRAs and AFM shown in (d) and (e). (d) AFM analysis of GST-Ex1Q48-CyPet/-YPet co-aggregates formed in the absence and presence of different seed concentrations; samples were analysed after 50 h. Scale bar: 500 nm. (e) Ex1Q48-CyPet/-YPet co-aggregates shown in (d) were analysed by DBAs and FRAs. Proteins were detected using an anti-GFP antibody.

### Preformed Ex1Q49 seeds shorten the onset of spontaneous aggregate assembly in a concentration-dependent manner

To study the impact of preformed Ex1Q49 seeds on aggregate formation, we designed a FRET-based HTTex1 aggregation assay. We first produced and purified recombinant GST-HTTex1 fusion proteins, which are C-terminally fused to the fluorescent proteins CyPet or YPet (GST-Ex1Q48-CyPet or -YPet; Figs S18a and b). We hypothesized that these fusion proteins, when co-incubated with PP, release the aggregation-prone fragments Ex1Q48-CyPet and - YPet that over time co-assemble into ordered fibrillar aggregates and induce FRET (Fig. S18b).

First, we incubated different concentrations of the fusion proteins GST-Ex1Q48-CyPet and -YPet (1:1 molar ratio) in 384-well plates with PP and quantified the appearance of FRET over time. We observed a steep increase of FRET efficiency starting after ∼4-10 h, indicating that the proteolytically released HTTex1 fusion proteins, after a lag phase, rapidly self-assemble into ordered, high-molecular weight aggregates (Fig. S18c). The aggregation profile is similar to the untagged Ex1Q49 proteins (Fig. 1c) but the lag phase is extended due to the C-terminal fluorescent tags that slow down spontaneous aggregation.

Next, we produced well-defined, fibrillar Ex1Q49 aggregates to investigate them for their seeding activity in the FRET-based HTTex1 aggregation assay. We incubated GST-Ex1Q49 fusion protein for 24 h with PP in order to obtain large fibrillar aggregates. The generated fibrils were sonicated for 1 min on ice to break the fibrils into small fibrillar seeds. Analysis by AFM confirmed that small, fibrillar Ex1Q49 seeds were obtained, while large bundles of HTTex1 fibrils (>1 μm) were detectable in non-sonicated samples (Fig. S18d).

Finally, we assessed whether the sonicated Ex1Q49 aggregates influence the co-polymerization of the reporter proteins Ex1Q48-CyPet and –YPet in FRET assays. We treated a mixture (1:1 molar ratio) of the fusion proteins GST-Ex1Q48-CyPet/-YPet with PP and various concentrations (∼1-100 nM) of preformed sonicated Ex1Q49 seeds and analysed the co-polymerization of the released fusion proteins Ex1Q48-CyPet/-YPet by quantification of FRET. We observed a robust shortening of the lag phase when preformed Ex1Q49 seeds were added to reactions (Fig. 7c), indicating that these structures stimulate Ex1Q48-CyPet/-YPet co-polymerization in a concentration-dependent manner. With the highest seed concentrations, the maximum FRET efficiency was reached after ∼4-5 h, with the lowest seed concentrations after ∼15 h. This supports our computational prediction for the untagged Ex1Q49 protein that addition of preformed HTTex1 seeds shortens the lag phase in amyloid formation reactions (Fig. 7a).

### High concentrations of preformed Ex1Q49 seeds cause the formation of small fibrillar HTTex1 aggregates

Our theoretical investigations suggest that large numbers of small, largely unbranched HTTex1 fibrils should be obtained when the spontaneous polymerization reactions are treated with very high concentrations of preformed seeds (Fig. 7b). To validate this prediction experimentally, we collected samples from both seeded and non-seeded aggregation reactions after 50 h (Fig. 7c, black arrow) and systematically analysed them by AFM. We observed few, very large bundles of HTTex1 fibrils in reactions that were treated with low (1 nM) seed concentrations, while larger numbers of smaller fibrils were observed with higher seed concentrations (5-50 nM) (Fig. 7d). Interestingly, a large number of very small HTTex1 fibrils with few or no branches were detected when samples with very high seed concentrations (100 nM) were analysed, confirming our theoretical predictions (Fig. 7b).

Together, these studies also indicate that the complex bundles of HTTex1 fibrils observed in spontaneous reactions (Fig. 1e) are indeed formed from small seeds/nuclei through monomer addition rather than the lateral association of fibrils. The analysis of seeded aggregation reactions by FRET shows that at high seed concentrations (10-100 nM) the formation of HTTex1 fibrils is completed after 5-10 h (Fig. 7c). As large numbers of small fibrils are detected by AFM after 50 h in seeded reactions (Fig. 7d), we assume that the small fibrils, which are already formed after 5-10 h, have a very low tendency to co-aggregate into large complex structures. In this way, our seeding experiments strongly support a nucleated branching mechanism in HTTex1 aggregation. Under the assumption that HTTex1 fibrillogenesis is dominated by a lateral association of small fibrils, we would expect that an incubation period of 50 h with high seed concentrations should lead to the assembly of large fibril bundles (Fig. S19).

Finally, we analysed all aggregation reactions after 50 h with FRAs, which exclusively detect large (>0.2 μm), SDS-stable fibrillar HTTex1 aggregates retained on cellulose acetate membranes [40]. In reactions treated with high concentrations of Ex1Q49 seeds, no Ex1Q48-CyPet/-YPet aggregates were detectable on filter membranes (Fig. 7e), while they were readily observed in untreated reactions and reactions treated with low seed concentrations. This confirms the initial hypothesis that addition of high seed concentrations to aggregation reactions leads to the formation of many small HTTex1 aggregates (Fig. 7b). In reactions treated with high seed concentrations, however, anti-HTT immunoreactivity was detectable with DBAs (Fig. 7e), which detect both soluble and aggregated HTTex1 proteins independent of their size. Together, these investigations support the observations made by AFM (Fig. 7d) that addition of high concentrations of preformed Ex1Q49 seeds to the Ex1Q48-CyPet/-YPet aggregation reactions leads to the formation of small fibrillar structures, whereas large, stable fibrils are obtained when low concentrations of seeds are added to reactions.

## Discussion

In this study, we systematically investigated the aggregation mechanism of a pathogenic N-terminal HTTex1 fragment released from a soluble recombinant GST-Ex1Q49 fusion protein by site-specific proteolytic cleavage using a combination of experimental and computational methods. We were able to show that Ex1Q49 molecules *in vitro* rapidly self-assemble into very stable complex fibril bundles with multiple ends and branching points. Furthermore, we obtained experimental evidence that two nucleation processes (primary nucleation and nucleated branching) and fibril elongation characterize Ex1Q49 aggregation. An earlier study already showed the importance of primary nucleation for the aggregation of polyQ peptides [31]. Here, we present experimental and theoretical results that three microscopic processes and in particular nucleated fibril branching play a critical role in the spontaneous assembly of Ex1Q49 aggregates.

To study the importance of nucleated branching in the Ex1Q49 aggregation cascade, we developed a theoretical aggregation model that includes primary nucleation, nucleated branching and templated polymerization. Nucleated branching is a special case of secondary nucleation, where new branches form on the surface of already formed fibrils and remain attached to them. This auto-catalytic pathway generates new extensions with a rate proportional to the amount of existing aggregates (see e.g. [12, 18, 21–23, 61]). Compared to previous theoretical models (e.g. [14, 16, 17, 21]), we specifically account for the formation of new fibril ends by a nucleated branching mechanism. This feature is essential to describe the branched structures observed *in vitro* (Fig. 1e) and the effect of seeds on the aggregate size (Fig. 7b). We use multiple approaches, such as chemical inhibitors and seeding, to identify via global fitting the underlying aggregation mechanism (for a summary of our overall strategy see Fig S.17). We found that time-resolved Ex1Q49 aggregation profiles, generated in the presence and absence of the aggregation-inhibiting small molecule O4, are accounted for by the nucleated branching model. Furthermore, this model correctly predicts the impact of a delayed addition of O4 on Ex1Q49 aggregation (Fig. 6) as well as of preformed Ex1Q49 seeds, which stimulate the aggregation process (Fig. 7). Together, these studies support our hypothesis that besides primary nucleation and templated polymerization also nucleated branching plays a critical role in the Ex1Q49 aggregation process. At the date of publication, active nucleated branching along existing fibrils has only been described for glucagon [20], but to our knowledge has not been theoretically modelled in the context of HD.

Our investigations of the early steps in the aggregate formation cascade indicate that uncleaved GST-Ex1Q49 fusion protein under non-denaturating conditions is a stable oligomer, which has a very low tendency to self-assemble into large aggregates. Upon proteolytic cleavage with PP, however, GST-Ex1Q49 oligomers disintegrate and release aggregation-prone Ex1Q49 molecules, which are most probably monomers or small oligomers (see our DLS experiments, Fig. 1d). The released molecules then rapidly polymerize into large SDS-stable aggregates with a typical fibrillar morphology (Fig. 1d and e). Our observations therefore lead us to conclude that the large bundles of fibrillar Ex1Q49 aggregates are spontaneously formed *in vitro* from directly interacting monomers or very small oligomers (dimers, trimers or tetramers), which is in agreement with recently reported studies [38]. However, they do not support observations that the assembly of transient, relatively large spherical oligomers is a critical early event in the spontaneous HTTex1 aggregation cascade [34].

Our analysis of the early events in the Ex1Q49 aggregation cascade with DBAs and the epitope-specific anti-polyQ antibodies MW1 and 3B5H10 indicate that both antibodies readily detect the polyQ domain in uncleaved, soluble GST-Ex1Q49 oligomers (Fig. 2). However, MW1 antibody binding to the polyQ domain was rapidly lost upon cleavage of the GST fusion protein with PP. It has been shown previously that the MW1 antibody recognizes short linear epitopes in unfolded polyQ chains [52]. Our results suggest that linear polyQ chains are exposed to MW1 in GST-Ex1Q49 oligomers but not in released Ex1Q49 molecules. This supports previous reports that pathogenic polyQ tracts collapse and assume compact geometries in aqueous environments [62]. Previously published experiments also indicate that long polyQ chains collapse when they are analysed with FRET or TR-FRET assays [63, 64]. In contrast to MW1, released soluble Ex1Q49 molecules are readily detectable by the anti-polyQ antibody 3B5H10 (Fig. 2a), indicating that this antibody is less polyQ conformation sensitive and recognizes both linear and collapsed polyQ chains under non-denaturating conditions.

We suggest that the proteolytically released 3B5H10-reactive Ex1Q49 molecules are compact structures that have a relatively low β-sheet content [54]. This is also supported by ThT dye binding experiments (Fig. 1c), which preferentially detect higher-molecular-weight, β-sheet-rich Ex1Q49 aggregates, but not soluble Ex1Q49 monomers and small oligomers. Because of the low β-sheet content, we hypothesize that the soluble molecules have to undergo a conformational change into a stable β-sheet structure in order to form fibrillar aggregates. Such a conformational conversion is an energetically unfavourable event; it may therefore occur only very rarely in a large population of proteolytically released molecules. However, once a stable β-sheet-rich molecule has spontaneously formed, it may function as a nucleus (or seed) that can efficiently convert other non-β-sheet-rich Ex1Q49 molecules into an aggregation-competent state. Thus, we propose a spontaneous fibril formation mechanism in which a rare folding event is the rate limiting step (Fig. 2a). This hypothesis is supported by our theoretical modelling studies indicating that a nucleus size of 1 (*n_c_* = 1) best describes the polymerization of Ex1Q49 molecules into amyloid fibrils (Fig. 5d). A conformational transition from a non-β-sheet into a β-sheet structure has previously been observed in soluble polyQ monomers with biophysical methods [65], further substantiating the theoretical results. However, our experimental data do not exclude the possibility that instead of monomers small Ex1Q49 oligomers (e.g. dimers, trimers or tetramers) function as seeds and template fibril assembly. They could form from interacting Ex1Q49 monomers by primary nucleation. Probably, they have a compact β-hairpin conformation, which was previously predicted [66] and was recently identified with high resolution structural methods in fibrillar polyQ-containing HTTex1 aggregates [67]. Interestingly, the formation of HTTex1 tetramers has lately been described in cell-free assays [33]. Whether they function as nuclei (seeds) and can promote amyloid fibrillogenesis, however, remains unknown. Additional studies are therefore necessary to better elucidate the very early steps in the HTTex1 aggregation cascade and to describe the formation of small metastable oligomers. Such structures have recently been directly observed in the Ure2 aggregation process by using single molecule fluorescence microscopy [68].

Previous studies indicate that HTTex1 fibrils, once they are formed, can further aggregate into even more complex amyloid structures [34]. Our experiments with preformed Ex1Q49 seeds (Fig. 7c-d) do not support these results. Rather, we observed that small Ex1Q49 fibrils, which are rapidly formed after 5-10 h (Fig. 7c), are very stable structures, which cannot grow larger when all monomers are used up and the aggregation reactions are incubated for additional 40-45 h. Thus, the observed large bundles of Ex1Q49 fibrils (Fig. 1e and 1f) are unlikely to be formed by lateral association of small fibrils (see Fig. S19) but rather are the result of nucleated branching and templated polymerization (Fig. 5a). In this case, an initially formed small seed or nucleus grows into a large fibril bundle with multiple fibril ends by monomer addition. Together, these studies indicate that the mechanism underlying Ex1Q49 aggregation is distinct from that of other polypeptides such as the 42-residue amyloid-β (Aβ42) peptide, which forms unbranched fibrils [60]. Recent studies have shown that Aβ42 fibrils can catalyse the assembly of neurotoxic oligomers [19, 69], indicating that surface-catalysed secondary nucleation events play a key role in amyloidogenesis. In strong contrast to Ex1Q49 branches, however, these oligomers are not stably attached to the nucleating amyloid fibrils.

High resolution studies indicate that, besides large perinuclear inclusion bodies with highly complex fibrillar structures [70, 71], also smaller bundles of fibrillar HTTex1 aggregates with pathogenic polyQ tracts are detectable in the cytoplasm of mammalian cells [72, 73], supporting our hypothesis that spontaneous HTTex1 aggregation both *in vitro* and *in vivo* is dominated by this fibril formation pathway. We suggest that complex bundles of fibrillar HTT aggregates are important disease-relevant structures. They are very stable and unlikely to be degraded efficiently by cellular protein quality systems such as autophagy. This process was shown to be impaired in HD models [74, 75]. Complex bundles of mutant HTTex1 fibrils also might be responsible for the recruitment of multiple critical factors, which could lead to aberrant cellular functions. Recent studies indicate that many cellular proteins aberrantly interact with mutant HTTex1 aggregates [76]. Also, experimental evidence was obtained that proline-rich domains are exposed on the surface of HTTex1 fibrils [77, 78], suggesting that such structures might efficiently recruit SH3 and WW domain-containing signalling proteins and thereby perturb their normal cellular functions. Such proteins have previously been demonstrated to bind to mutant HTTex1 aggregates in various disease models [79, 80]. Furthermore, recent studies indicate that highly complex fibrillar HTTex1 structures interact with endomembranes in cells and lead to their local impairment [71]. Thus, the formation of complex bundles of fibrillar HTT aggregates in neurons through a nucleated branching mechanism may be of high relevance for the development and progression of HD.

Our mechanistic studies that suggest a critical role of nucleated branching for Ex1Q49 aggregation have several important implications for the development of disease-modifying therapeutic strategies for HD. For example, we found that the small molecule O4 extends the lag phase in the Ex1Q49 aggregation process (Fig. 3a-f), indicating that it slows down the formation of aggregation-competent conformations, which are a prerequisite for fibril assembly. If we assume that seeding-competent, fibrillar HTT aggregates are the toxic species in pathogenesis [28], compounds like O4 should delay fibrillogenesis and improve symptoms in HD patients. In order to be effective, however, treatment with such compounds most probably would need to start before fibrillar, seeding-competent aggregates have formed, because propagation of such structures in patient brains most likely cannot be halted. Such specific questions can now be followed up in experiments with transgenic and knock-in mouse models of HD using our set of epitope-specific antibodies and small molecules that target specific events in the Ex1Q49 aggregation cascade.

## Materials and Methods

### Protein purification and *in vitro* analysis of aggregates

The proteins GST-Ex1Q49, GST-Ex1Q48-CyPet, GST-Ex1Q48-YPet and GST were produced in *E. coli* BL21-RP cells and purified as described previously [25]. Filter retardation assays (FRAs), dot blot assays (DBAs), SDS-PAGE, dynamic light scattering, size exclusion chromatography and atomic force microscopy of aggregates were performed using standard methods. Details are provided in Supplemental Methods,

### HTTex1 protein aggregation

Aggregation of the Ex1Q49 fragment was initiated by addition of 0.28 U PreScission (PP) protease to 2 µM GST-Ex1Q49 fusion protein (50 mM Tris pH 7.4, 150 mM NaCl, 1 mM EDTA and 1 mM DTT). Reactions were incubated at 20°C (300 rpm) for 1-8 h. Samples were taken at indicated time points, frozen on dry ice and stored at −80°C. Aggregation of the Ex1Q48-CyPet/-YPet fragments was performed under comparable conditions. Details are provided in Supplemental Methods.

### Computational details for docking studies and MD simulations

O4-Ex1Q47 docking studies were performed using the program AutodockVina [55] and the previously reported structural model Ex1Q47 [53]. Quantum Mechanics/Molecular Mechanics Molecular Dynamics (QM/MM MD) simulations in explicit solvent were performed using the program CHARMM (v33b13) [81] with the CHARMM22 force field [82] for the MM region and the SCC-DFTB-D [83] method for the QM region (O4 molecule). Classical MD simulations of Ex1Q47 and Ex1Q46 [54] were performed with the NAMD program [84]. More details are provided in Supplemental Methods - Computational Details for Docking Studies and MD Simulations.

### Kinetic modelling

Models for the various aggregation mechanisms are given by sets of ordinary differential equations (ODEs). Binding and conformational transitions are described by mass-action kinetics. We used moments of the distribution to derive a finite set of equations for describing the aggregation process. These equations are solved numerically using MATLAB ode15s function. All details are provided in Supplemental Methods - Kinetic Modeling of Ex1Q49 Aggregation. In the model the rate constant *k*_1_ accounts for the binding to the template, i.e. to all aggregates equal or larger than the growth nucleus, and a conformational change of the monomers. To define a normalization procedure and compare model simulations to the FRA data we used serial dilutions of reaction mixtures and estimated the relationship between relative protein amounts and relative immunoreactivities for the antibodies applied for aggregate detection in FRAs (HD1 and CAG53b, Fig. S14a, Supplemental Data, Eqs. S30, S31). A log2 dependency was used to convert simulated protein amounts to relative immunoreactivities, which could then be compared to the measured values. We measured GST-Ex1Q49 cleavage by PP over time and found that the cleavage followed a Michaelis-Menten kinetics for which we estimated the kinetic parameters (Fig. S14b, Supplemental Methods, Eq. S5). We also performed DBAs with the CAG53b antibody at different time points after addition of PP (Fig. S14c), which showed that CAG53b detection efficiency was independent of the Ex1Q49 aggregation state. This allowed to define the relation between protein amounts and CAG53b signal (Supplemental Methods, Eqs. S32-S33).

## Acknowledgments

E.S.-G. thanks N. V. Dokholyan, University of North Carolina and R. Zhou, Columbia University, for kindly providing their calculated Ex1Q47 and Ex1Q46 structures and M. Doerr for useful discussions. E.S.-G. acknowledges a Plus-3 Grant by the Boehringer Ingelheim foundation. E.S.-G. also acknowledges the financial support of the Cluster of Excellence RESOLV (EXC 1069, infrastructure support) and the SFB1093, both funded by the DFG. E.E.W thanks K. Rau and G. Grelle for AFM studies, B. Purfürst for performing electron microscopy investigations and S. Schnögl for critical reading of the manuscript. This project was supported by grants from the HDSA, DFG (SFB740 and SFB618), BMBF (NGFN-Plus: NeuroNet), EU (EuroSpin and SynSys) and the Helmholtz Association (MSBN and HeIMA). The relevant grant numbers are SFB740: 740/2-11, SFB618: 618/3-09, NeuroNet: 01GS08169-73, EuroSpin: Health-F2-2009-241498, SynSys: HEALTH-F2-2009-242167, MSBN: NW2, HelMA: HA-215. J.W. was supported by the FORSYS program of the BMBF (grant number 0315289), and the Helmholtz Association (Personalized Medicines Initiative ‘iMED’). E.E.W. also acknowledges the support of the Berlin Institute of Health (CRG 2a3). The authors declare that they have no conflict of interest.

## Author contributions

A.S.W. designed and performed all time-resolved aggregation experiments in this study; N.U.S. and A.B. performed AFM experiments; S.P. produced and characterized recombinant proteins; E.E.W. coordinated experimental and theoretical studies and designed the overall experimental strategy; A.Z.P. performed the DLS experiments; K.K. analyzed peptide arrays; A.Z.P. designed the kinetic models and performed the mathematical simulations with the help of J.W. and K.B.; K.B-R and J.M.R-A performed the docking experiments, the MD simulations and processed the corresponding computational data; E.S-G performed the QM/MM MD simulations, processed the data, planned and directed the computational work; A.A., A.B. and L.B. performed seeding experiments; C.H. analyzed data and generated figures; A.S.W., E.E.W., A.Z.P., J.W., and E.S-G. wrote the paper.

## Supplemental Data

Supplemental Methods

Supplemental References

Legends of Supplemental Figures S1 to S19

Legends of Supplemental Movies S1 and S2

Supplemental Tables S1 to S5 and Legends

Supplemental Figures S1 to S19

Supplemental Movies S1 and S2 (separate files)

