## Supplementary Materials for "Self-assembly of mutant huntingtin exon-1 fragments into large complex fibrillar structures involves nucleated branching"

### Supplemental Methods

#### Cloning of expression vectors

For the construction of plasmids encoding CyPet- and YPet-tagged HTTEx1Q48 fusion proteins, the coding sequence of HTTEx1Q48 was PCR-amplified from pGEX-6P1-HTTEx1Q48 using the primers 5'-gacgacgaattcatggcgaccctg-3' and 5'-gacgacctcgagtggtcggtgcagcgg-3'. The resulting PCR product was digested with the restriction enzymes EcoRI and NotI. Additionally, CyPet cDNA was PCR amplified from pBAD33-CyPet-His (Addgene plasmid #14030) [1] with the primers 5'-acgacctcgagggtggcggtggcggtatgtctaaaggtgaagaattattcgg-3' and 5'-gacgacgcggccgcttatttgtacaattcatccataccatg-3'. YPet cDNA was amplified from pBAD33-YPet-His (Addgene plasmid #14031) [1] with the primers 5'-gacgacctcgagggtggcggtggcggtatgtctaaaggtgaagaattattcactgg-3' and 5'-gacgacgcggccgcttatttgtacaattcattcataccctcg-3'. The resulting PCR fragments were cloned into the plasmids pGEX-6P1 using the EcoRI/XhoI/NotI restriction sites to obtain the plasmids pGEX-6P1-HTTEx1Q48-CyPet and -YPet, respectively.

#### Proteins, antibodies and chemical compounds

The proteins GST-Ex1Q49 and GST-Ex1Q48-CyPet and -YPet were produced in *E. coli* BL21-CodonPlus-RP and purified under native conditions by affinity chromatography on glutathione agarose beads as described [2]. Purified proteins were dialyzed over night at 4 °C against 50 mM Tris-HCl pH 7.4, 150 mM NaCl, 1 mM EDTA and 5% glycerol, shock-frozen in liquid nitrogen and stored at -80°C. Protein concentrations were determined with a NanoDrop spectrophotometer. Before aggregation experiments were performed, proteins were ultracentrifuged (Beckman Coulter, Optima TL, rotor TLA100.3) at 208,000 g for 20 min at 4°C. The polyclonal anti-HTT antibodies CAG53b and HD1 have been described previously [3]. The monoclonal antibodies MW1 and MW8 [4] were obtained from the Developmental Studies Hybridoma Bank (University of Iowa). The antibody 3B5H10 [5] was purchased from Sigma (Germany). The anti-GFP antibody (ab290) was purchased from Abcam. The compound O4 (ZIZ074584) was purchased from Zelinsky Institute Inc., Newark (USA). Stock solutions of the compound (20 mM) were prepared in DMSO and stored at -20°C.

**Filter retardation assays (FRAs)**

FRAs were performed as described previously [6]. Briefly, equal volumes of 500 ng of Ex1Q49 aggregation reactions and 4% sodium dodecyl sulfate (SDS) solution containing 100 mM dithiothreitol (DTT) were mixed and boiled at 96°C for 5 min. Samples were filtered through a cellulose acetate membrane with 0.2 µm pores (OE66, Schleicher and Schuell, Germany) and washed twice with 100 µl 0.2% SDS. Membranes were blocked in Tris-buffered saline containing 5% skim milk and 0.05% Tween 20. Aggregates retained on the filter membrane were detected using the CAG53b antibody (1:2000), HD1 antibody (1: 10,000) or anti-GFP antibody (1:5000) and secondary antibodies conjugated to alkaline phosphatase or peroxidase (Promega, Germany). Signals were quantified using the AIDA image analysis software (Raytest, Straubenhardt, Germany).

**SDS-PAGE and immunoblotting**

Samples of aggregation reactions were mixed with loading buffer [50 mM Tris HCl, pH 6.8, 2% SDS, 10% (v/v) glycerol and 0.1% bromophenol blue] and boiled at 96°C for 5 min. Samples were loaded onto 10% SDS gels. SDS-PAGE and Western blotting were performed according to a standard protocol. Membranes were blocked with 5% skim milk in Tris-buffered saline (TBS) containing 0.05% Tween 20 and incubated with the HD1 (1:2000) antibody and secondary antibodies conjugated to alkaline phosphatase (Promega, Germany). Signals were quantified using the AIDA image analysis software (Raytest, Straubenhardt, Germany).

**Dot blot assays (DBAs)**

To detect proteins under non-denaturing conditions DBAs were performed as described previously [7]. Briefly, 250 ng of Ex1Q49 aggregation reactions were spotted onto nitrocellulose membranes. Membranes were blocked for 30 min with 5% skim milk in TBS containing 0.05% Tween 20. Membranes were then incubated with the monoclonal antibodies MW1, 3B5H10 or MW8 dissolved in 5% skim milk in TBS containing 0.05% Tween 20 (MW1 1:2000; MW8 1:2000 and 3B5H10 1:5000) and developed using alkaline phosphatase conjugated secondary antibodies (Promega, Germany). Signals were quantified using the AIDA image analysis software (Raytest, Straubenhardt, Germany).

**Dynamic light scattering (DLS) measurements**

The size of aggregates was measured at 20°C without agitation by dynamic light scattering (Malvern Nano Zetasizer ZS, with a green laser $\lambda$=532 nm, and backscattered detection at an angle $\theta$=173 deg). The GST-Ex1Q49 fusion protein was centrifuged (208,000 g and 4°C) for 20 min prior to investigation by DLS (Beckman Coulter, Optima TL, rotor TLA100.3). To remove potential dust, samples were additionally centrifuged at 4°C using a table-top centrifuge (18,000 rpm) prior to the transfer to a quartz cuvette for light scattering measurements. Due to the high measurement noise with DLS at 2 µM GST-Ex1Q49 all measurements were performed using 20.9 µM GST-Ex1Q49 and correspondingly higher PP concentration. O4 was diluted to a final concentration of 20.9 µM.

Dynamic light scattering yields the normalized time autocorrelation function of the intensity of the scattered light $g^{(2)}(\tau)$

$$g^{(2)}(\tau) = \frac{\langle I(t)I(t+\tau)\rangle}{\langle I(t)\rangle^2} \tag{S1}$$

Each correlation spectrum is the average of 10 traces à 3 s each. The size distribution by intensity has been derived by deconvolution of the autocorrelation function computed using the CONTIN algorithm as implemented in the Malvern software.

The z-averaged diffusion coefficient $D_z$ of the aggregates in solution was computed as described in [8]. Briefly, the autocorrelation function is expanded in terms of the moments around the mean of the decay rate distribution. An expansion up to order two in $\tau$ gives the approximation

$$g^{(2)}(\tau) \approx B + \beta \exp(-2\,\mu_1)\left(1 + \frac{\mu_2\tau^2}{2!}\right)^2 \tag{S2}$$

Where $\mu_i$ is $i$-th moment around the mean, $\beta$ a factor that depends on the experimental geometry and $B$ the baseline, which can differ from its theoretical value of 1 due to noise. The first moment relates to the z-averaged diffusion constant

$$D_z = \mu_1/q^2\,, with\ q = \frac{4\pi n}{\lambda}\sin\left(\frac{\theta}{2}\right). \tag{S3}$$

For our setup and a refractive index $n = 1.333$ the scattering wave vector was $q = 31.4281\ \mu m^{-1}$. Equation S2 was fitted to the autocorrelation data, using the non-linear fitting routine lsqnonlin from Matlab to obtain $\mu_1$, $\mu_2$, $B$ and $\beta$. The z-averaged diameter $2R_z$ is calculated from the Einstein-Stokes equation

$$2R_z = \frac{kT}{3\pi\,\eta\,D_z}. \qquad \text{(S4)}$$

Where $k$ is the Boltzmann-constant, $T$ the temperature, and the viscosity $\eta$ = 1.03424 cP.

### Atomic Force Microscopy (AFM)

Aliquots of 500 ng of Ex1Q49 aggregation reactions, pre-cleaved for 3 h at 5°C (300 rpm) before initiation of aggregation at 20°C, were spotted onto freshly cleaved mica and allowed to adhere for 10 min. Then, they were washed 4 times with 40 µl distilled water. The samples were dried to completion at room temperature and imaged in air with a digital multimode NanoscopeIII scanning probe microscope operating in tapping mode.

### Size-exclusion chromatography

Gel-filtration experiments were performed with the Äkta purifier system (Amersham Pharmacia) using a Superdex 30/100 GL column. Proteins were eluted in TBS buffer (50 mM Tris-HCl, pH 7.4, 150 mM NaCl, 0.01 mM EDTA). All experiments were carried out at 4°C and a flow rate of 300 µl $min^{-1}$. The column was calibrated under the same conditions with ribonuclease A (13.7 kDa), ovalbumin (43 kDa), conalbumin (75 kDa), aldolase (158 kDa), ferritin (440 kDa) and thyroglobulin (669 kDa). The exclusion volume was determined with blue Dextran (2,000 kDa).

**Protein precipitation**

Proteins were concentrated by TCA precipitation. TCA was added to a final concentration of 13% to the protein solution. Proteins were incubated at -20°C for 5 min and 15 min at 4°C and then centrifuged for 15 min at 13,500 rpm in a Hettich Micro 22R centrifuge at 4°C. Pellets were dried and resuspended in loading buffer (50 mM Tris HCl, pH 6.8, 2% SDS, 10% (v/v) glycerol, and 0.1% bromophenol blue).

**FRET-based HTT aggregation assay**

The aggregation of GST-Ex1Q48-CyPet and GST-Ex1Q48-YPet was performed at indicated concentrations (using an equimolar ratio of both sensor proteins) in 50 mM Tris-HCl pH 7.4, 150 mM NaCl, 1 mM EDTA and 1 mM DTT. Spontaneous aggregation was initiated by addition of 14 U PreScission protease (GE Healthcare) per nmol purified GST-Ex1Q48-CyPet/YPet fusion protein. The solution was mixed with preformed sonicated Ex1Q49 aggregates (seeds) at varying concentrations and transferred to a black 384-well plate. Fluorescence signals were measured every 20 min following a 5 s pulse of vertical shaking with a Tecan M200 fluorescence plate reader at 25 °C for up to 50 h. CyPet donor fluorescence was measured at excitation (Ex): 435 nm/emission (Em): 475 nm; YPet acceptor fluorescence at Ex: 500 nm/Em: 530 nm; the FRET channel (*DA*) was recorded at Ex: 435 nm/Em: 530 nm. Fluorescence signals in all channels were background corrected by subtracting the background fluorescence of unlabelled Ex1Q48. To calculate the sensitized emission, signals in the FRET channel were corrected for donor bleed-through (*cD*) and acceptor cross excitation (*cA*) using donor- and acceptor-only samples. Finally, sensitized emission was normalized to the acceptor signals. In brief, the FRET efficiency $E$ (in %) was calculated as follows: $E = (DA - cD * DD - cA * AA) / AA$ with $DD$ = donor channel signal and $AA$ = acceptor channel signal.

### Computational details for docking studies and molecular dynamics simulations

The crystal structure of the non-pathogenic Ex1Q17 fragment was reported by Kim et al [9]. However, less information is available concerning the structural properties of HTT peptides with pathogenic polyQ tracts. Here, the Ex1Q47 structure of the HTT peptide with flanking regions and a glutamine repeat length of 47 computed by Dokholyan [10] was used as a starting point for our calculations. In addition, a different conformation of Ex1Q46, corresponding to the most populated structure with β-sheet content (third cluster), was also employed [11]. Like the very similar Ex1Q49 fragment used in our experiments, Ex1Q47 and Ex1Q46 provide a reliable model to investigate the effect of O4 in HTTex1 peptides with a pathogenic polyQ expansion.

For the docking calculations of O4 on Ex1Q47 two grids were used: the flexible grid **A** with a size of 48x46x48 Å that includes Ex1Q47 in its full extension and the grid **D** with a size 38x28x30 Å that comprises the N-terminal region (residues Gln25 to Gln37, Gln43 to Gln53, Pro70 to Gln76 and Leu88 to Pro100) of Ex1Q47. Additional tests with other grid sizes or rigid docking corroborated the results from **A** and **D.** Based on the docking affinities and the geometrical similarity between the resulting Ex1Q47-O4 structures, the best scored cases from both grids were considered for further simulations. Since the docking affinities only provide a preliminary overview of the binding sites, we also performed Quantum Mechanics/Molecular Mechanics Molecular Dynamics simulations (QM/MM MD, 400 ps, time step of 1 fs). The QM region (O4 molecule) was treated using the SCC-DFTB method including dispersion corrections (SCC-DFTB-D) [12, 13], the CHARMM22 force field [14] to describe the protein and the TIP3P model [15] for water. The QM/MM MD simulations allowed the evaluation at the QM level of the relative stability of O4 in different Ex1Q47 regions and were performed following a work methodology described by us elsewhere [16]. The VMD program [17] was used for visualizing all the MD trajectories.

The four most stable complexes of O4 and Ex1Q47 as indicated by the QM energies during the QM/MM MD were selected for further classical MD simulations. In addition, the complex of O4 with Ex1Q46 was simulated using two independent replicas. In all cases the reference system without ligand was also simulated. The MD simulations also performed using the NAMD program [18] and were extended to 200 ns in each case for a total time of 1.6 microseconds (considering both Ex1Q47 and Ex1Q46). The O4 parameters were generated by the Swissparam server [19] and their quality verified by comparison with QM and QM/MM MD calculations. Inter-residues contact maps were created using Gromacs (v4.5.6) with a 1 nm cut-off [20].

### Kinetic modeling of Ex1Q49 aggregation

Each model describes an aggregation mechanism and is given by a set of ordinary differential equations (ODEs).

#### PreScission protease-mediated protein cleavage

Since we do not observe Ex1Q49 aggregation in the absence of the PreScission protease (PP) (**Fig. 1b**), we here assume that only Ex1Q49 molecules without GST can aggregate (**Fig. 5a**). We described the cleavage of GST from GST-Ex1Q49 by PP using different kinetics and found the best description by using a Michaelis-Menten kinetics:

$$\dot{X}_g = -v_P = -v_{max}\frac{X_g}{X_g+K_P}\ . \qquad \text{(S5)}$$

Here, $X_g$ is the concentration of GST-Ex1Q49, $K_P$ the Michaelis-Menten constant, and $v_{max}$ the maximal velocity. In the experiments, the amount of PP is increased proportionally to the initial concentration of GST-Ex1Q49, thus $v_{max} = k_r X_g(0)$. The parameters are fitted to PP digestion data for two initial concentrations of GST-Ex1Q49 (2 µM and 20 µM) as detected with HD1 (**Fig. S14b**) using the relative immunoreactivity to protein conversion for HD1 as described in Fitting of HTTex1 kinetic models to data. We obtained $K_P$ = 4.21 µM and $k_r$ = 1.99/h.

#### Primary nucleation and templated polymerization

Fibril formation often involves a nucleation-dependent polymerization reaction where the formation of growth nuclei from soluble proteins is slow [21-23]. We denote this initial step primary nucleation. The binding of monomers to a nucleus (also called template) [24] in the sense of elongation growth is denoted templated polymerization. This leads to a conformational change of the newly bound protein unit and the formation of a stable complex, which can further serve as template. In our case, the templated polymerization reaction can be considered as quasi-irreversible as we observed SDS-stable complexes. Mathematical models for such mechanisms have been previously described (see e.g. [25-27]). Here we recapitulate the assumptions and present model equations in the context of the additional GST cleavage.

We denote by $x_1$ the concentration of Ex1Q49 monomers prior to formation of the nucleus. We denote this conformation C1. The concentrations of all complexes in this conformation are captured in lower-case $x_i$, where $i$ denotes the number of Ex1Q49 proteins in a complex. The parameter $n_c$ gives the size of the first stable complex, the nucleus, that acts as template. All complexes in this conformation are captured in upper-case $X_i$, where $i$ denotes the number of Ex1Q49 proteins in the complex. We denote this conformation C2.

Using mass action kinetics for protein binding and conformational changes the infinite set of ODEs reads

$$\dot{x}_1 = v_P - k_{n1}\, n_c\, x_1{}^{n_c} + k_{-n1} n_c\, X_{n_c} - k_1 x_1 \sum_{i=1}^{\infty} X_i \tag{S6}$$

$$\dot{X}_i = k_{n1} \delta_{i,n_c} x_1{}^{n_c} - k_{-n1} \delta_{i,n_c} X_{n_c} + k_1 x_1 (X_{i-1} - X_i), \quad i \geq n_c. \tag{S7}$$

By definition, $X_i = 0$ in all cases $i < n_c$. The Kronecker symbol is defined by $\delta_{i,j} = 1$ for $i = j$ and 0 otherwise. Monomers $x_1$ are produced by the cleavage of GST with rate $v_P$ (Eq. (S5) which also belongs to the equation system of the model). Primary nucleation is characterized by the forward rate constant $k_{n1}$. The term $k_{n1}\, x_1{}^{n_c}$ (Eq. (S7)) gives the rate of nucleus formation, and $-k_{n1}\, n_c\, x_1{}^{n_c}$ (Eq. (S6)) describes the consumption of $x_1$ molecules by this process. The scaling factor $n_c$ in the latter accounts for the stoichiometry of the process in which $n_c$ monomers form one nucleus. For $n_c > 1$, the higher order of the nucleation reaction can be accounted for by reversible binding of monomers to form small complexes and a fast equilibrium of $x_1$ with $x_{n_c-1}$. In this case, we have $x_{n_c-1} = x_1^{(n_c-1)}/K$, where $K$ is the dissociation constant of the dissociation of $x_{n_c-1}$ into $x_1$. Further binding of a monomer to $x_{n_c-1}$ with rate constant $k_1$ causes a conformational change to C2 leading to a stable complex $X_{n_c}$. Therefore, $k_{n1} = k_1/K$ denotes an effective rate constant for the formation of the first stable complex $X_{n_c}$. Assuming $x_{n_c-1} \ll x_1$ (i.e. $K \gg 1$) we only need to consider an equation for $x_1$. For $n_c$ = 1, there is no fast equilibrium and the nucleation process describes a slow conformational change from $x_1$ to $X_1$ which we describe by a reversible nucleation process with the backward, de-nucleation reaction characterized by the rate constant $k_{-n1}$. This avoids that the characteristic time for the conformational change becomes extremely long (1/$k_{n1}$ > 30 h for the model without reversible nucleation, parameters not shown) compared to less than 12 min in case of a reversible nucleation (1/($k_{n1}$ + $k_{-n1}$) < 12 min, **Table S5**). For a fair comparison between models, we also considered a de-nucleation reaction for $n_c$ > 1, but found that a value of $k_{-n1} = 0$ does not change the conclusions of the models concerning the role of branching and $n_c$ = 1.

In the literature the complex $x_{n_c-1}$ is also often referred to as the nucleus. However, this leads to confusion for $n_c = 1$ and the distinction to spontaneous aggregation. We thus denote by the nucleus the first stable complex $X_{n_c}$ of size $n_c$.

During templated polymerization a complex elongates by binding of Ex1Q49 monomers in conformation C1, thereby, the newly attached protein unit switches to the stable conformation C2. The process is assumed to be irreversible and occurs with rate constant $k_1$. The terms $k_1 x_1 X_{i-1}$ and $-k_1 x_1 X_i$ in Eq. (S7) account, respectively, for the appearance and disappearance of a complex of size $i$ by binding of a monomer. The term describing the consumption of $x_1$ by templated polymerization is given by the sum of the elongation reaction rates $k_1 x_1 X_i$ over every possible complex: $k_1 x_1 \sum_{i=1}^{\infty} X_i = k_1 x_1 (m_0 - x_1)$ , where, $m_0$ is the 0th moment of the protein distribution.

We introduce the moments of the protein size distribution as

$$m_k = x_1 + \sum_{i=1}^{\infty} i^k X_i. \quad \text{(S8)}$$

The ODEs for the first 3 moments read

$$\dot{m}_0 = v_P + (1 - n_c) k_{n1} {x_1}^{n_c} - (1 - n_c)\, k_{-n1} X_{n_c} - k_1 x_1 (m_0 - x_1) \quad \text{(S9)}$$

$$\dot{m}_1 = v_P \quad \text{(S10)}$$

$$\dot{m}_2 = v_p + n_c(n_c - 1) k_{n1} {x_1}^{n_c} - n_c(n_c - 1)\, k_{-n1} X_{n_c} + 2\, k_1 x_1 (m_1 - x_1) \quad \text{(S11)}$$

The 0th moment $m_0$ gives the sum of the concentrations of all complexes, the 1st moment $m_1$ gives the total concentration of Ex1Q49 proteins, the 2nd moment $m_2$ is proportional to the weighted average size of the Ex1Q49 complexes (for t > 0, without GST-Ex1Q49) given by $m_2/m_1$ [28]. The moments are used to derive a finite set of equations for comparison to the experimental data (see Fitting of HTTex1 kinetic models to data).

For $n_c = 1$, it is possible to allow for a non-templated polymerization step where a monomer in conformation C2 can also elongate a fibril. The introduction of such additional step did not significantly improve the quality of the fits and is therefore not considered here.

For spontaneous polymerization, that is $n_c = 0$, and $x_1 = 0$, the model equations (S6), (S7), (S9), (S10) (S11) simplify to

$$\dot{X}_1 = v_P - k_1 X_1 (m_0 + X_1) \quad \text{(S12)}$$

$$\dot{X}_i = k_1 X_1 (X_{i-1} - X_i), i \geq 2 \quad \text{(S13)}$$

$$\dot{m}_0 = v_P - k_1 X_1 m_0 \quad \text{(S14)}$$

$$\dot{m}_1 = v_P \quad \text{(S15)}$$

$$\dot{m}_2 = v_P + 2k_1 X_1 m_1. \quad \text{(S16)}$$

**Primary nucleation, templated polymerization and nucleated branching**

In a model that includes templated polymerization and nucleated branching the number of proteins in a complex can increase by attaching Ex1Q49 monomers in conformation C1 to it either by elongation of an existing fibril (**Fig. 5a**, red arrows, templated polymerization) or by branching from the surface of an existing fibril by secondary nucleation (**Fig. 5a**, blue arrows, nucleated branching). As for templated polymerization, nucleated branching can only occur at proteins in conformation C2. For simplicity, we assume that each protein unit has one polymerization and one branching site, furthermore we neglect sterical impairments between branches. The nucleated branching reaction allows nucleating a new growing filament and we assume a reaction order $n_b$ for the monomers, i.e. $n_b$ Ex1Q49 monomers are attached, along their polymerization sites, to the branching site of a protein for forming a branch. We refer to an unoccupied polymerization site within a complex in conformation C2 as growing end, and each growing end defines one branch of the complex. Therefore, templated polymerization is proportional to the number of branches of a complex and nucleated branching is proportional to the number of unoccupied branching sites in a complex. Consequently, the reactions of complexes with similar numbers of branches and polymerization sites can be described by the same rate of templated polymerization and of nucleated branching, despite the possibly different spatial structures of the complexes. We hence categorize the complexes according to their number of branches and number of protein units.

Denote by $X_{i,j}$ the concentration of complexes in conformation C2 with $i$ protein units and $j$ the number of growing ends or branches. Thereby, $i \geq n_c$ due to primary nucleation with nucleus size $n_c$ and $j \geq 1$. A nucleus of size $n_c$ has one growing end, and thus the number of branches in a complex can be at most equal to the number of monomers in the complex ($i \geq j$). Therefore, $X_{i,j} = 0$ for $j > i$ (and for $j < 1$). Equality of $i$ and $j$ occurs only in a monomer, i.e. in $X_{1,1}$, or if a complex only contains branches of length 1 (which both requires $n_c = 1$).

For a model with nucleated branching, we can define the moments of the system composed of $x_1$ and all $X_{i,j}$ as

$$m_{a,b} = x_1 + \sum_{i=n_c}^{\infty} \sum_{j=1}^{i} i^a j^b X_{i,j} \qquad \text{(S17)}$$

$m_{0,0}$ is the sum of the concentration of each complex, and consequently it contains the information about the number of aggregates of any size in the system. In $m_{0,1}$ the number of branches of each aggregate is taken into account and thus it is the overall concentration of branches in the system. $m_{1,0}$ counts the number of Ex1Q49 proteins of which each aggregate is composed and thus contains the total concentration of Ex1Q49 proteins in the system. The ODE system for the protein aggregation model with primary nucleation, templated polymerization and nucleated branching can be expressed as function of the moments as well as the monomer and complex concentrations

$$\dot{x}_1 = v_p - k_{n1} n_c x_1^{n_c} + k_{-n1} n_c X_{n_c,1} - k_1 x_1 (m_{0,1} - x_1) \quad \text{(S18)}$$

$$- n_b k_b x_1^{n_b} (m_{1,0} - m_{0,1} + m_{0,0} - x_1)$$

$$\dot{X}_{i,j} = \delta_{n_c,i}\, \delta_{1,j}\, (k_{n1} x_1{}^{n_c} - k_{-n1} X_{n_c,1}) + k_1\, j x_1 X_{i-1,j} \quad \text{(S19)}$$

$$+ k_b[(i - n_b) - (j - 1) + 1] x_1^{n_b} X_{i-n_b,j-1} - k_1\, j x_1 X_{i,j}$$

$$- k_b[i - j + 1] x_1^{n_b}\, X_{i,j}$$

where $i \geq n_c$, $j \leq i$. We describe each term in detail in the following. For equation (S18) recall that $v_p$ is the rate with which Ex1Q49 monomers flow into the system (Eq. (S5) which also belongs to the equation system of the model). Primary nucleation and templated polymerization are accounted for as in the model without nucleated branching (see primary nucleation and templated polymerization for further details). We assume that nucleated branching does not occur for complexes in conformation C1 (i.e. $x_1$). The term $k_1 x_1 (m_{0,1} - x_1)$ denotes the consumption of $x_1$ via templated polymerization. Here monomers in conformation C1 bind to growing ends of complexes in conformation C2. The concentration of growing ends in C2 is given by $m_{0,1} - x_1$ (see Eq. (S17) and explanation). For $X_{i,j}$ (S19), templated polymerization increases its concentration by the term $k_1\, j x_1 X_{i-1,j}$. This accounts for the elongation growth reaction of an aggregate containing one monomer less (but the same number of branches *j*), $X_{i-1,j}$, on any of its branches and thus being proportional to the number of branches $j$. Additionally, $X_{i,j}$ can be elongated along any of its branches and thus its concentration is reduced by the term $k_1\, j x_1 X_{i,j}$.

The nucleated branching reactions are characterized by the rate constant $k_b$. The term $n_b k_b x_1^{n_b} (m_{1,0} - m_{0,1} + m_{0,0} - x_1)$ in (S18) accounts for the loss of monomers due to branching. A branch can be formed by nucleation of $n_b$ monomers at once similar to primary nucleation. We assumed that nucleated branching can occur at any EX1Q49 protein in conformation C2 that does not have yet a branch and the expression $m_{1,0} - m_{0,1} + m_{0,0} - x_1$ gives the concentration of unoccupied branching sites of complexes in C2. The total number of branching sites, occupied and not occupied, equals the total concentration of Ex1Q49 proteins ($m_{1,0}$). From this, the concentration of branches, $m_{0,1}$, is subtracted since each branch occupies one branching site – except for the first branch of every aggregate, which is there from the beginning. We have to correct the number of free binding sites by exactly one binding site per aggregate and consequently we have to add the total number of aggregates (minus the amount of proteins in conformation C1), $m_{0,0} - x_1$, to obtain the correct number of free branching sites. For $X_{i,j}$, the term $k_b[(i - n_b) - (j - 1) + 1]x_1^{n_b} X_{i-n_b,j-1}$ in (S19) represents nucleated branching at an aggregate with one branch less and also $n_b$ monomers less, $X_{i-n_b,j-1}$. It is proportional to the number of free branching sites in the aggregate $X_{i-n_b,j-1}$, $(i - n_b) - (j - 1) + 1$, which is the total number of monomers in the aggregate, $(i - n_b)$, subtracted by the number of branches in the aggregate $(j - 1)$ (since every branch occupies one binding site), and finally correcting for the one branching site of the initial nucleus, which is a branch although not having an occupied branching site. The term $k_b[i - j + 1]x_1^{n_b} X_{i,j}$ in (S19) constitutes nucleated branching on any of the ($i$ - $j$ + 1) free branching sites of $X_{i,j}$. Note that by setting $k_b = 0$, the model described by (S18) and (S19) is the same as that described by (S6) and (S7) with $X_{i,1} = X_i$.

The derivatives of the moments of the system can be calculated from the ODE system given in (S18) and (S19). We obtain for the derivatives of the total concentration of aggregates of any size, the concentration of branches and the total concentration number of Ex1Q49 monomers in the system:

$$\dot{m}_{0,0} = v_p + (1 - n_c)k_{n1}{x_1}^{n_c} - (1 - n_c)k_{-n1}X_{n_c,1} - k_1 x_1(m_{0,1} - x_1) - n_b k_b x_1^{n_b}(m_{1,0} - m_{0,1} + m_{0,0} - x_1) \quad \text{(S20)}$$

$$\dot{m}_{0,1} = v_p + (1-n_c)k_{n1}{x_1}^{n_c} - (1-n_c)k_{-n1}X_{n_c,1} - k_1x_1(m_{0,1}-x_1) + (1 - n_b)k_bx_1^{n_b}(m_{1,0} - m_{0,1} + m_{0,0} - x_1) \qquad \text{(S21)}$$

$$\dot{m}_{1,0} = v_p. \qquad \text{(S22)}$$

$$\dot{m}_{1,1} = v_p + k_bx_1^{n_b}(m_{1,0} + (n_b-1)m_{1,1} - n_bm_{0,2} + m_{2,0} + n_bm_{0,1} - (1+n_b)x_1) \qquad \text{(S23)}$$

$$\dot{m}_{0,2} = v_p + k_bx_1^{n_b}\left((1-n_b)(m_{0,0}+m_{1,0}) + (1+n_b)m_{0,1} + 2m_{1,1} - 2m_{0,2} + (n_b-3)x_1\right) - k_1x_1(m_{0,1}-x_1) + (1-n_c)k_{n1}\,x_1^{n_c} + (n_c-1)k_{-n1}X_{nc,1} \qquad \text{(S24)}$$

$$\dot{m}_{2,0} = v_p + k_bx_1^{n_b}\left((n_b^2+n_b)(m_{1,0}-x_1) + (n_b^2-n_b)(m_{0,0}-m_{0,1}) + 2\,n_b(m_{2,0}-m_{1,1})\right) + 2\,k_1x_1(m_{1,1}-x_1) + k_{n1}(n_c^2-n_c)x_1^{n_c} - k_{-n1}(n_c^2-n_c)X_{nc,1} \qquad \text{(S25)}$$

The second moments characterize the distribution of protein units and branches in the complexes. The weighted average number of branches (*t* > 0), *B*, and weighted average number of protein units *M* per aggregate, i.e. an average aggregate size, is given (for t > 0, without GST-Ex1Q49) are given by

$$B = \frac{m_{0,2}}{m_{0,1}}, M = \frac{m_{2,0}}{m_{1,0}}. \qquad \text{(S26)}$$

respectively. The values shown in **Fig. 7b**, stem from simulations up to *t* = 50h after addition of PP. A longer simulation gave a quasi-identical curve indicating that the system has reached a steady state.

Seeding was modeled by altering the initial conditions of the system. We simulated the initial GST-Ex1Q49 concentration of 2 μM and assumed seeds of size $M_s$ = 2.

Thus, we set $X_{2,1}(0) = X_{2,2}(0) = fX_g(0)/(M_s * 2)$ for the fraction of seeds *f*=[Seed]/[GST-Ex1Q49], where [GST-Ex1Q49] is the initial concentration of GST-Ex1Q49, $X_g(0)$.

### Kinetic modeling of the inhibitory drug O4

To identify the mode of action of O4 we model three different inhibition scenarios (**Fig. 5b**). Inhibition of the primary nucleation reaction (green symbol, **Fig. 5d**) according to

$$k_{n1}([O4]) = k_{n1}\frac{{K_{I1}}^{n_1}}{{K_{I1}}^{n_1} + [O4]^{n_1}}, k_{-n1} = \text{ const.}, \quad k_b = \text{const.}, k_1 = \text{ const.} \tag{S27}$$

where only the primary nucleation rate is affected by O4, whereas the other processes remain unaffected. The other two scenarios are inhibition of nucleated branching (blue symbol, **Fig. 5d**)

$$k_b([O4]) = k_b\frac{{K_{I2}}^{n_2}}{{K_{I2}}^{n_2} + [O4]^{n_2}}, k_{-n1} = \text{ const.}, \quad k_{n1} = \text{const.}, k_1 = \text{ const.} \tag{S28}$$

and inhibition of templated polymerization (red symbol, **Fig. 5d**)

$$k_1([O4]) = k_1\frac{{K_{I3}}^{n_3}}{{K_{I3}}^{n_3} + [O4]^{n_3}}, k_{-n1} = \text{ const.}, \quad k_{n1} = \text{const.}, k_b = \text{ const.} \tag{S29}$$

The concentration dependency on O4 was chosen as general and simple as possible and of the form as used for non-competitive enzyme inhibition.

For $n_c$ = 1, we also tested the case where O4 targets the back reaction of the reversible primary nucleation process. In this case we assume that O4 promotes an increase in the rate constant $k_{-n1}([O4]) = k_{-n1} + \bar{k}_{-n1}\frac{[O4]^{n_m}}{{K_m}^{n_m}+[O4]^{n_m}}$ causing accumulation of aggregation incompetent monomers. We found that this inhibition mode could reproduce the data. However, the fit is best for the case where $k_{n1}$ is influenced by O4 (data not shown). We therefore only show the results for the first case (inhibition of the primary nucleation).

### Fitting of HTTex1 kinetic models to data

#### Conversion of protein amount into relative immunoreactivity

For comparison of models and data, the simulated protein concentrations need to be compared with immunoreactivities. We convert protein concentrations to relative immunoreactivities (*RI*) using experimentally derived calibration curves for the employed antibodies (see below and **Fig. S14a**). Calibration curves are constructed by measuring the immunoreactivities from different dilutions of the reaction mixture (see Ex1Q49 protein aggregation in Material and Methods). For the antibody HD1 (used to detect the cleavage of GST-Ex1Q49 by PP) we used the reaction mixture at time point 0h just after addition of PP. For CAG53b we used the reaction mixture after 8h. For both antibodies we found that the relative immunoreactivity *RI* depends non-linearly on the relative protein amount *P* (see **Fig. S14a**). Here, the relative protein amount *P* is given as percentage of the protein amount recognized by the antibody in the respective immuno-detection assay for the undiluted aggregation mixture. RIs are obtained by normalizing by the signal intensity measured for the undiluted aggregation mixture.

To quantify the non-linear relationships we tested linear, hyperbolic and log dependencies.

For CAG53b, a log2 dependency described the data best

$$RI = r_1 \log_2(P + r_2) + r_3 \qquad \text{(S30)}$$

with the coefficients $r_1$=18.01, $r_2$=2.24, $r_3$=-20.25. For HD1 we used a combination of a linear function at low protein amounts and a log2 dependency for higher amounts

$$RI = \begin{cases} r_3 P, \text{ for } P \leq r_4 \\ r_1 \log_2(P + r_2) + r_5, \text{ for } P > r_4 \end{cases} \qquad \text{(S31)}$$

The coefficients are $r_1$=32.07, $r_2$=6.12, $r_3$=0.09, $r_4$=6.26. The last parameter is set in a way that *RI* is a continuous function of *P*, that is $r_5 = r_3 r_4 - r_1 \log_2(r_4 + r_2)$=-115.85. In addition, we performed model fits to the experimental data assuming a linear relationship between relative protein amount and relative immunoreactivity. We found that the conclusions about the importance of nucleated branching and nucleus of size 1 remained unchanged.

**Fitting of the models to SDS stable aggregates detected by CAG53b and estimation of kinetic parameters**

Experimentally, we found that dot blots of the reaction mixture yielded the same CAG53b signal for up to 24 hours (**Fig. S14c**). This suggests that each Ex1Q49 protein, independently of its aggregate state, contributes equally to the signal. The filter retardation assay (FRA) retains only SDS stable aggregates larger than a certain size. We model this by assuming that the FRA detects complexes larger than or equal to a size $C_{min}$, whose value we fit together with the other kinetic parameters of the model to the data. Thus, the measured CAG53b signal relates to the sum of all Ex1Q49 proteins bound in complexes of size larger or equal $C_{\min}$ in the model.

For the fit of the kinetic models with and without nucleated branching assuming detection of very small aggregates in the FRA assay (**Fig. S16b and c**), we fixed $C_{\min}=n_c$, meaning that all molecules of conformation C2 could be readily detected and thus contribute to the signal.

We denote by *CAGP* the simulated relative amount of protein, normalized to the total amount of protein $X_g(0)$, detected in the filter retardation assay by CAG53b

$$CAGP(t) = \frac{c_1}{X_g(0)} \sum_{i=C_{min}}^{\infty} iX_i \qquad \text{(S32)}$$

$$= \frac{c_1}{X_g(0)} \left( m_1 - x_1 - \sum_{i=n_c}^{C_{min}-1} iX_i \right)$$

For the model with nucleated branching this reads

$$CAGP(t) = \frac{c_1}{X_g(0)} \left( \sum_{i=C_{min}}^{\infty} \sum_{j=1}^{i} iX_{i,j} \right) \qquad \text{(S33)}$$

$$= \frac{c_1}{X_g(0)} \left( m_{1,0} - x_1 - \sum_{i=n_c}^{C_{\min}-1} \sum_{j=1}^{i} i\, X_{i,j} \right).$$

Here $c_1$ is a conversion coefficient with $c_1 = 100/c_e$ with $c_e$ accounting for the FRA detection efficiency in experiments, which is fitted with the kinetic model parameters. Eqs. (S32) and (S33) show that given a value of $C_{min}$, for an exact numerical solution of CAGP we only need to solve the ODEs for the 0th and 1st moment and the ODEs for the complex up to a size of $C_{min} - 1$. If the explicitly modeled maximal complex size is $N_{\max}$, then $C_{min}$ can vary between $n_c$ and $N_{\max} + 1$.

The fitting procedure is schematically explained in **Fig. S14d**. Denote by $CAG_{exp}(t,c)$ and $\sigma_{\exp(t,c)}$ the RI and standard deviation obtained with FRA experiments at time point *t* and for condition *c,* respectively. The data was normalized to the mean maximal value of the DMSO control between 0 and 8h after PP addition. For the data with 20 µM GST-Ex1Q49 (**Fig. 5c**) we used the corresponding DMSO control at 2 µM GST-Ex1Q49 for normalization as for this experiment the amount of protein blotted was nominally the same (500 ng). The data is then converted to relative protein amounts $CAGP_{exp}(t,c)$ using the inverse of Eq. (S30) and then a background value, estimated from the first 2 time points, is subtracted. Given a kinetic parameter set $\psi$ we first numerically integrate the model for the different experimental conditions. In a first step of the fitting the best value of $C_{min}$ given the kinetic parameters is computed. For this *CAGP* curves are computed for all possible values of $C_{min}$ (Eq. (S32) or (S33)) and optimal values of $c_1$ are obtained by linear regression. We add back to *CAGP* the background values estimated experimentally and convert to relative immunoreactivities using Eq. (S30) yielding simulated relative immunoreactivities $CAG_{mod}(t,c,C_{min})$ for each condition *c* and each possible value of $C_{min}$. We then computed the weighted squared distance

$$\chi^2(\psi, C_{min}) = \sum_c \sum_t \left(\frac{CAG_{exp}(t,c) - CAG_{mod}(t,c,C_{min})}{\sigma_{\exp(t,c)}}\right)^2 \qquad \text{(S34)}$$

and selected the value of $C_{min}$ that gave the lowest $\chi^2$. This gave $\chi^2(\psi)$ characterizing the parameter set $\psi$.

For **Fig. S16** we globally fitted the kinetic parameters, $c_1$ and $C_{min}$ to the FRA time course data for the 5 experimental conditions defined by the 5 different initial GST-Ex1Q49 concentrations (symbols and error bars in **Fig. S16a and c**) using the same parameter set for all conditions. For **Figs 5c-d** and **S15**, we modeled 5 different experimental conditions: Ex1Q49 aggregation at 2 and 20 µM initial GST-Ex1Q49 concentrations without O4 and at 2, 4 and 20 µM initial GST-Ex1Q49 concentrations with O4 (symbols and error bars **Fig. 5c** and **Fig. S15**). In addition to the kinetic parameters and the CAG53b detection parameters $c_1$ and $C_{min}$ we also fitted the 2 parameters characterizing the action of O4 Eqs. (S27)-(S29).

For each model we solved numerically the equations for the moments and for complexes containing up to 40 molecules (ode15s, MATLAB7.1, The MathWorks Inc., Natick, MA, 2000). The range of parameter values allowed in the model is given in **Table S4**. Parameter optimization was performed in two steps: (*i*) starting from a random parameter set we performed a simulated annealing with a non-zero asymptotical temperature (see **Fig. S14d**) followed by (*ii*) a non-linear least-square solver to refine the parameters (lsqnonlin, MATLAB7.1, The MathWorks Inc., Natick, MA, 2000). This procedure was repeated 30 times and repeatedly led to minima with a $\chi^2$ within 1-3% of the best possible $\chi^2$. Finally, a Markov-Chain-Monte-Carlo procedure was used to estimate the confidence interval of the parameters using the best $\psi_{opt}$ as starting value. For all the simulations we obtained $C_{min}$ values (Eqs. (S32)-(S33)) far below $N_{\max}$= 40 (**Table S5**) indicating that, given the data, a model that explicitly simulates complexes up to a size of 40 provides correct numerical solutions. The parameters from the best fitting model with nucleated branching used in **Fig. 5c,d**, **6a**, **7a,b** with $n_c = n_b = 1$, are given in **Table S5**.

For model comparison we used the corrected Akaike information criterion (AICc) [29]. This reads for a model *k* with $N_k$ number of parameters

$$AICc(k) = N\log\left(\frac{\chi^2}{N}\right) + 2N_k + 2\frac{(N_k+1)N_k}{N-N_k-1}, \tag{S35}$$

where *N*=52 is the number of data points. **Fig. 5d** and **S16b** show the differences to the AICc of the best fitting models for the respective data set.

### Legends of Supplemental Figures S1 to S19

**Supplemental Figure S1 related to Figure 1. Characterization of GST-Ex1Q49 fusion protein.**

(a) Purified GST-Ex1Q49 and GST proteins were analysed on a 4-12% SDS-gradient gel. Coomassie staining of recombinant proteins (left panel); Western blots with anti-GST (middle panel) and anti-HTT antibodies (HD1 right panel) were performed. GST-Ex1Q49 migrates at ~60 kDa; the protein migrates at a higher molecular weight than expected from the amino acid sequence. The anti-HTT antibody HD1 recognized GST-Ex1Q49 but not GST alone. The anti-GST antibody recognized both GST and the GST-Ex1Q49 fusion protein.

(b) Size exclusion chromatography of GST-Ex1Q49 fusion protein. Absorbance was recorded at 280 nm; the predominant species eluted at a molecular size of ~510 kDa. Size markers are indicated in kDa. Fractions were dot blotted and membranes were developed with the HD1 antibody. Fractions 6-12 exhibited the highest HTTex1 immunoreactivity (Pool 1). Pool 1 was concentrated and analysed by SDS-PAGE and Western blotting using HD1 antibody. Oligomeric GST-Ex1Q49 fusion protein disassembles under denaturing conditions into monomers migrating at ~60 kDa.

(c) Amino acid sequence of the Ex1Q49 fragment released by PP cleavage from GST-Ex1Q49 fusion protein.

(d) Representative picture of a peptide array of the HTTex1 protein probed with CAG53b antibody and detected with HRP fused secondary antibody. The 15 amino acids per peptide spot are displayed to the right of each dot. Intensity indicates antibody-antigen recognition. The CAG53b antibody detects amino acids within the first 17 residues and in the proline-rich regions following the polyQ.

(e) Cleavage of the GST-Ex1Q49 fusion protein with PreScission protease (PP). GST-Ex1Q49 was incubated at a concentration of 2 µM in the presence of 0.28 U PP per µg fusion protein. Cleavage was performed at 20°C and shaking at 300 rpm. Aliquots were taken from the reaction at the indicated time-points; samples were analysed by SDS-PAGE and Western blotting using the HD1 antibody. More than 90% of the GST-Ex1Q49 fusion protein was cleaved after 3 h. Quantification of HD1 immunoreactivity was performed by densitometry using the AIDA software. All signals were normalized to $t$ = 0 h.

**Supplemental Figure S2 related to Figure 1. Investigation of the impact agitation speed and potential modulator proteins on the formation of Ex1Q49 aggregates.**

(a) Spontaneous formation of Ex1Q49 aggregates. GST-Ex1Q49 fusion protein (2 µM) was incubated with PP at three different agitation speeds (0, 300, 600 rpm). Aliquots taken at the indicated time points were analysed by FRAs, and aggregates were immunodetected with CAG53b antibody. Signals were normalized to the 7 h signal at 300 rpm.

(b) The addition of BSA protein (0.5, 2, and 5 µM) to standard reactions (2 µM GST-Ex1Q49 + PP) does not influence spontaneous Ex1Q49 aggregation. Aliquots were taken at the indicated time-points and analysed by FRAs. Retarded aggregates were immunodetected with HD1 antibody and quantified densitometrically using the AIDA software. All signals were normalized to the 8 h signal in the absence of additional BSA. Values are means of triplicates ± standard deviation.

**Supplemental Figure S3 related to Figure 1. Investigating spontaneous Ex1Q49 aggregation by AFM and TEM.**

(a) Representative AFM picture of spontaneously formed Ex1Q49 fibril bundles. Standard aggregation reactions were analyzed, where GST-Ex1Q49 fusion protein (2 μM) was incubated with PP at 20°C. The height, width and length of fibril bundles were determined using the JPK data processing software. The arrows indicate the scanned regions in fibril bundles (green: length and height, blue: width and height).

(b) Time-dependent quantification of the height, width and length of fibrillar Ex1Q49 bundles by AFM. Standard aggregation reactions were analyzed as in (*a*). In spontaneous Ex1Q49 aggregation reactions large, highly complex fibrillar structures are formed that grow in size over time. Bars show mean and standard deviation over $n$ = 10 bundles (for $t$ = 3 h and 6 h) and $n$ = 6 bundles (for $t$ = 4 h).

(c) Systematic analysis of aggregation reactions with GST-Ex1Q49 (2 μM) and PP by transmission electron microscopy (TEM) show multiple branching points in spontaneously formed Ex1Q49 fibrils (red arrows). Samples were analysed after an incubation period of 8 h.

**Supplemental Figure S4 related to Figure 2. Investigation of spontaneous Ex1Q49 aggregation with dot blot assays using epitope-specific antibodies.**

(a) Quantification of Ex1Q49 protein on Western blots using the antibodies HD1 and MW1. With both antibodies similar curves were obtained, indicating that proteolytic cleavage of GST-Ex1Q49 protein with PP leads to a time-dependent decrease of MW1 antibody binding.

(b) Insoluble Ex1Q49 aggregates are not recognized by 3B5H10. Fibrillar Ex1Q49 aggregates were produced by incubating 2 µM GST-Ex1Q49 fusion protein for 24 h with PP. Aggregates were washed to remove potential soluble protein; samples were analyzed by dot blot assays using the antibodies 3B5H10 and HD1, respectively. The insoluble Ex1Q49 aggregates were recognized by the polyclonal antibody HD1, which recognizes a C-terminal proline-rich region, but not by the anti-polyQ antibody 3B5H10.

(c) Comparison of the Ex1Q49 aggregation curves obtained by filter retardation assays (FRAs) and dot blot assays (DBAs). The epitope-specific antibody MW8 detects insoluble Ex1Q49 aggregates in non-denaturating DBAs as well as denaturating FRAs. The polyclonal antibody CAG53b was used as a control.

**Supplemental Figure S5 related to Figure 3. Effects of small molecules on Ex1Q49 aggregation.**

(a-c) Aggregation of Ex1Q49 was not altered by addition of the chemical compounds curcumin (Curc), methylen blue (MB) and PGL034. Aggregation reactions of 2 µM GST-Ex1Q49 were incubated in the absence or presence of compounds. Aliquots taken at the indicated time-points and were analysed by FRA. The aggregates were immunodetected with CAG53b antibody and quantified by densitometry using the AIDA software. The signals were normalized to the 24 h signal in the absence of compound. The compounds (a) Curc, (b) MB and (c) PGL034 had no effect on the formation of SDS-resistant aggregates.

(d) O4 has no effect on PP cleavage. Aggregation of Ex1Q49 (2 µM) was examined in the presence or absence of equimolar concentrations of O4. Cleavage of GST-Ex1Q49 fusion protein was analysed by SDS-PAGE and immunoblotting using the anti-HTT antibody HD1. Bands were quantified by densitometry and signals were normalized to $t = 0$ h.

(e) Quantitative analysis of FRA results shown in Fig. 3b. The highest signal obtained with control aggregation reactions in the absence of O4 was set to 100%. The lag phase duration (lag-time, green) and growth rate given as the maximal slope (blue) can be measured. Quantification of these two parameters for the FRA traces gave a lag-time of 1.86 ± 0.62 h in controls compared to 4.22 ± 0.79 h in O4 treated samples (2 µM O4, n = 13, paired *t*-test, *p*-value = 5e-10). The growth rate, was not influenced by compound treatment (control 38.85 ± 9.67 h-1, growth rate 2 µM O4 45.08 ± 9.41 h-1, *n* = 13, paired *t*-test, *p*-value = 0.096).

(f) Dynamic light scattering analysis of spontaneous Ex1Q49 aggregation (20.9 µM) in the presence (20.9 µM) and absence of O4 (*n* = 4). The z-averaged diameter was computed according to Eq. S4.

(g) GST-Ex1Q49 fusion protein (2 µM) was digested with PP to cleave off the GST tag and incubated with DMSO or O4 at 37°C. After 24h of incubation aliquots were analysed by atomic force (AFM) and transmission electron (TEM) microscopy (right). The picture in the middle is a magnification of the framed area in the left AFM picture. The treatment of aggregation reactions with O4 did not significantly influence the morphology of Ex1Q49 fibril bundles. Scale bar for AFM pictures: 1µm; Scale bar for EM pictures: 200 nm.

**Supplemental Figure S6 related to Figure 4. Snapshots from molecular dynamics simulations.**

Snapshots were taken at 0, 50, 100 and 200 ns of the MD simulations with Ex1Q47 with and without O4 (binding sites A7, D1, A5 and A8). Ex1Q47 is shown coloured by region with the same colour code as in Figure 4. The addition of O4 affects the β-sheet strands that are otherwise conserved during the simulations without O4.

**Supplemental Figure S7 related to Figure 4. Snapshots from molecular dynamics simulations.**

Snapshots were taken at 0, 50, 100 and 200 ns of the MD simulations with Ex1Q46 with and without O4. For each simulation two replicas were performed (a and b). Ex1Q46 is shown coloured by region with the same colour code as in Fig. 4. The addition of O4 affects the β-sheet strands that are otherwise conserved during the simulations without O4.

**Supplemental Figure S8 related to Figure 4. Secondary structure propensity per residue, molecular dynamics simulations.**

Secondary structure propensity per residue for the MD simulations performed for Ex1Q47 with and without O4 (binding sites A7, D1, A5 and A8). The addition of O4 affects the propensity of Ex1Q47 residues to adopt β-sheet conformations.

**Supplemental Figure S9 related to Figure 4. Secondary structure propensity per residue, molecular dynamics simulations.**

Secondary structure propensity per residue for the MD simulations performed for Ex1Q46 with and without O4. For each simulation two replicas were performed (a and b). The addition of O4 affects the propensity of Ex1Q46 residues to adopt β-sheet conformations.

**Supplemental Figure S10 related to Figure 4. Molecular dynamics simulations.**

After destroying the β-strands in Ex1Q47, O4 explores the Ex1Q47 surface. O4 is highlighted in yellow sticks; Ex1Q47 is shown using surface representations. Protein residues are coloured according to the normalized probability to find O4 within 5 Å of each residue during the entire MD simulations. The MD simulations for Ex1Q47 (binding sites A5, A7, A8 and D1) with O4 are shown.

**Supplemental Figure S11 related to Figure 4. Molecular dynamics simulations.**

After destroying the β-strands in Ex1Q46, O4 explores the Ex1Q46 surface. O4 is highlighted in yellow sticks, Ex1Q46 is shown using surface representations. Protein residues are coloured according to the normalized probability to find O4 within 5 Å of each residue during the entire MD simulations. Both replicas of the MD simulations for Ex1Q46 with O4 are shown.

**Supplemental Figure S12 related to Figure 4. Inter-residue contact maps, molecular dynamics simulations.**

Inter-residue contact maps for the MD simulations performed for Ex1Q47 with and without O4 (binding sites A7, D1, A5 and A8). The contact maps indicate fewer interactions between β-strands upon addition of O4.

**Supplemental Figure S13 related to Figure 4. Inter-residue contact maps, molecular dynamics simulations.**

Inter-residue contact maps for the MD simulations performed for Ex1Q46 with and without O4. For each simulation two replicas were performed (a and b). The contact maps indicate fewer interactions between β-strands upon addition of O4.

**Supplemental Figure S14 related to Figure 5. Conversion of immunoreactivities to protein amounts, estimation of the PP cleavage rate, and the parameter fitting procedure.**

(a) Relation between protein amount and immunoreactivity. Calibration curves were obtained by serial dilution of the aggregation reaction mixture at time 0 h (HD1) or 8 h after addition of PP (CAG35b). Western blotting experiments and FRAs were performed as described. Proteins on Western blots were detected with HD1 antibody (squares). Aggregates retained on membrane (FRA) were detected with CAG53b antibody (diamonds). Bands were quantified by densitometry and normalized to a signal of 100% protein, that is 250 ng for HD1, and 500 ng for CAG53b. Lines are the calibration curves Eq. S30 for CAG53b, and Eq. S31 for HD1.

(b) Cleavage of 2 µM (square) and 20 µM (circle) GST-Ex1Q49 fusion protein with PreScission protease (PP) as described. Fusion protein cleavage was analysed by SDS-PAGE and immunoblotting using the HD1 antibody. Relative immunoreactivities were converted into relative protein amounts using the calibration curve shown in panel (a) (Eq. S31). The cleavage shows a typical Michaelis-Menten kinetics (solid and dashed lines, Eq. S5).

(c) Dot blot of the Ex1Q49 aggregation mixture at the indicated time points after addition of PP to GST-Ex1Q49 fusion protein detected with CAG53b. The signal of CAG53b is only proportional to the total amount of Ex1Q49 but not its aggregation state.

(d) Pseudo-code of the fitting procedure of model parameters to the data. $C_{min}$ denotes the minimal size of an aggregate, which can be detected by CAG53b. The intervals of allowed parameter values are given in Supplemental Table S4.

**Supplemental Figure S15 related to Figure 5. The kinetic model without nucleated branching and O4 inhibition of primary nucleation does not reproduce the effect of O4 on the aggregation of Ex1Q49.**

(a) Simulated FRA data from the model with nucleated branching ($n_c = 1$ and $n_b = 1$) without (solid lines) and with O4 (dashed lines), assuming an action of O4 on nucleated branching. The experimental data (symbols, mean ± std) show aggregation for 2 different initial GST-Ex1Q49 concentrations and 3 different O4 concentrations as indicated using the CAG53b antibody.

Experimental data and model were normalized to the maximal value of the control with 2 µM initial GST-Ex1Q49.

(b) Simulated FRA data from the model with nucleated branching ($n_c = 1$ and $n_b = 1$) without (solid lines) and with O4 (dashed lines), assuming an action of O4 on templated polymerization. Experimental data as in (a).

(c) The model without nucleated branching and O4 inhibition of primary nucleation does not reproduce the effect of O4 on the aggregation of Ex1Q49. Experimental data as in (a).

**Supplemental Figure S16 related to Figure 5: Fitting the theoretical models to concentration dependencies is not sufficient to distinguish specific aggregation mechanisms.**

(a) Aggregation of Ex1Q49 at different initial GST-Ex1Q49 concentrations monitored using the FRA assay and the CAG53b antibody (dots). Solid lines show the best fitting results for models without nucleated branching ($k_b = 0$, $n_c = 2$, left panel) or with nucleated branching ($n_c = 2$, $n_b = 1$, right panel). A size threshold for the Ex1Q49 aggregates which can be detected, $C_{min}$, is estimated together with the model parameters (see Supplemental Data).

(b) Differences in the corrected Akaike information criterion (*AICc*, Eq. S35) with respect to the best fitting model for models without nucleated branching (circles) and for models with nucleated branching and different values of the nucleus sizes $n_b$ and $n_c$. A large value indicates a less good fit compared to the best model. Models were fitted to the experimental data from (a) as described in Supplemental Data under different assumptions on the size of the aggregates detected by FRA (see (a) and (c)).

(c) Solid lines show the best fitting results for models without nucleated branching ($k_b = 0$, $n_c = 2$, left panel) or with nucleated branching ($k_b = 0$, $n_c =$

2, $n_b$ = 1, right panel) to the data from (a) (dots) assuming that aggregation-competent aggregates of all sizes equally contribute to the signal (detect size $\geq n_c$).

**Supplemental Figure 17 relating to Figures 5, 6 and 7: Summary of the theoretical modelling approach.**

A global quantitative fit of the models without and with nucleated branching to experimental aggregation data using 2 different initial GST-Ex1Q49 concentrations and 3 different O4 concentrations and the CAG53b antibody suggests that the Ex1Q49 aggregation process comprises a nucleated branching reaction. The model without nucleated branching cannot reproduce these data (**Figs 5c, d** and **S15**). However, whether O4 inhibits primary nucleation, templated polymerization or nucleated branching cannot be elucidated from those experiments. The kinetic parameters for the best fit of the model with nucleated branching are given in the supplemental information, **Table S5**. Fitting to the data from an experiment in which only initial protein concentrations are altered, the models with and without nucleated branching were equally probable (**Fig. S16**) under the realistic assumption that only larger aggregates are detected. However, when assuming that FRAs detect aggregates of all sizes the fit to concentration-dependent data favours the model with nucleated branching (Fig. **S16**).

Qualitatively, the experiment in which O4 was added at different time points validated the nucleated branching model with O4 inhibiting primary nucleation (**Fig. 6**). In addition, the model with nucleated branching correctly predicted the qualitative results on the altered lag phase and the maximal aggregate size when adding seeds in different concentrations (compare **Figs 7a,b** to **7c** and **f**).

**Supplemental Figure S18 related to Figure 7. Establishment of a FRET-based assay to quantify the impact of seeds on the formation of HTTex1 fibrils**

(a) SDS-PAGE analysis of purified recombinant GST-Ex1Q48-CyPet and -YPet fusion proteins. Proteins were stained with the dye Coomassie Blue R.

(b) Schematic model of the spontaneous FRET-inducing Ex1Q48-CyPet/-YPet co-aggregation in cell-free assays. Initially, the N-terminal GST-tag keeps the fusion proteins in a soluble state and prevents spontaneous aggregation. After proteolytic cleavage of the fusion proteins, Ex1Q48-CyPet and -YPet fragments are released and spontaneously co-aggregate over time. Co-aggregation is monitored by quantification of FRET, arising when fluorescent tags come into close proximity in ordered protein aggregates.

(c) Analysis of GST-Ex1Q48-CyPet/-YPet co-aggregation by time-resolved quantification of FRET. Aggregation was initiated by proteolytic cleavage of the GST-tag with PP.

(d) Preformed Ex1Q49 fibrils were visualized by AFM before and after fragmentation through sonication for 60 seconds. Scale bar: 1 μm.

**Supplemental Figure S19 related to Figure 7. A schematic representation of the potential impact of fibril-fibril interactions on the time-dependent formation of HTTex1 aggregates in seeding experiments.**

This scheme illustrates two possible outcomes of templated fibrillogenesis depending on whether a lateral association of fibrils contributes to the formation of aggregates or not. HTTex1 monomers incubated with high concentrations of seeds (0 h) will rapid polymerize into many small fibrils until all monomers are consumed (3h). Assuming that HTTex1 fibrils can laterally associate, a prolonged incubation should lead to the formation of large fibril bundles (50 h). In strong contrast, if the fibrils do not associate their size and morphology should not change under these conditions.

### Legends of Supplemental Movies S1 and S2

#### Supplemental Movie 1

MD simulation in the absence of O4 shows that Ex1Q47 retains the four β strands initially present in the structure.

#### Supplemental Movie 2

MD simulation of Ex1Q47 in the presence of O4 (binding site A7) shows the rapid loss of all four β strands present in the initial structure of Ex1Q47.

### Supplemental Tables S1 to S5 and Legends

#### Supplemental Table S1: Chemical structures of compounds tested.

Chemical structure and names of compounds tested in cell-free aggregation assays.

| Structure | Name | Short name | Literature |
|---|---|---|---|
| 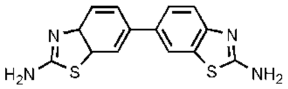 | PGL-034<br>(6-(2-amino-1,3-benzothiazol-6-yl)3H-1lambda-4,3- | PGL034 | Heiser et al., 2002 |
| 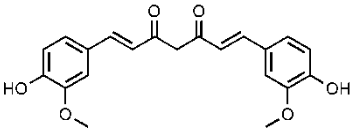 | Curcumin<br>((E,E)-1,7-bis(4-Hydroxy-3-methoxyphenyl)-1,6- | Curc | Dikshit et al., 2006, Yang et al., 2005 |
| 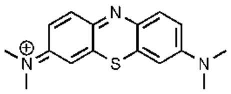 | Methylen blue<br>(3,7-bis(Dimethylamino) phenazathionium chloride) | MB | Oz et al., 2009 |
| 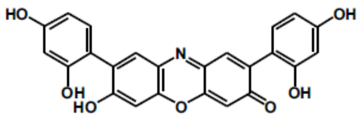 | Orcein-related O4<br>(2,8-bis-(2,4-dihydroxy-phenyl)-7- | O4 | Bieschke et al., 2012 |

**Supplemental Table S2: Computational modeling results: Molecular dynamics and docking.**

Calculated docking affinities and energies of the QM region (O4 molecule) averaged during the last 300 ps of the SCCDFTB-D/CHARMM MD simulations show that O4 is most stable when interacting simultaneously with the β-sheets at the polyQ, N- and C-terminal regions (A7 and D1).

| | Interacting with | Docking affinity (kcal/mol) | SCCDFTB-D/CHARMM MD simulations | |
|---|---|---|---|---|
| | | | Absolute QM energy (kcal/mol) | Relative QM energy (kcal/mol) |
| A1 | polyQ, polyP, N-terminus, C-terminus | -10.4 | -45546.7 ± 4.45 | 23.6 |
| A3 | polyQ | -8.5 | -45554.8 ± 4.85 | 15.5 |
| A5 | polyQ, polyP, N-terminus | -8.2 | -45557.5 ± 4.75 | 12.8 |
| A6 | polyQ | -8.2 | -45547.2 ± 4.37 | 23.1 |
| **A7** | **polyQ, N- and C-terminus** | -8.1 | **-45570.3 ± 5.14** | **0** |
| A8 | polyQ, polyP | -7.8 | -45556.1 ± 5.60 | 14.2 |
| **D1** | **polyQ, N- and C-terminus** | -7.6 | **-45569.3 ± 5.19** | **1.0** |

**Supplemental Table S3: Computational modeling results: Average secondary structure propensity.**

O4 lowers the propensity of Ex1Q47 and Ex1Q46 to adopt β-sheet structures.[a]

| System | α-Helix | Coil | Turn | β-Sheet | Bridge | $3_{10}$-Helix | π-Helix |
|---|---|---|---|---|---|---|---|
| Ex1Q47 | 3 | 53 | 31 | 12 | 1 | 0 | 0 |
| A5 | 0 | 61 | 28 | 9 | 1 | 1 | 0 |
| A7 | 4 | 62 | 28 | 2 | 2 | 2 | 0 |
| A8 | 5 | 55 | 27 | 12 | 1 | 0 | 0 |
| D1 | 4 | 52 | 36 | 7 | 0 | 1 | 0 |
| Ex1Q46 | 0 | 61 | 25 | 10 | 4 | 0 | 0 |
| Ex1Q46 | 0 | 57 | 28 | 11 | 4 | 0 | 0 |
| Ex1Q46-O4 | 0 | 61 | 27 | 7 | 5 | 0 | 0 |
| Ex1Q46-O4 | 0 | 63 | 26 | 7 | 4 | 0 | 0 |

[a] Both replicas of the MD simulations performed with Ex1Q46 are shown.

**Supplemental Table S4: Parameter names and ranges**

Kinetic and O4 inhibition parameters used in the models with primary nucleation, templated polymerization and with or without nucleated branching. The parameter range gives the lowest and highest parameter value allowed in the fitting. The lowest value of $10^{-6}$ has been chosen for numerical reasons and efficient computation. None of the fitted parameters are close to the lowest or highest possible values at the end of the optimization procedure (**Fig. S14d**); r.c. means rate constant. The characteristic time for primary nucleation is given by $1/(k_{n1}+k_{-n1})$. For the fitting procedure, $N_k$=5 parameters ($k_{n1}$, $k_{-n1}$, $k_b$, $k_1$, $C_{min}$) were subject to optimization for the model with nucleated branching, $N_k$=7 if also incorporating an effect of O4. The corresponding models without nucleated branching had one parameter less ($k_b$=0).

| Parameter | Symbol | Parameter range |
|---|---|---|
| **Kinetic parameters** | | |
| Primary nucleation r.c. | $k_{n1}$ | $10^{-6}$ - 1 $h^{-1}\mu M^{-(nc-1)}$ |
| De-nucleation r.c. | $k_{-n1}$ | $10^{-6}$ - 100 $h^{-1}$ |
| Primary nucleus size | $n_c$ | 0 - 4* |
| Nucleated branching r.c. | $k_b$ | $10^{-6}$ - 100 $h^{-1}\mu M^{-nb}$ |
| Nucleated branching order | $n_b$ | 1 - 4 |
| Templated polymerization r.c. | $k_1$ | $10^{-6}$ - 100 $h^{-1}\mu M^{-1}$ |
| **Action of O4** | | |
| Half-maximal inhibition primary nucleation | $K_{I1}$ | $10^{-6}$ - 20 μM |
| Hill-coefficient inhibition primary nucleation | $n_1$ | $10^{-6}$ - 20 |
| Half-maximal inhibition nucleated branching | $K_{I2}$ | $10^{-6}$ - 20 μM |
| Hill-coefficient inhibition nucleated branching | $n_2$ | $10^{-6}$ - 20 |
| Half-maximal inhibition templated polymerization | $K_{I3}$ | $10^{-6}$ - 20 μM |
| Hill-coefficient inhibition templated polymerization | $n_3$ | $10^{-6}$ - 20 |
| Half-maximal of de-nucleation | $K_m$ | $10^{-6}$ - 20 μM |
| Hill-coefficient for de-nucleation | $n_m$ | $10^{-6}$ - 20 |
| Maximal increase of de-nucleation r.c. | $\bar{k}_{-n1}$ | $10^{-6}$ - 2000 $h^{-1}$ |
| **Antibody assays** | | |
| Minimal complex size for detection by FRA | $C_{min}$ | $n_c$ - $N_{max}$+1 |
| Conversion coefficient for FRA | $c_1$ | 0 - 200 |

* a nucleus size of $n_c$ = 0 is spontaneous aggregation without primary nucleation

**Supplemental Table S5: Parameter values for the model with nucleated branching**

The PP parameters are kept fixed and were determined from the data in **Fig. S14b**. The primary nucleus size and nucleated branching order, $n_c$=$n_b$, were not directly fitted but estimated by systematically varying them (**Fig. 5d**). After obtaining a candidate best fit using the pipeline in **Fig. S14d** the most probable parameter distribution given the data is estimated using the Markov-Chain-Monte Carlo method (MCMC). The table gives the median and in squared brackets the lower and upper quartile of the parameter distribution; r.c. means rate constant.

| Parameter | Symbol | Parameter values |
|---|---|---|
| **PP parameters** | | |
| Maximal PP r.c. | $k_r$ | 1.99 $h^{-1}$ |
| Half-maximal PP activity | $K_p$ | 4.211 μM |
| **Kinetic parameters** | | |
| Primary nucleation r.c. | $k_{n1}$ | 0.1225 [0.081- 0.184] $h^{-1}$ |
| De-nucleation r.c. | $k_{-n1}$ | 6.274 [5.235-7.412] $h^{-1}$ |
| Primary nucleus size | $n_c$ | 1 |
| Nucleated branching r.c. | $k_b$ | 0.51 [0.315-0.78] $h^{-1}\mu M^{-1}$ |
| Nucleated branching order | $n_b$ | 1 |
| Templated polymerization r.c. | $k_1$ | 0.4361 [0.275-0.705] $h^{-1}\mu M^{-1}$ |
| **Action of O4** | | |
| Half-maximal inhibition of primary nucleation | $K_{I1}$ | 1.061 [0.921- 1.203] μM |
| Hill-coefficient inhibition of primary nucleation | $n_1$ | 4.711 [4.289-5.223] |
| **Antibody assays** | | |
| Minimal complex size for detection by FRA | $C_{min}$ | 17 [12-22] |
| Conversion coefficient for FRA | $c_1$ | 134.8 [133.24-143.46] |

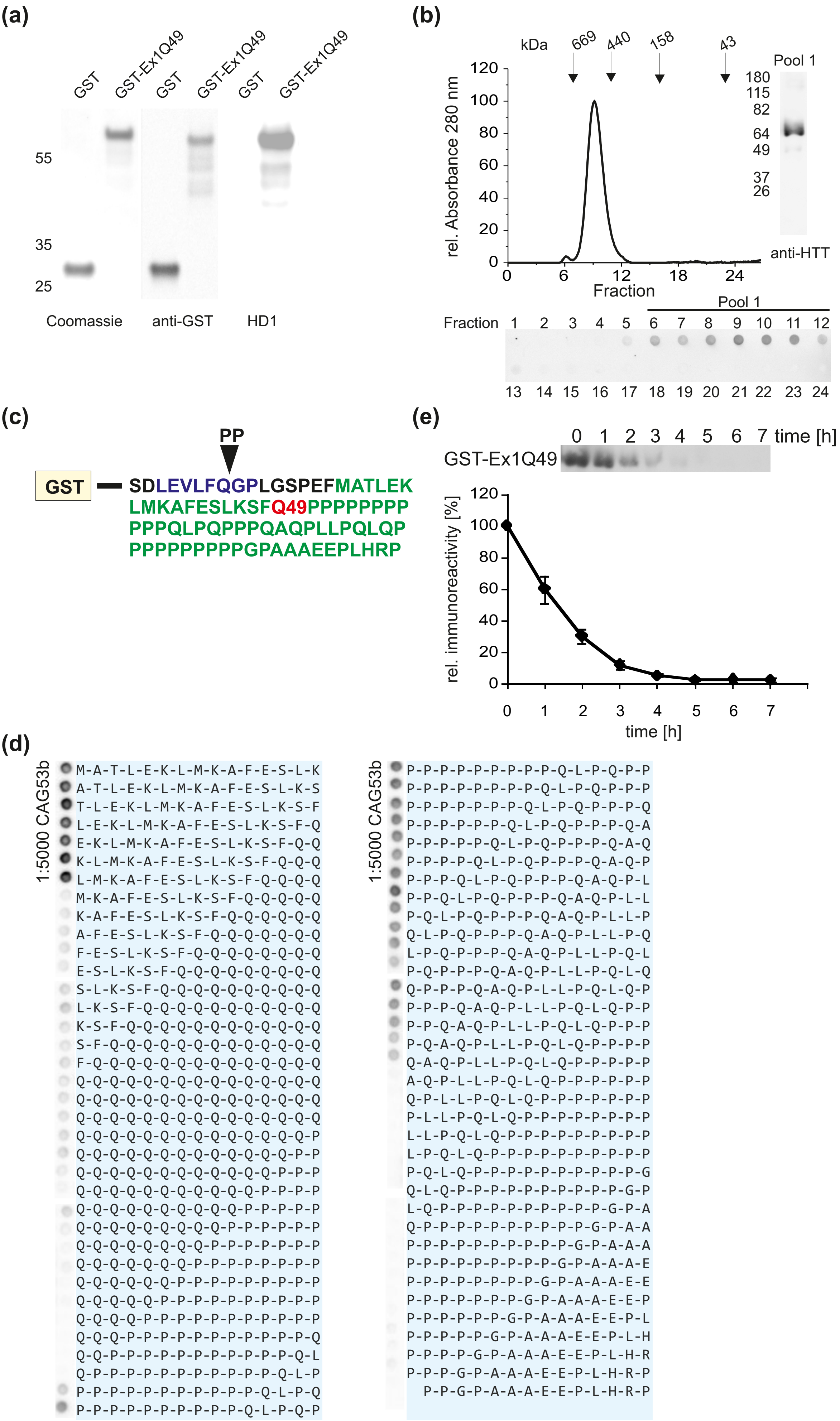

**Suppl. Figure S1**

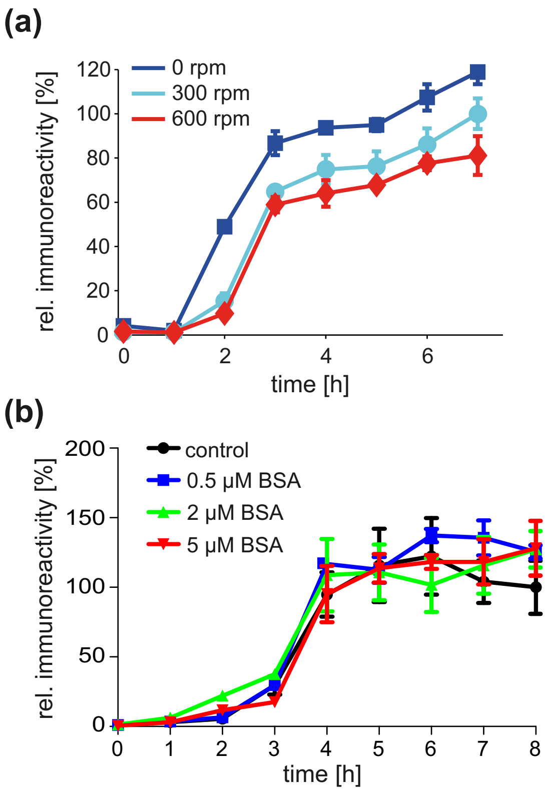

**Suppl. Figure S2**

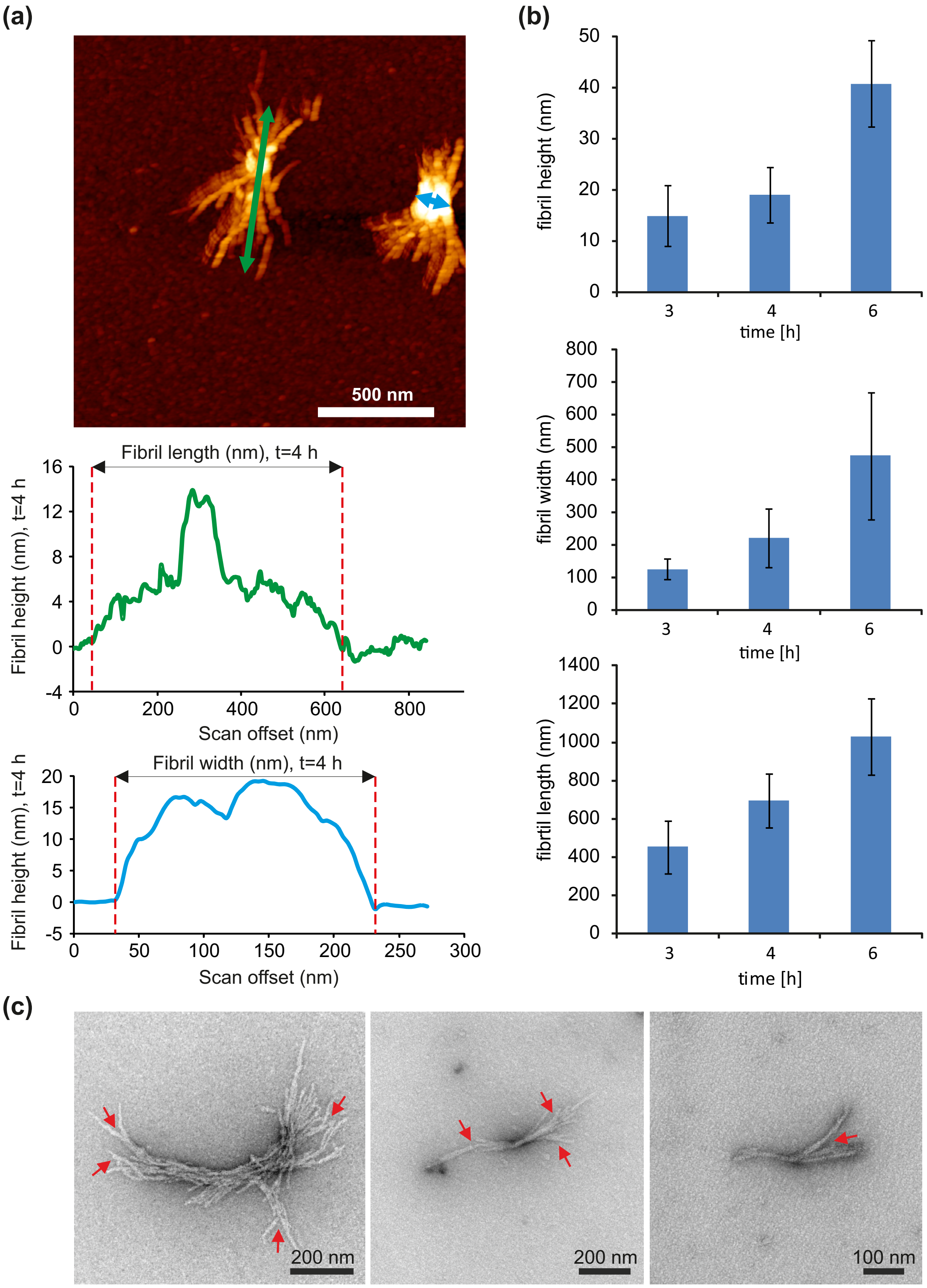

**Suppl. Figure S3**

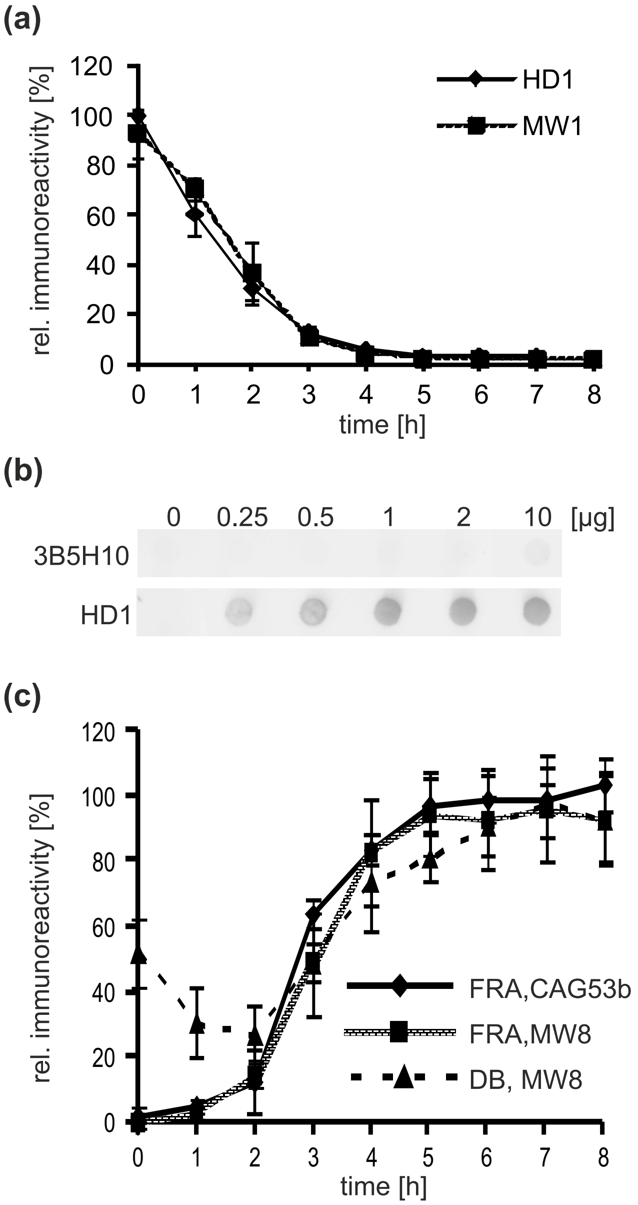

**Suppl. Figure S4**

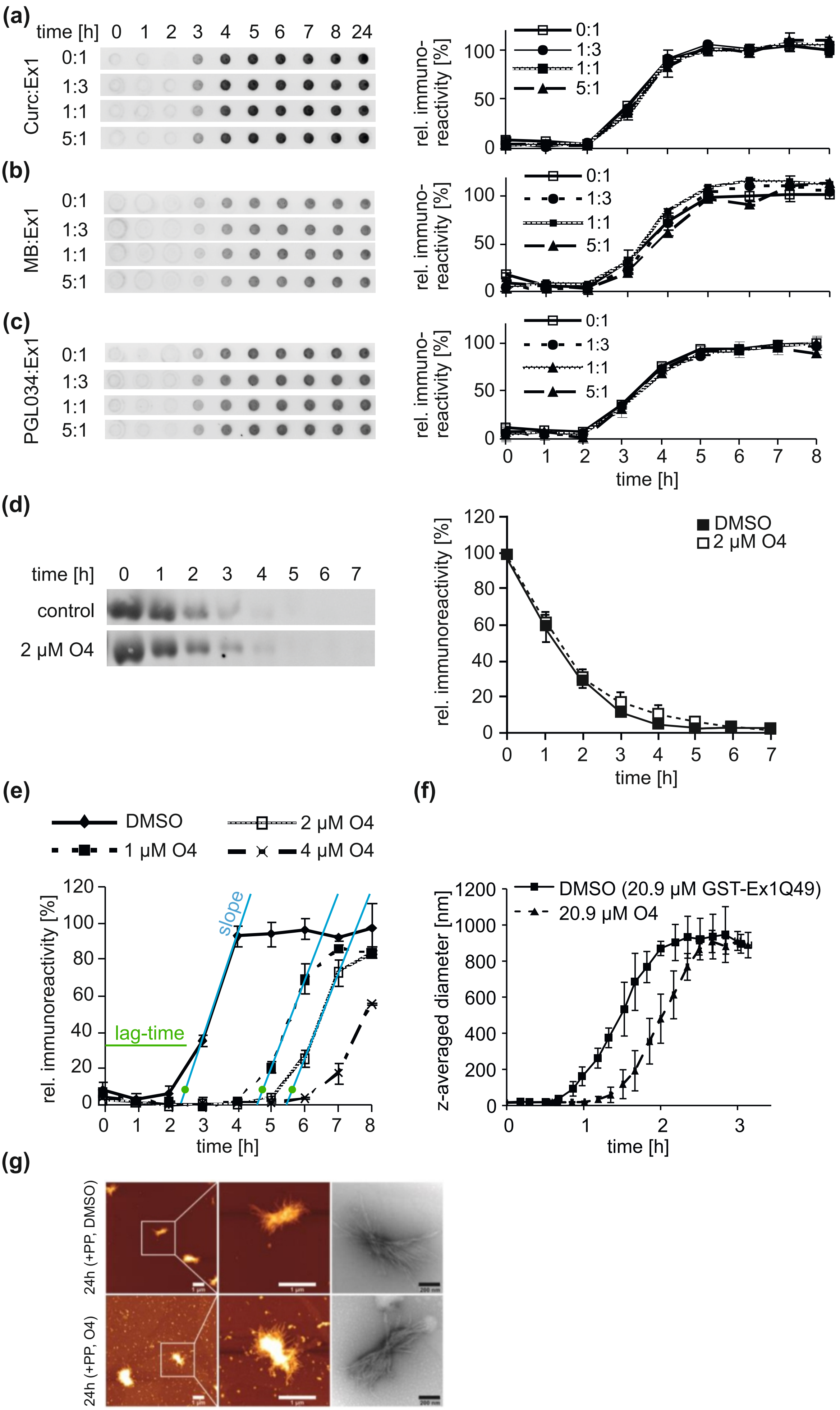

**Suppl. Figure S5**

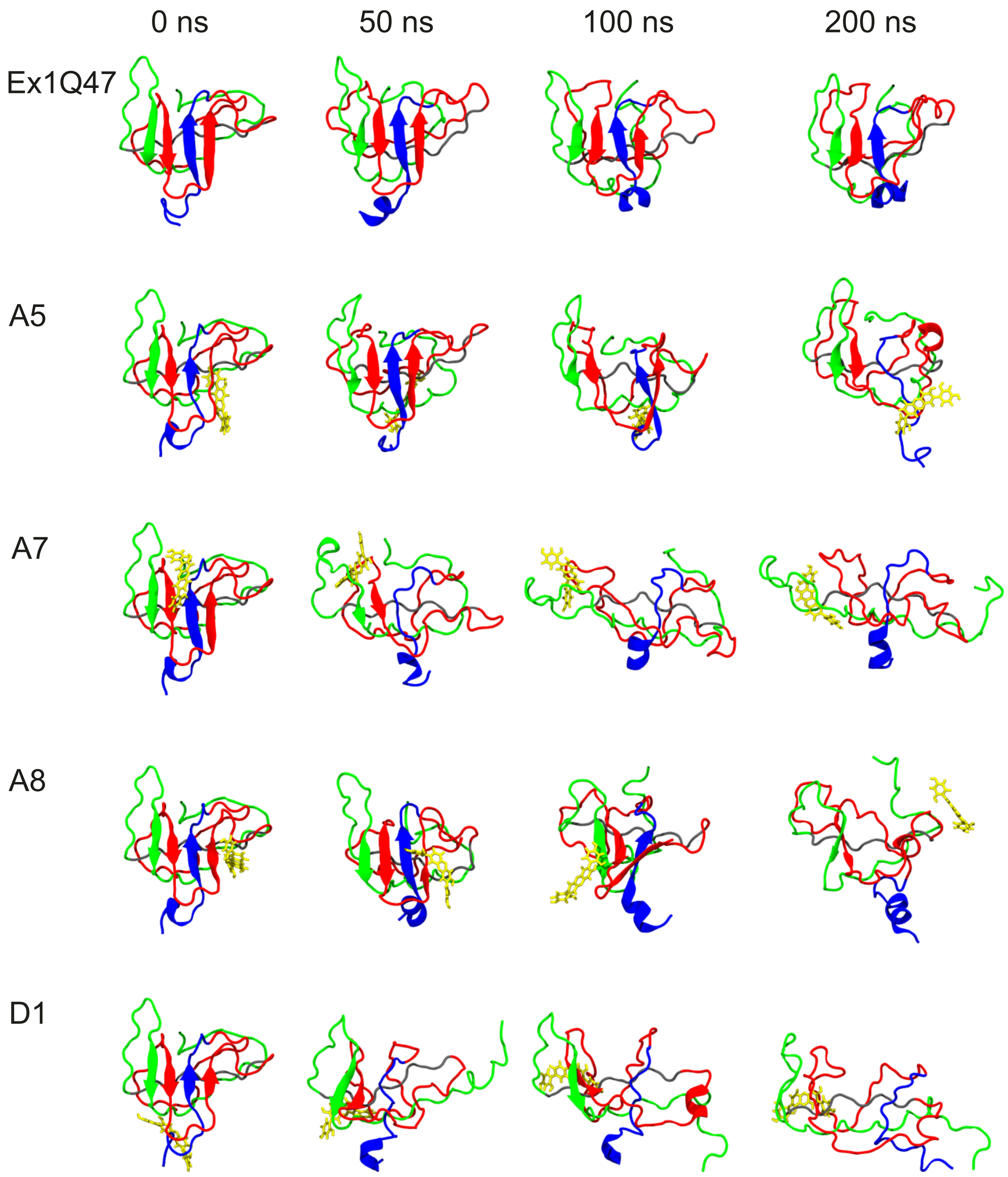

**Suppl. Figure S6**

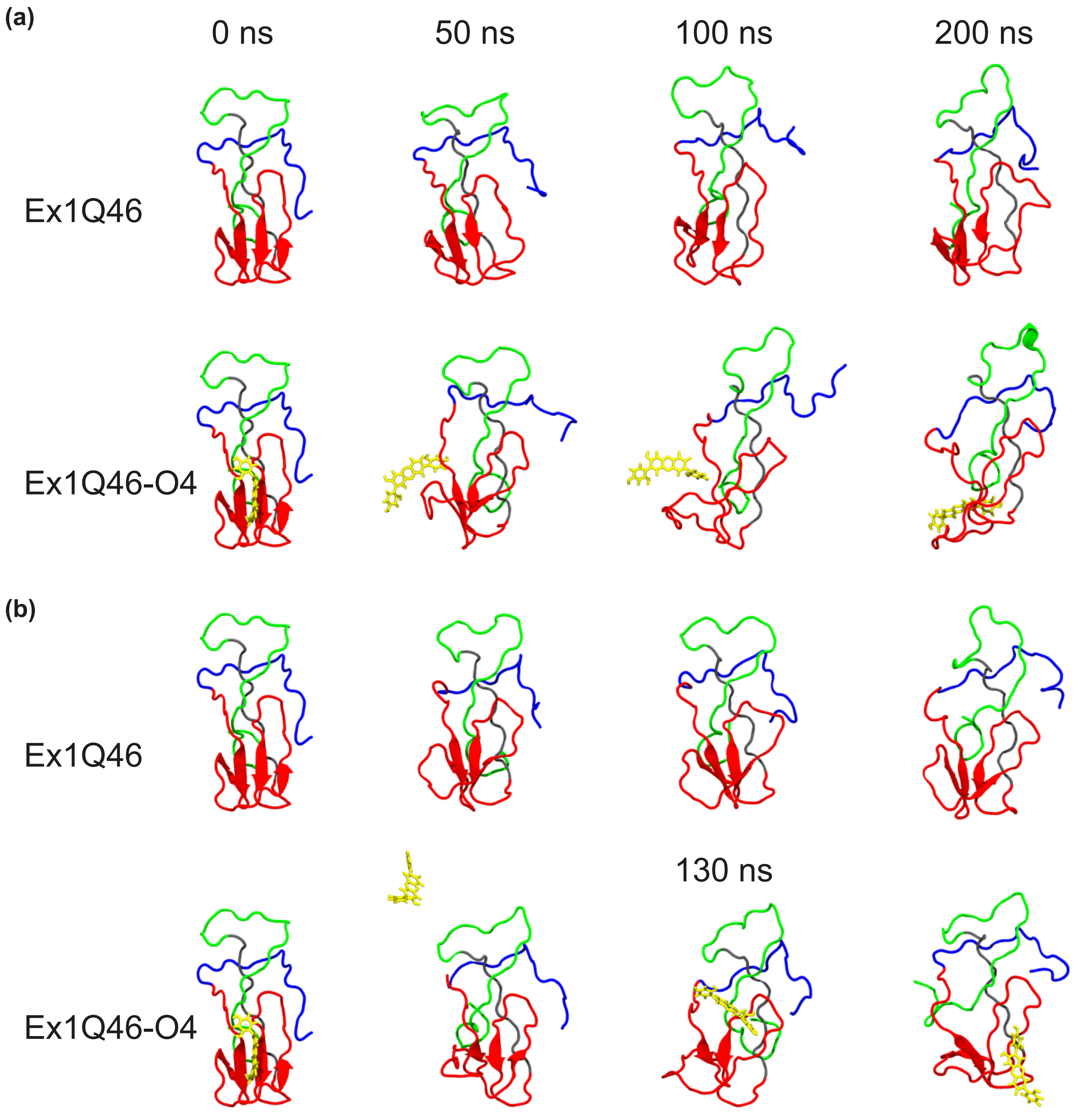

**Suppl. Figure S7**

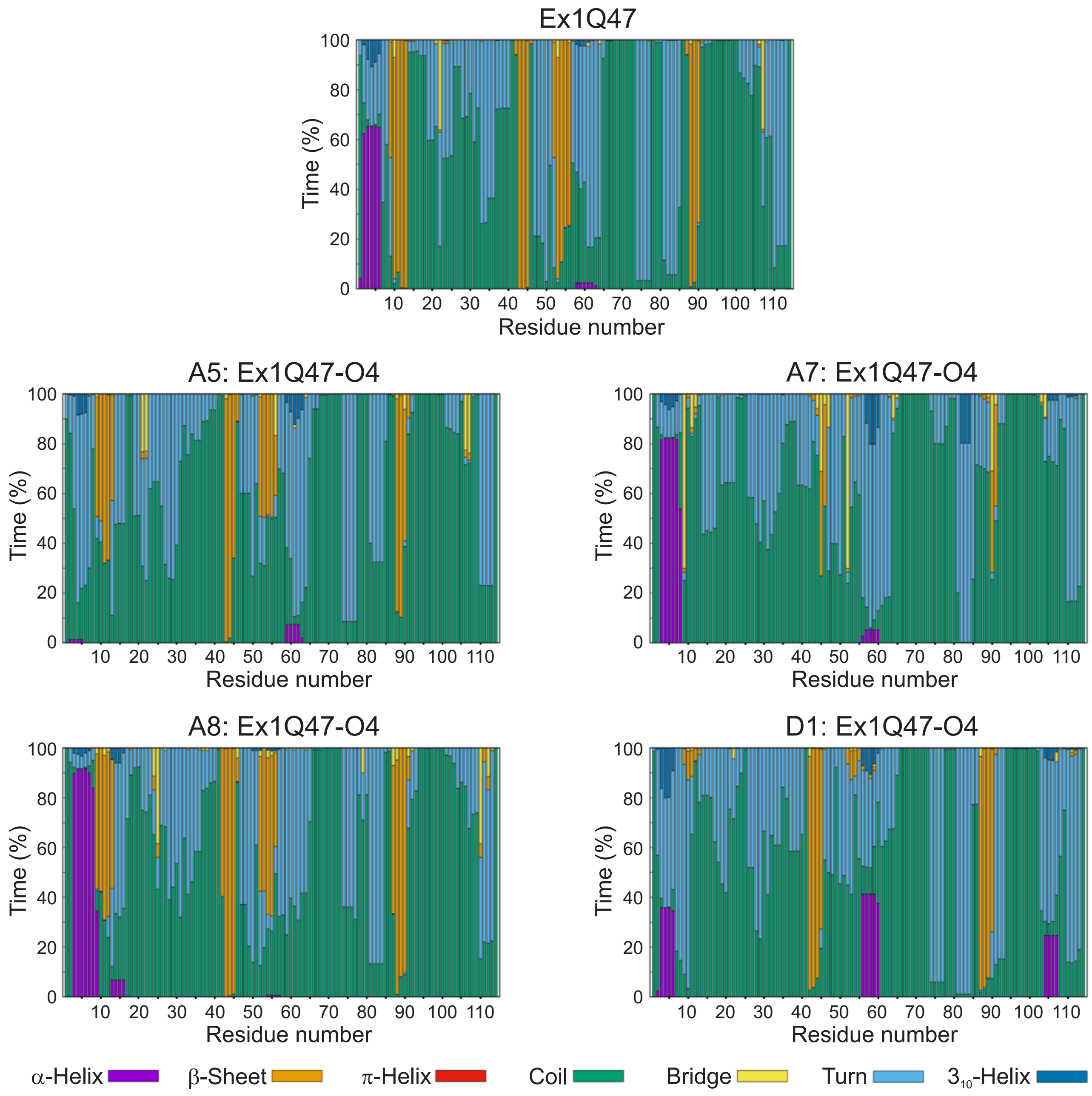

**Suppl. Figure S8**

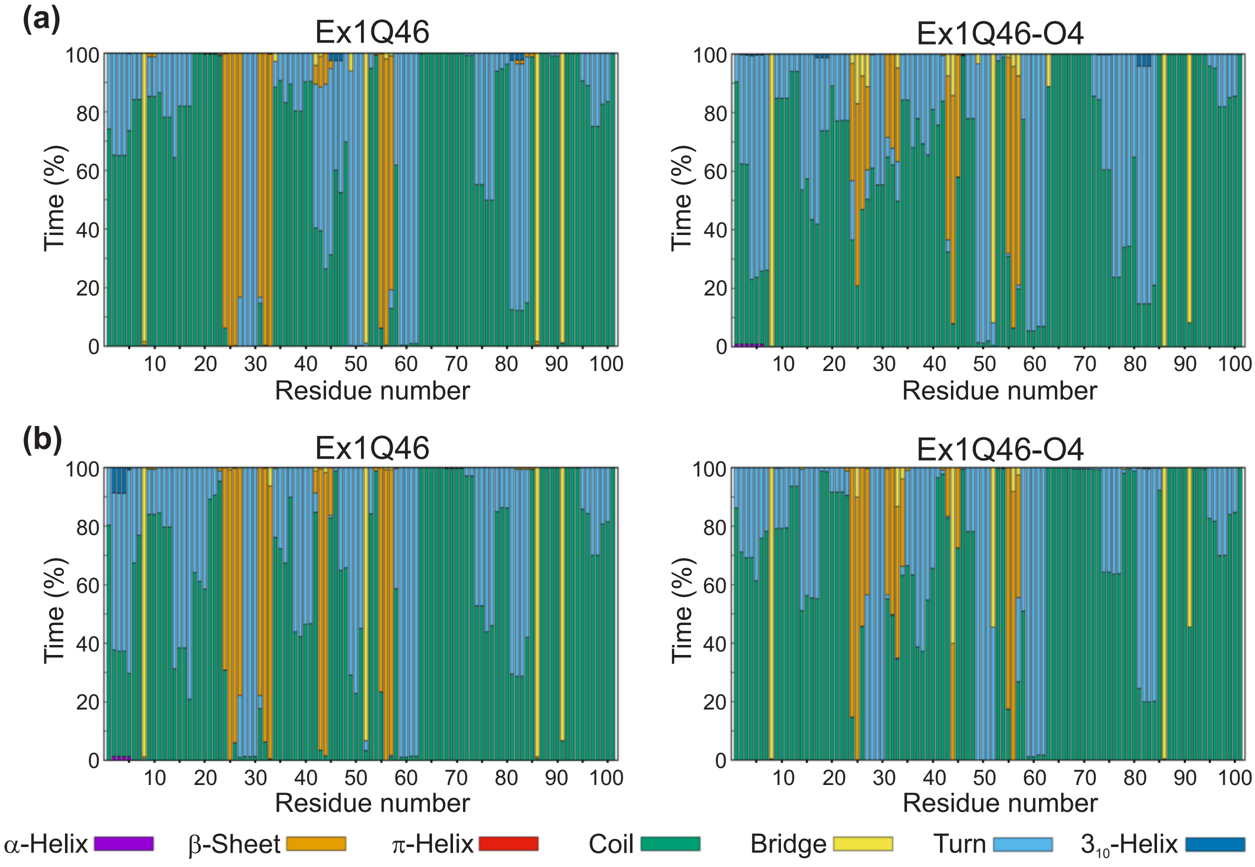

**Suppl. Figure S9**

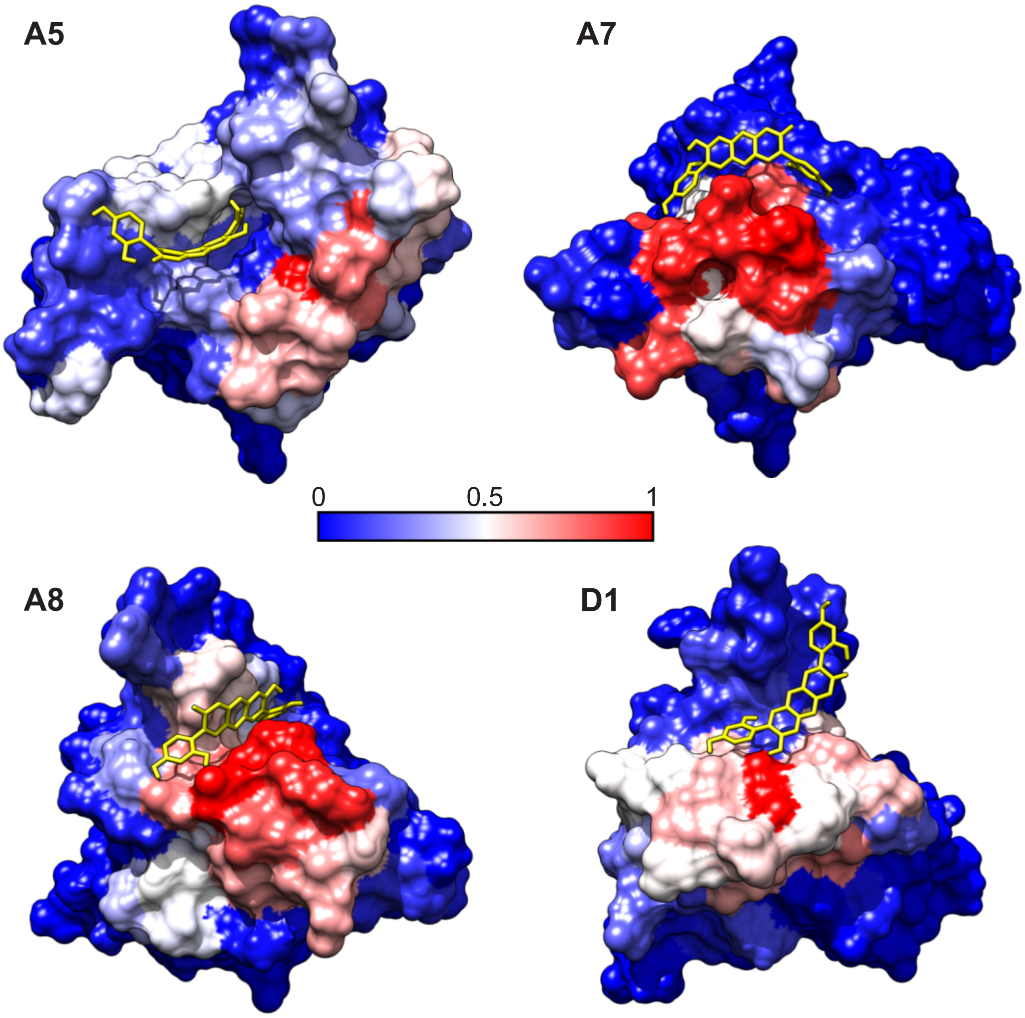

Suppl. Figure S10

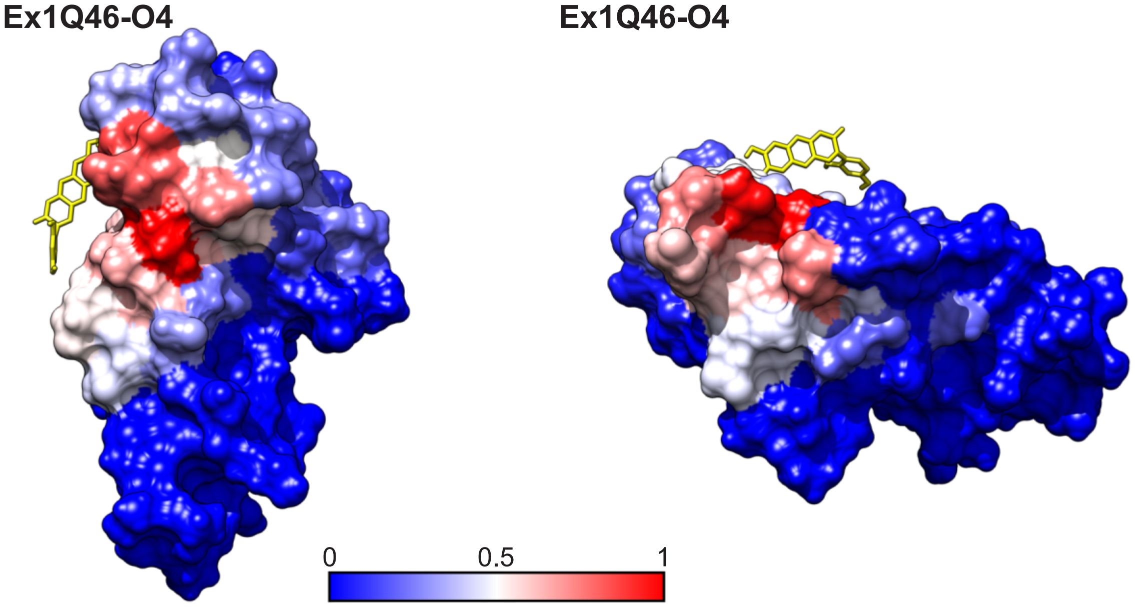

Suppl. Figure S11

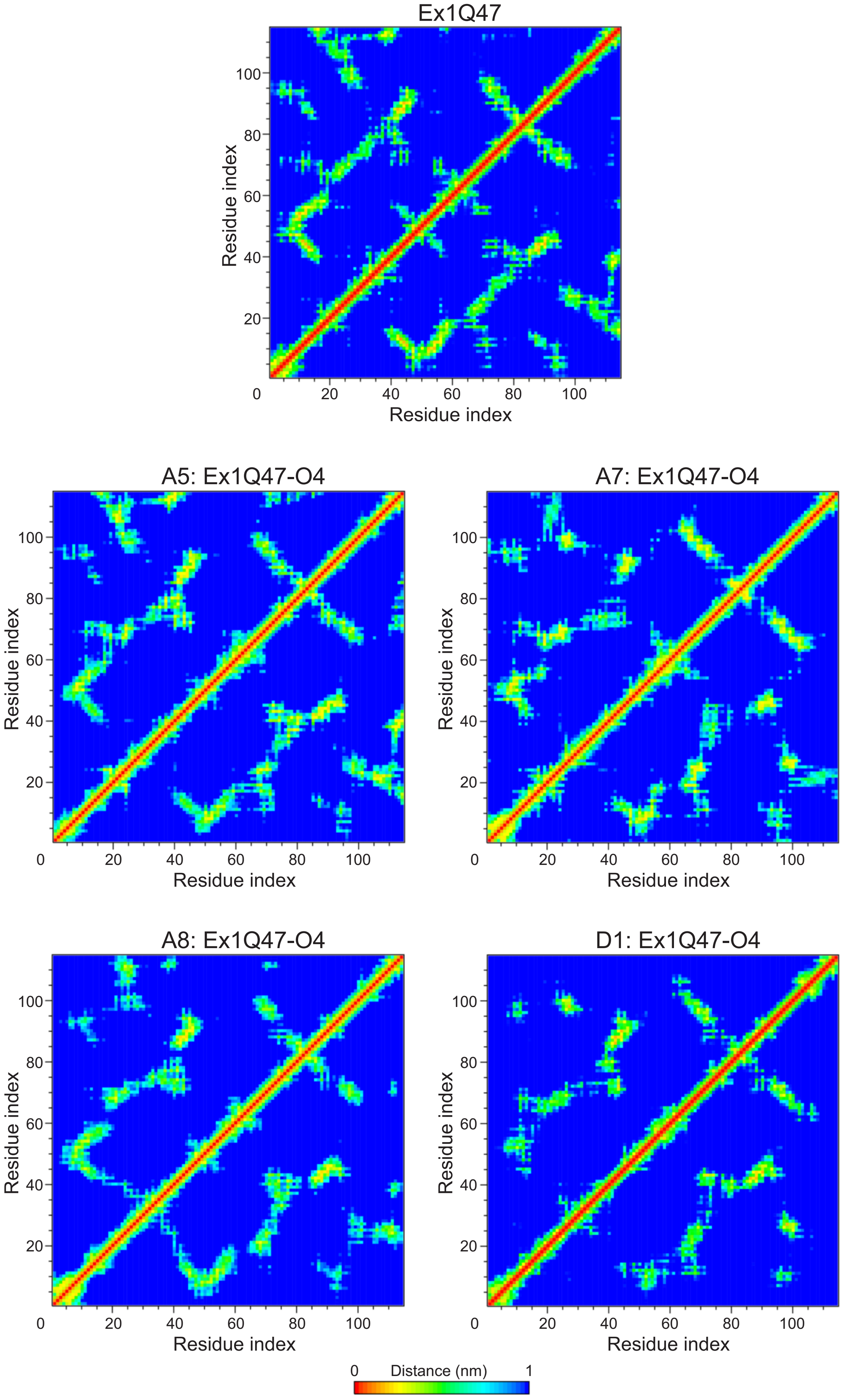

**Suppl. Figure S12**

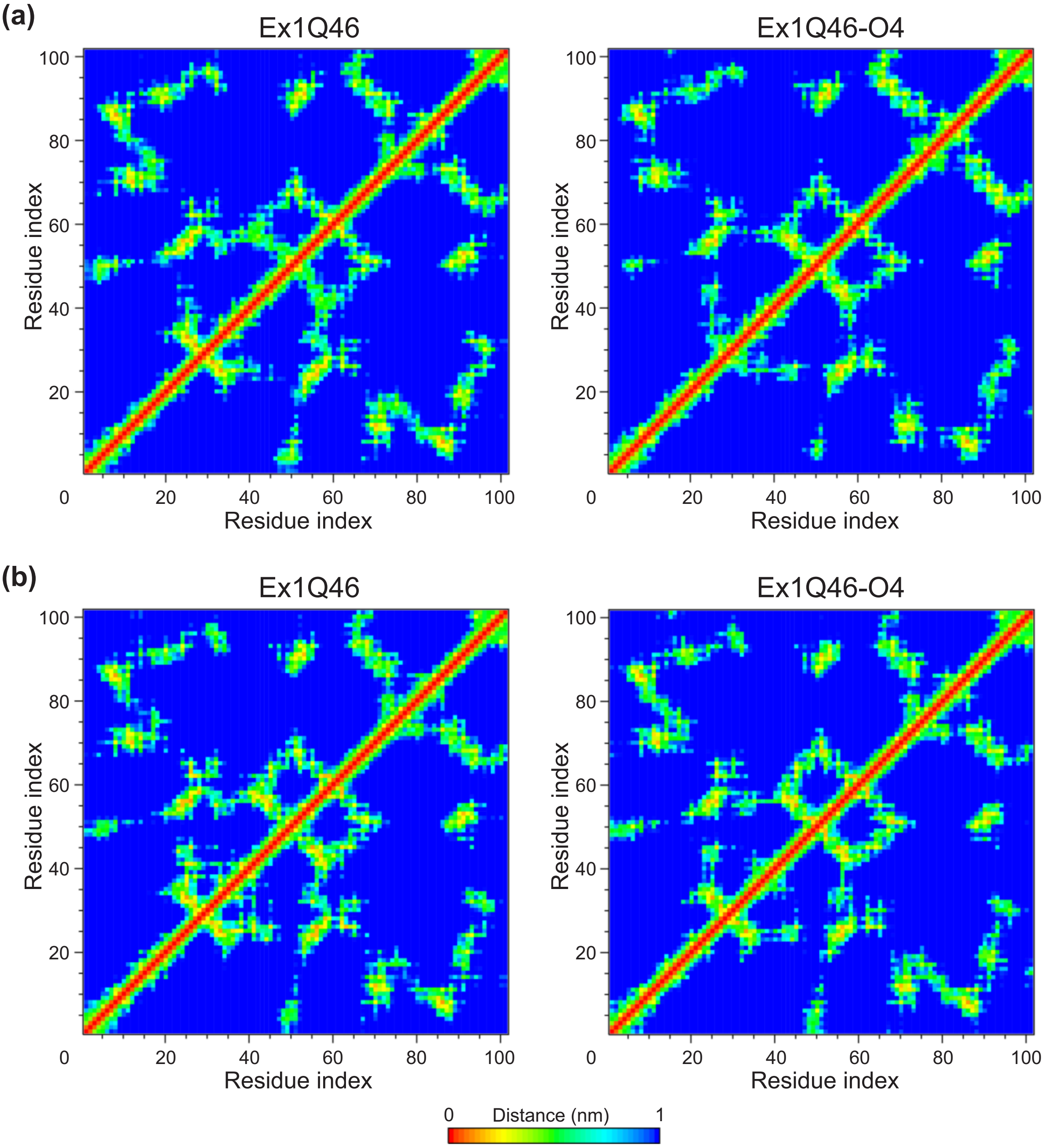

**Suppl. Figure S13**

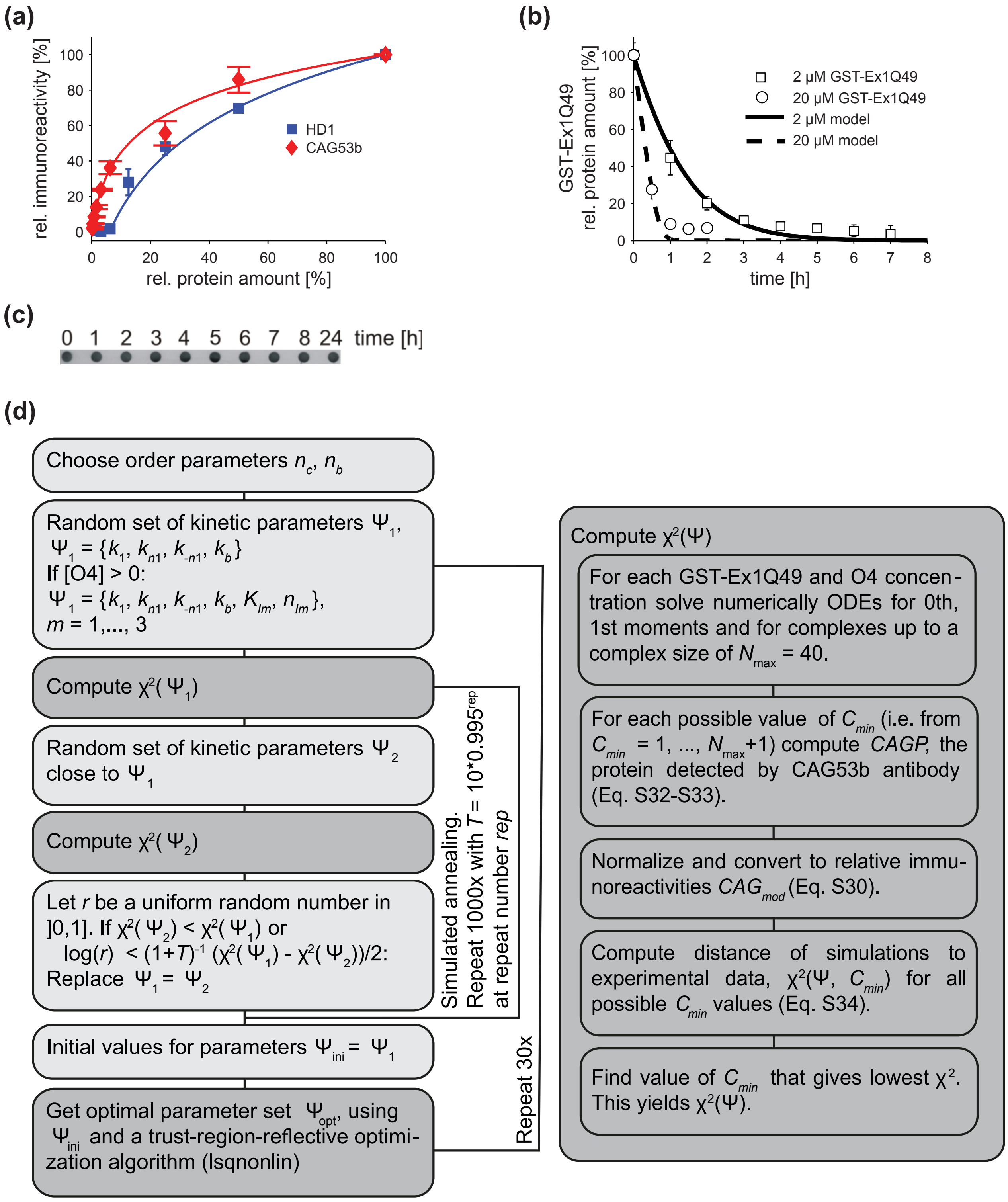

**Suppl. Figure S14**

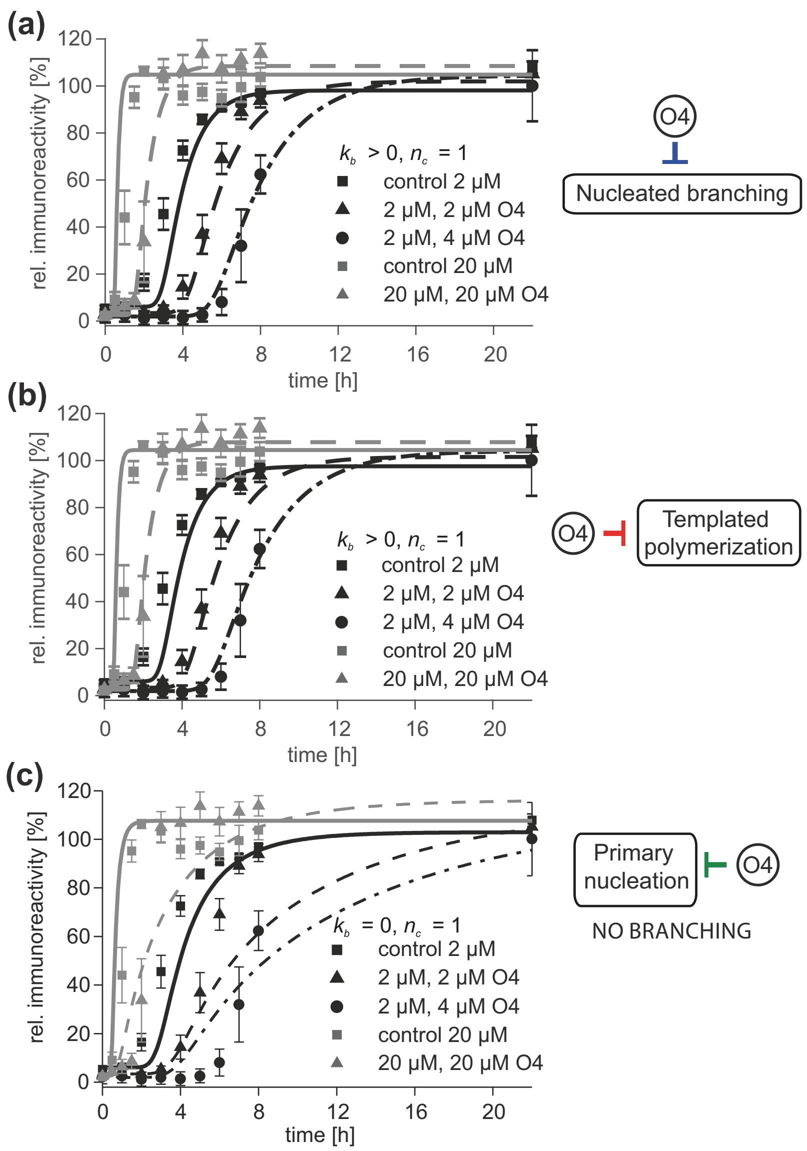

**Suppl. Figure S15**

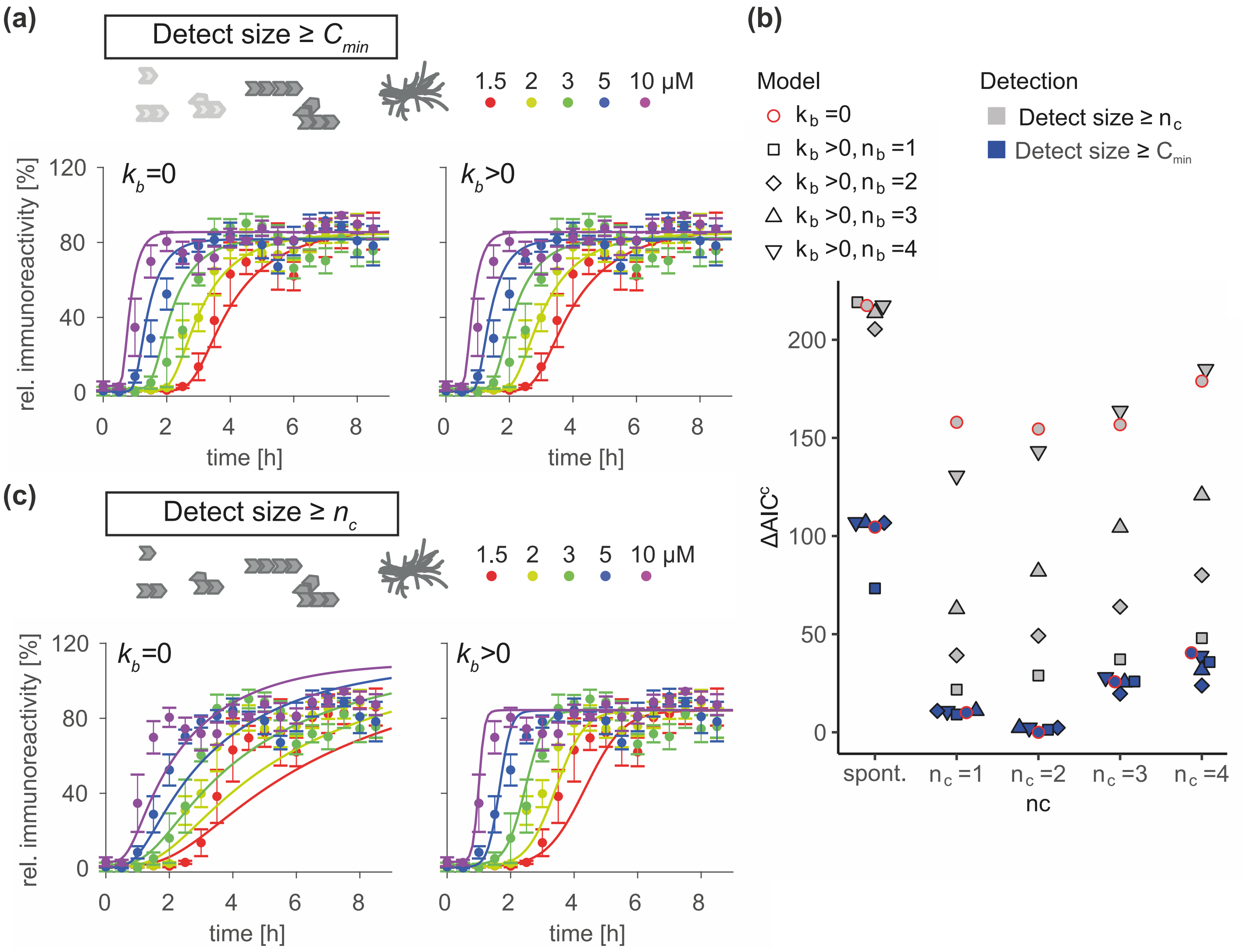

**Suppl. Figure S16**

| Experiment | additional assumption | **Model without nucleated branching**<br>primary nucleation<br>templated polymerization | **Model with nucleated branching**<br>primary nucleation<br>templated polymerization<br>nucleated branching |
|---|---|---|---|
| | | *Global quantitative model fit* | *Global quantitative model fit* |
| adding O4 at t = 0h in different concentrations + different initial concentrations<br>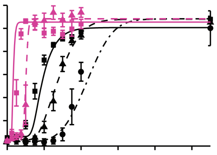<br>Fig. 5 (c) | O4 acts on primary nucleation | ✘ | ✔ |
| | O4 acts on templated polymerization | ✘ | ✔ |
| | O4 acts on nucleated branching | Not applicable | ✔ |
| concentration-dependency<br>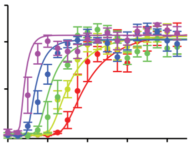<br>Fig. S16 | detection of larger aggregates only* | ✔ | ✔ |
| | detection of aggregates of all sizes | ✘ | ✔ |
| | | | *Qualitative assessment* |
| adding O4 at different time points<br>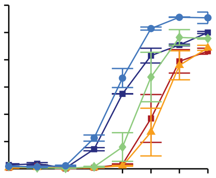<br>Fig. 6 (d) | O4 acts on primary nucleation | Not applicable | ✔ |
| | O4 acts on templated polymerization | Not applicable | ✘ |
| | O4 acts on nucleated branching | Not applicable | ✘ |
| adding different concentrations of seeds<br>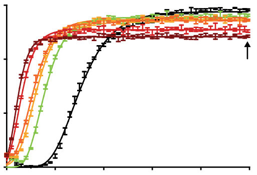<br>Fig. 7 (c) | 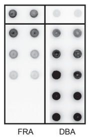<br><br>Fig. 7 (f) | Not applicable | ✔ |

*FRA detects SDS-stable and heat-stable aggregates with sizes >0.2μm, approximately.

**Suppl. Figure S17**

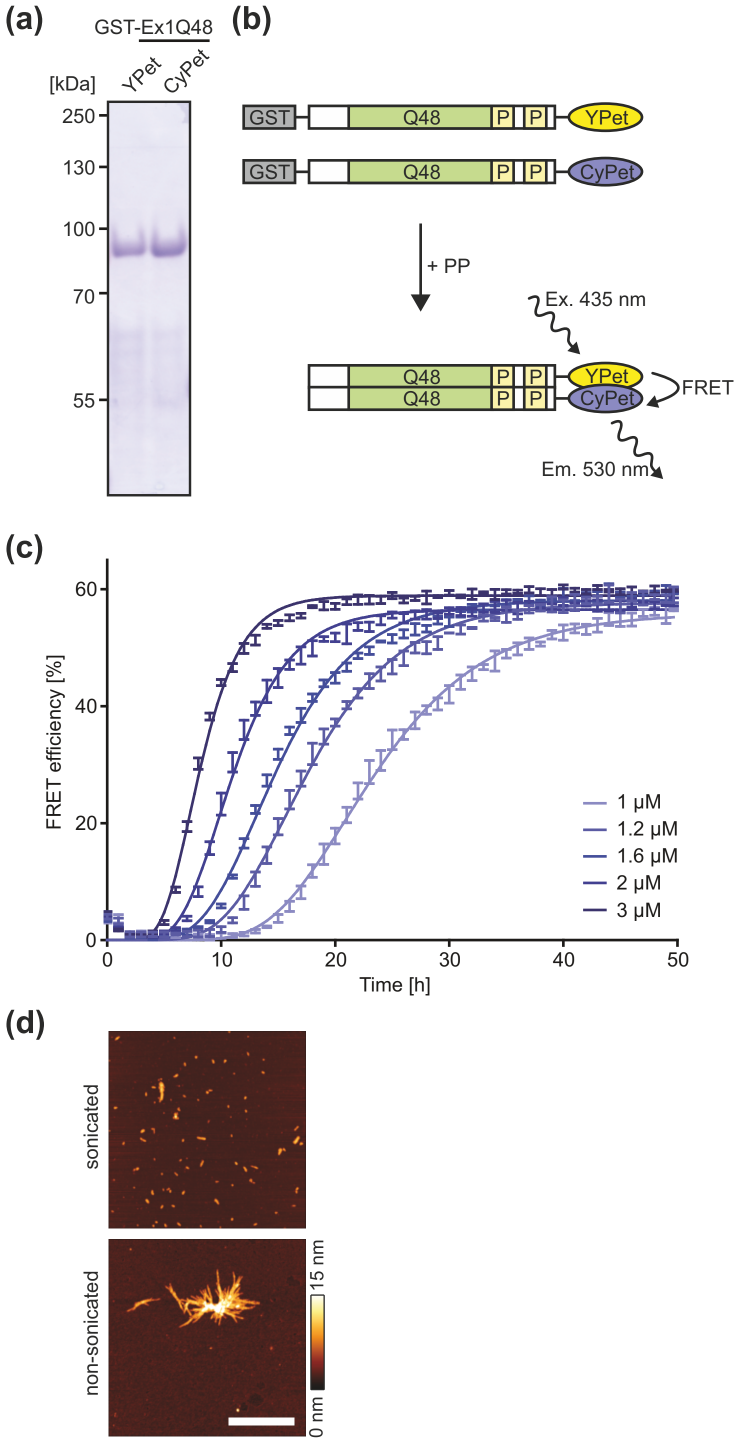

**Suppl. Figure S18**

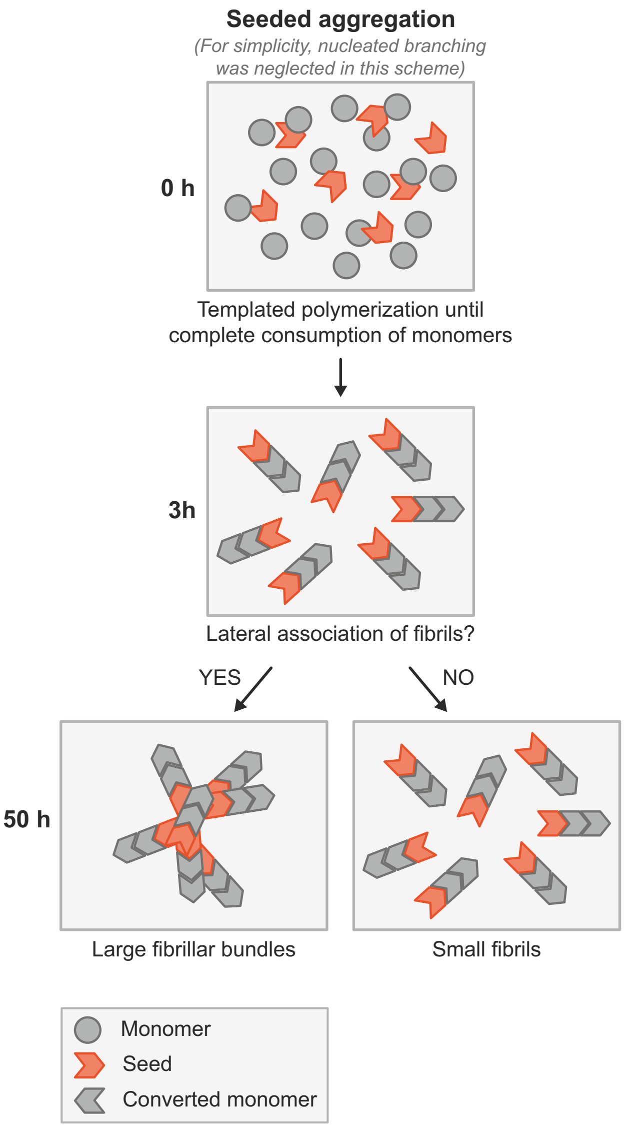

**Suppl. Figure S19**
